# SARS-CoV-2 Requires Cholesterol for Viral Entry and Pathological Syncytia Formation

**DOI:** 10.1101/2020.12.14.422737

**Authors:** David W. Sanders, Chanelle C. Jumper, Paul J. Ackerman, Dan Bracha, Anita Donlic, Hahn Kim, Devin Kenney, Ivan Castello-Serrano, Saori Suzuki, Tomokazu Tamura, Alexander H. Tavares, Mohsan Saeed, Alex S. Holehouse, Alexander Ploss, Ilya Levental, Florian Douam, Robert F. Padera, Bruce D. Levy, Clifford P. Brangwynne

## Abstract

Many enveloped viruses induce multinucleated cells (syncytia), reflective of membrane fusion events caused by the same machinery that underlies viral entry. These syncytia are thought to facilitate replication and evasion of the host immune response. Here, we report that co-culture of human cells expressing the receptor ACE2 with cells expressing SARS-CoV-2 spike, results in synapse-like intercellular contacts that initiate cell-cell fusion, producing syncytia resembling those we identify in lungs of COVID-19 patients. To assess the mechanism of spike/ACE2-driven membrane fusion, we developed a microscopy-based, cell-cell fusion assay to screen ∼6000 drugs and >30 spike variants. Together with cell biological and biophysical approaches, the screen reveals an essential role for membrane cholesterol in spike-mediated fusion, which extends to replication-competent SARS-CoV-2 isolates. Our findings provide a molecular basis for positive outcomes reported in COVID-19 patients taking statins, and suggest new strategies for therapeutics targeting the membrane of SARS-CoV-2 and other fusogenic viruses.

**Highlights:**

- Cell-cell fusion at ACE2-spike clusters cause pathological syncytia in COVID-19
- Drug screen reveals critical role for membrane lipid composition in fusion
- Spike’s unusual membrane-proximal cysteines and aromatics are essential for fusion
- Cholesterol tunes relative infectivity of SARS-CoV-2 viral particles

## Introduction

COVID-19 has caused over a million deaths in the year following identification of its causative pathogen, severe acute respiratory syndrome coronavirus 2 (SARS-CoV-2) (Zhu et al., 2020a). Building on knowledge of similar enveloped coronaviruses (Belouzard et al., 2012; Heald-Sargent and Gallagher, 2012), recent studies made astonishing progress toward a holistic understanding of SARS-CoV-2 pathobiology, suggesting amenable targets to therapeutic intervention (Haynes et al., 2020; Stratton et al., 2020; Tay et al., 2020). In particular, the unprecedented pace of SARS-CoV-2 research led to key insights into viral fusion (V’kovski et al., 2020). Many early studies, including most small molecule and genetic screens, focused on entry (Chen et al., 2020; Dittmar et al., 2020; Riva et al., 2020; Wei et al., 2020; Zhu et al., 2020b). Central to these efforts are trimeric spike glycoproteins (or “peplomers”), which give the viral envelope its crown-like appearance, and ACE2, their essential human receptor (Duan et al., 2020; Mittal et al., 2020). Association of the two proteins underlies virus-cell adhesion, which precedes a conformational change in spike that unleashes its fusion machinery to infiltrate the cell (Hoffmann et al., 2020a; 2020b; Ke et al., 2020; Lan et al., 2020; Shang et al., 2020; Wrapp et al., 2020; Yan et al., 2020).

While essential protein-protein interactions for infectivity have been forthcoming, equally important aspects of SARS-CoV-2 pathobiology have received less attention. For example, amplification of systemic infection requires mass production of functional virions, each of which relies on a specific set of biomolecules to orchestrate the optimal number and spacing of spike trimers in its envelope (Ke et al., 2020; Klein et al., 2020). How assembly occurs efficiently in the crowded cellular environment is unclear. One favored hypothesis is that viral proteins are similarly trafficked to the ER-Golgi intermediate compartment (ERGIC). While cargo receptor-binding likely plays a role, an alternative possibility is that such proteins feature an intrinsic preference for membrane domains of distinct lipid composition (Cattin-Ortolá et al., 2020; Li et al., 2007; Liao et al., 2006; Lu et al., 2008a; McBride et al., 2007; Thorp and Gallagher, 2004). Indeed, certain viruses require association between receptor proteins and specific lipids to trigger endocytosis (Levental et al., 2020; Pelkmans, 2005). Whether this is the case for ACE2 remains to be determined. Regardless, lipid bilayers’ differential propensity to incorporate spike vs. ACE2 might determine whether premature interactions promote unproductive membrane fusion in the cell interior, or if present at the cell surface, fusion of apposing cells (Buchrieser et al., 2020; Cattin-Ortolá et al., 2020; Li et al., 2003; McBride and Machamer, 2010a; Ou et al., 2020; Papa et al., 2020; Xia et al., 2020).

For many enveloped viruses, infection indeed causes fusogenic viral protein display on the host cell plasma membrane, which allows neighboring cells to fuse into multinucleated “syncytia” (Ciechonska and Duncan, 2014; Compton and Schwartz, 2017; Duelli and Lazebnik, 2007). Past studies of respiratory syncytial virus (RSV), human immunodeficiency virus (HIV), and others suggest that cell-cell fusion can play key roles in pathogenicity, whether it be in viral replication, or evasion of the host immune response (Frankel et al., 1996; Johnson et al., 2007; Maudgal and Missotten, 1978). Recent studies on SARS-CoV-2 identified similar syncytia (Buchrieser et al., 2020; Cattin-Ortolá et al., 2020; Ou et al., 2020; Papa et al., 2020; Xia et al., 2020), which may or may not be relevant to patient pathology (Bryce et al., 2020; Giacca et al., 2020; Rockx et al., 2020; Tian et al., 2020). It remains an open question if syncytia are related to viral and host cell membrane composition, and whether their formation provides mechanistic insights into membrane-targeting therapeutics directed toward enveloped viruses including SARS-CoV-2 (Daniels et al., 2020; Zhang et al., 2020).

Here, we address these significant gaps in our understanding of COVID-19 pathobiology by employing a suite of microscopy-based approaches built around the finding that co-cultures of ACE2- and spike-expressing cells amass widespread syncytia. Mechanistically, ACE2-spike clusters assemble at transcellular, synapse-like contacts, which precede fusion pore formation and multinucleation. A high-throughput screen for modulators of cell-cell fusion, involving ∼6000 compounds and >30 spike variants, collectively underscore an essential role for cholesterol-spike association in SARS-CoV-2 infection. Our results suggest that modulation of membrane composition may inhibit viral propagation, and further informs critical lipid-protein assemblies in physiological syncytia and cell adhesion.

## Results

### Syncytia derive from fusion events at synapse-like, spike-ACE2 protein clusters

Given the central role of the ACE2-spike interaction in viral infection (Hoffmann et al., 2020b; Li et al., 2003; Mittal et al., 2020), we sought to develop a live cell microscopy assay of binding and membrane fusion. We generated human osteosarcoma (U2OS) cells, chosen for their flat morphology, stably expressing fluorescently tagged ACE2 or spike (full-length, “FL” vs. receptor-binding domain, “RBD”; see Figure 1A for domain organization), using the B7 transmembrane (“TM”) domain (Liao et al., 2001; Lin et al., 2013) as a control. Upon co-culture, ACE2 and spike RBD cluster at cell-cell interfaces in a binding-dependent manner (Figure 1B). By contrast, and in agreement with others (Buchrieser et al., 2020; Cattin-Ortolá et al., 2020; Ou et al., 2020; Xia et al., 2020), spike FL/ACE2 interactions drove membrane fusion, with the vast majority of cells joining multinucleated syncytia after a day of co-culture (Figure 1C).

**Figure 1.**
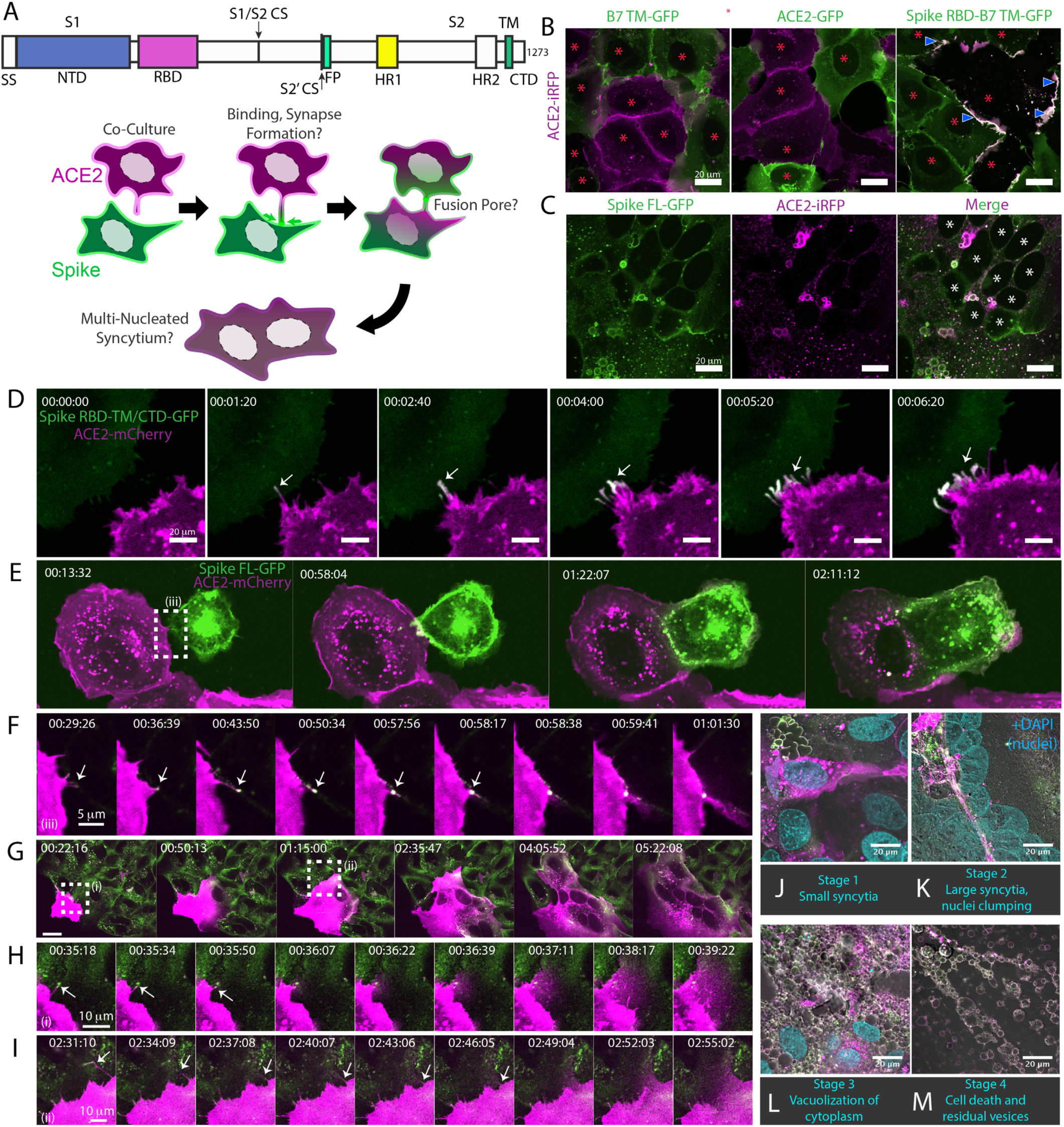
Syncytia derive from fusion events at synapse-like, spike-ACE2 protein clusters. **(A)** Top: Domain structure of a single monomer of the SARS-CoV-2 spike trimer. Domains/motifs is from left to right (see **KEY RESOURCES** table for residue boundaries): SS (signal sequence), NTD (N-terminal domain), RBD (receptor-binding domain), S1/S2 (subdomain-1/2 cleavage site), S2’ (subdomain-2’ cleavage site), FP (fusion peptide), HR1 (heptad repeat 1), HR2 (heptad repeat 2), TM (transmembrane alpha helix), CTD (cytoplasmic domain). Bottom: Cartoon depiction of live cell co-culture assays to detect spike-ACE2 binding and cell-cell fusion. Magenta, acceptor cells (human ACE2-mCherry or ACE2-iRFP); Green, donor cells (GFP-tagged spike variant). **(B)** ACE2-iRFP U2OS (human osteosarcoma) acceptor cells (magenta) co-cultured for 24-hours with U2OS cells expressing GFP-tagged proteins (green): B7 transmembrane (TM, left), ACE2 (middle), spike receptor-binding domain (RBD-TM/CTD, right). Red asterisks indicate single cell nuclei in isolation (no syncytia); arrowhead, synapses (select examples noted). **(C)** Co-culture of U2OS acceptor cells expressing ACE2-iRFP (magenta) with spike full-length (“FL”)-GFP U2OS cells (green). White asterisks: cell nuclei in syncytium. **(D)** Co-culture of ACE2-mCherry (magenta) and spike RBD-TM/CTD-GFP (green) cells for indicated amount of time (hours:minutes:seconds). Arrow: synapse-like interfaces between cells. Scale-bar, 5 µm. See also **Video S1** for long-lived ACE2-Spike FL synapses. **(E)** Similar to Figure 1D, but spike FL-GFP and ACE2-mCherry co-culture. Dashed box indicates site of synapse formation and cell-cell fusion. See **Video S2** for time-lapse movie. **(F)** Zoom-in on synapse formation (arrow, left image) and fusion event (arrow, sixth image from right) of dashed box in Figure 1E. **(G)** ACE2-mCherry cell added to pre-plated spike FL-GFP U2OS cell monolayer (time since ACE2 cell plating indicated). Syncytium forms by multiple cell-cell fusion events (dashed boxes). See Figure 1H,I for zoom-in events (i) and (ii). See **Video S3** for time-lapse movie. **(H)** First cell fusion event (i from Figure 1G) at spike-ACE2 synapse. Time since ACE2-mCherry cell plating indicated. Arrow: retracting synapse prior to cell fusion. **(I)** Similar to Figure 1H but second cell-cell fusion event (ii from Figure 1G). **(J)** Representative image of small syncytia (stage 1) common at early time points following co-culture of ACE2-mCherry (magenta) and spike-GFP (green) U2OS cells but rare at 24-hours (blue, Hoechst DNA stain). **(K)** Similar to Figure 1J, but representative of more common, larger syncytia (stage 2) at 24-hours. Nuclei (blue) clump in center of syncytium. **(L)** Similar to Figure 1J, but representative of typical syncytium with extensive vacuolization (stage 3) at 48-hours. **(M)** Similar to Figure 1J, but representative of remnants (spherical membranous structures) of dead syncytium at 72-hours (stage 4). See also Figure S1**; Videos S1-S3**.

We reasoned that if this co-culture system recapitulates established findings regarding spike/ACE2-mediated viral entry, it might serve as a useful high-throughput proxy assay for infection, without need for enhanced biosafety protocols. To examine this possibility, we first confirmed that fusion events occur following co-culture of spike cells with infection-competent cell lines (VeroE6, Calu3) in absence of ACE2 overexpression, but not with those that do not support infection (Beas2B, U2OS without ACE2) (Figure S1A) (Hoffmann et al., 2020b). We further validated the relevance of the assay by showing that domains required for virus-cell entry (e.g. binding domain, RBD; fusion machinery, heptad repeats and fusion peptide) are needed for cell-cell fusion (Figure S1B). Similar to results obtained with infectious virus (Hoffmann et al., 2020a), fusion required the spike S2’ cleavage site but not the S1/S2 site (Figure S1C). Finally, different fluorescent tags (GFP, mCherry, iRFP) gave similar results (Figure S1D,E), expanding the fluorescent toolkit for live cell studies.

We hypothesized that spike/ACE2-mediated syncytia form in a stepwise manner, which might illuminate mechanisms of formation and pathogenesis. We performed live cell microscopy of co-cultures, documenting dozens of fusion events preceding large syncytia. Invariably, contact between opposite cell types (spike vs. ACE2) results in near instantaneous accumulation of spike protein clusters at ACE2-containing membrane protrusions (Figure 1D-E). These punctate structures are long-lived (minutes) **(Video S1)**, similar to physiological synapses (e.g. neuronal, immunological) (Cohen and Ziv, 2019; Dustin, 2014). In all observed cases, fusion events proceed from such synapses (Figure 1E-F; **Video S2)**, often within a few minutes of their formation, but frequently following longer durations of time. In most (but not all) examples, fusion pore dilation follows retraction of an individual spike cluster toward the interior of an ACE2 cell (Figure 1F-I), suggesting that motility-associated mechanical forces (e.g. actomyosin contractility) and/or endocytosis is pivotal to overcoming the energetic barrier to lipid bilayer mixing. When cells are plated at high density, most “primary” fusion events occur within 60 minutes (Figure 1G-J), with latter “secondary” amalgamation of small syncytia into progressively larger structures (Figure 1K**; Video S3)**. Over time, syncytia undergo vacuolization, likely from fusion-driven collapse of intracellular organelles into hybrid compartments (Figure 1L). By 48-72 hours, cells disintegrate into immobile spike/ACE2-coated vesicles, having eaten themselves from within (Figure 1M).

### Syncytia are a defining pathological feature of COVID-19

While clearly useful for interrogating spike domains that modulate membrane fusion, a critical gap in our knowledge concerns the pathophysiological relevance of the syncytia themselves. Given the cytotoxic consequences of cell fusion (Figure 1M), an appealing hypothesis is that ACE2/spike-mediated cell vacuolization contributes in part to the diffuse alveolar damage observed in the lungs of COVID-19 patients (Menter et al., 2020). Intriguingly, SARS-CoV-2 spike is a particularly potent mediator of syncytia formation relative to both SARS-CoV-1 spike and commonly studied fusogens (e.g. p14 FAST, MYMK/MYMX) (Bi et al., 2017; Chan et al., 2020b) based on side-by-side comparisons (Figure 2A-C).

**Figure 2.**
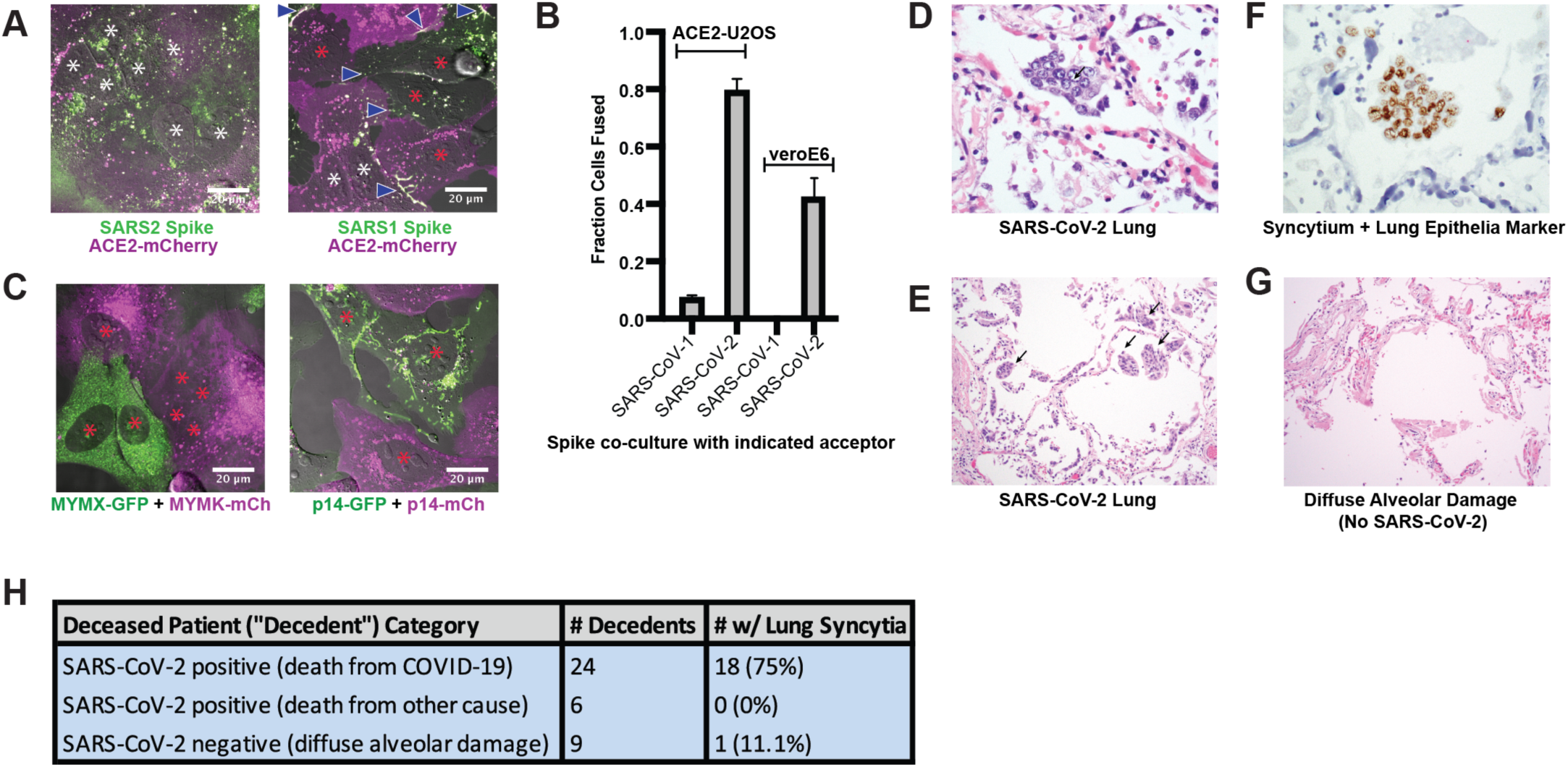
Syncytia are a defining pathological feature of COVID-19. **(A)** ACE2-mCherry (magenta) U2OS cells co-cultured with GFP-tagged (green) SARS-CoV-2 spike cells (left) or SARS-CoV-1 spike cells (right) for 24-hours. White asterisks indicate nuclei in syncytia; red, in isolation; arrowhead, synapses (select examples noted). **(B)** Quantification of Figure 2A by percent cells fused (also tested VeroE6 donor cells, which express endogenous ACE2). Mean and SEM: n=4 biological replicates (16 images per). **(C)** Indicated fusogen-expressing cells lines co-cultured for 24-hours. Red asterisk indicates nuclei of single cells (not in syncytia). **(D)** Lung from SARS-CoV-2 decedent demonstrating syncytia formation (H&E stained section, 400x original magnification). Syncytium labeled with arrow. **(E)** Similar to Figure 2D, but sample obtained from different deceased COVID-19 patient and at 400x magnification. Syncytium labeled with arrow. **(F)** Immunohistochemistry for lung epithelia marker TTF-1 (NKX2-1; brown) showing nuclear positivity in the syncytial cells (400x original magnification). **(G)** Lung from control decedent with diffuse alveolar damage unrelated to SARS-CoV-2 infection (died pre-2019), showing hyaline membranes (remnants of dead cells; bright pink) but no syncytia (H&E stained section, 100x original magnification). **(H)** Table summarizing decedents examined for syncytia pathology.

We speculated that this superior ability to promote cell-cell fusion might be reflected by unique pathological attributes *in vivo*. If so, we predicted that syncytia would be readily detected in the lungs of COVID-19 patients. To test this, we histologically evaluated lung samples from 24 deceased patients (“decedents”) with diffuse alveolar damage secondary to SARS-CoV-2 infection; 6 decedents who were positive for SARS-CoV-2 but did not have pulmonary manifestations (died of other causes); and 9 control decedents with diffuse alveolar damage (died prior to SARS-CoV-2 discovery in 2019). Multinucleated syncytia were detected in 18 of 24 decedents who died as a direct consequence of SARS-CoV-2 infection (Figure 2D-H), a finding supported by other patient cohorts (Giacca et al., 2020; Tian et al., 2020). These syncytia were of lung epithelial origin, as demonstrated by nuclear staining for TTF-1 (NKX2-1) (Figure 2F). In contrast, only 1 of the 9 decedents with diffuse alveolar damage from other causes demonstrated multinucleated syncytia, indicating that these syncytia are not a common feature of lung inflammation (Figure 2G,H). They were also absent in lung tissue from the 6 SARS-CoV-2 decedents who did not show pulmonary manifestations and died of other causes. Thus, pathological syncytia are a direct consequence of pulmonary involvement by SARS-CoV-2 (Figure 2H). These syncytia, however, were generally not positive for the SARS-CoV-2 nucleocapsid protein, similar to previous reports (Bryce et al., 2020; Rockx et al., 2020). Thus, we cannot rule out a yet-to-be identified pulmonary abnormality specific to SARS-CoV-2 infection (but spike-independent), or related to free spike proteins in the alveolar debris, in pathological syncytia formation.

### A novel high-throughput screening platform identifies modulators of syncytia formation

Given the potential pathological relevance of syncytia and their ability to interrogate SARS-CoV-2 entry, we sought to adapt our cell model into a high-throughput assay to uncover molecular mechanisms and drug targets. We developed and evaluated three different fixed-cell microscopy assays, each of which used fluorescent proteins as readouts for fusion (Figure S2B,D), with total nuclei number serving as a toxicity measure. Two of these assays (human U2OS-ACE2 vs. monkey VeroE6 heterokaryon) leveraged RNA-binding proteins’ ability to shuttle between the nucleus and cytosol (Iijima et al., 2006; Zinszner et al., 1997), with nuclear co-localization of mCherry/GFP reflecting cell fusion. The third assay used split-GFP (Buchrieser et al., 2020; Feng et al., 2017), which only fluoresces when its two halves come into contact (e.g. after fusion). After careful assessment, the U2OS-ACE2 heterokaryon system was shown to be the superior assay based on its Z’-factor (0.85), a measure of separation between positive (spike RBD/ACE2 co-culture, no fusion) and negative (spike FL/ACE2 co-culture) controls (Figures 3A, S2D); generally Z’-factor > 0.5 is considered excellent for a high-throughput assay (Zhang et al., 1999). To determine an optimal time point for quantification, co-cultures were performed for 1-6 hours: fusion was detectable by 1-hour, and Z’-factor peaked by 4-5 hours (Figure S2B).

**Figure 3.**
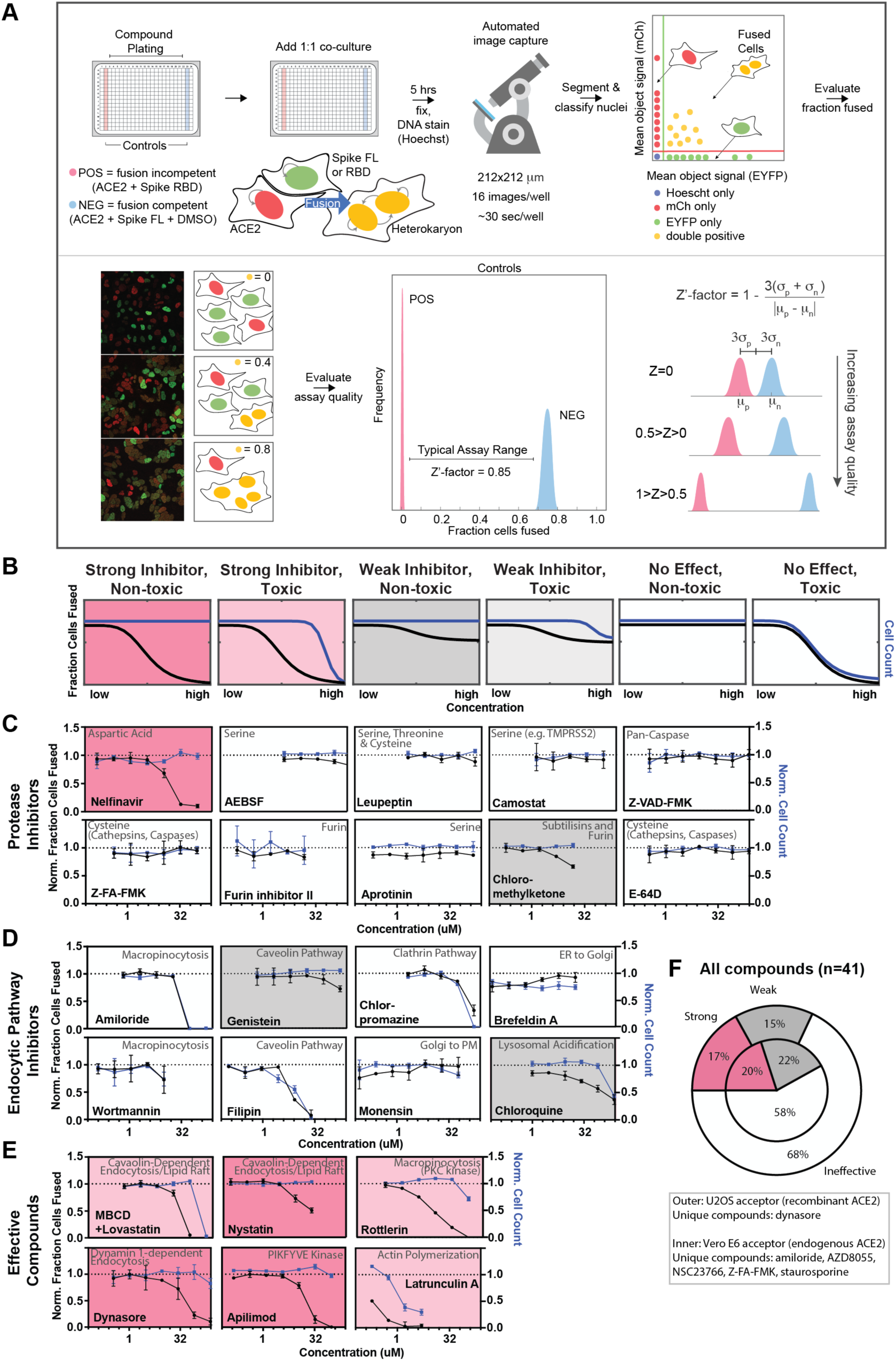
A novel high-throughput screening platform identifies modulators of syncytia formation. **(A)** Heterokaryon assay workflow overview (top) and characterization (bottom). Equal parts acceptor cells (express ACE2-iRFP and FUS-mCherry) plus donor cells (express spike FL-iRFP and HNRNPA1-EYFP) are co-cultured in 384-well microtiter plate: positive control (spike RBD, red column), negative control (DMSO, blue column), test compounds (other columns). After 5-hours, cells are fixed and nuclei are stained (Hoechst) then identified/segmented by automated confocal microscopy. Fraction cells fused is determined by percent co-positive (mCherry and EYFP) nuclei. Sample images and schematic interpretation of a positive control well (top), test well with reduced fusion (middle), negative control well (bottom). Z’-factor measures window size with higher score indicating a more robust screening platform. **(B)** Schematic of assay dose-response and interpretation. Fraction of cells fused (black curve) relative to cell count (toxicity measure, blue curve), both normalized by plate negative control, indicate compound efficacy (pink, strong inhibitor; gray, weak inhibitor; white, no-effect). **(C)** A panel of spike protease inhibitors (n=10), which includes antagonists of both established SARS-CoV-2 entry pathways (cathepsin-dependent endocytosis vs. TMPRSS2/furin-mediated direct fusion), was tested. Mean and SEM: n=4 biological replicates (16 images per). **(D)** Similar to Figure 3C, but displaying dose-response relationships for select inhibitors of indicated routes of endocytosis (e.g. clathrin, macropinocytosis) and steps in secretory pathway (e.g. ER-Golgi transport). See Figure S2F for additional tested compounds. **(E)** Similar to Figure 3C, but displaying dose-response relationships for compounds that strongly inhibit cell-cell fusion in ACE2-U2OS heterokaryon assay. See Figure S3 for similar effect in VeroE6 assay (no exogenous ACE2 expression). **(F)** Summary of targeted drugs (n=41) in U2OS and Vero based co-culture assays. Identified inhibitors are largely similar between cell types. Cell type-specific effects are noted. See also Figures S2, S3; **Table S4**.

Armed with a tractable assay (Figure 3A,B), we sought to characterize essential pathways for fusion, employing dose-response studies of a panel of drugs with well-characterized mechanism of action. Given that protease-mediated S2’ cleavage is essential for cell-cell fusion (Figure S1C), we anticipated a large effect for specific classes of protease inhibitors. We thus tested a panel of spike protease inhibitors (n=10), which included antagonists of both established SARS-CoV-2 entry pathways (cathepsin-mediated endocytic vs. TMPRSS2/furin-dependent direct fusion) (Hoffmann et al., 2020a; 2020b; 2020c; Kawase et al., 2012; Millet and Whittaker, 2014; Ou et al., 2020; Shirato et al., 2018; Zhou et al., 2015). Surprisingly, the antiretroviral protease inhibitor nelfinavir was unique in blocking fusion (Figure 3C). Given that other serine protease inhibitors (AEBSF, leupeptin, camostat) lacked efficacy, inhibition by nelfinavir may be related to its proteolysis-independent targets (Brüning et al., 2013; De Gassart et al., 2016; Kirby et al., 2011).

Next, we screened drugs that target specific routes of endocytosis or steps in the secretory pathway. Notably, no single endocytic route (clathrin, caveolae, macropinocytosis) was essential (Figures 3D; S2E,F), and markers of each pathway did not co-localize with spike at sites of membrane-fusion or in the vesicles that resulted (Figure S2A). Nevertheless, certain drugs prevented syncytia formation (Figure 3E,F). For example, apilimod, a promising COVID-19 drug candidate that inhibits PIKFYVE kinase (Cai et al., 2013; Kang et al., 2020; Riva et al., 2020), was particularly potent (Figure 3E). Inhibition of actin polymerization blocked fusion (Figure 3E), consistent with a putative role for cortical actomyosin-generated mechanical forces in fusion pore formation (Figure 1) (Chan et al., 2020b; Shilagardi et al., 2013). Finally, several drugs that perturb membrane lipid composition were identified (MBCD/lovastatin, genistein, nystatin) (Figure 3E). These compounds could conceivably act by permeabilizing the plasma membrane. To rule out this possibility, cell fusion was examined following treatment with membrane permeabilizers digitonin or ethanol, both of which had no effect on fusion at non-toxic concentrations (Figure S2C). Finally, most compounds acted similarly in the VeroE6 heterokaryon assay (Figures 3F; S3), which expresses endogenous ACE2. We therefore conclude that our syncytium-based screening platform can identify fusion-inhibiting drugs that act independently of canonical entry pathways and might uncover yet-to-be described determinants of membrane fusion.

### A drug repurposing screen implicates membrane lipid composition in cell-cell fusion

To gain further mechanistic insight into cell-cell fusion, we performed a drug repurposing screen of ∼6000 small molecules at 30 µM (Figure 4A). Of these, 167 (2.8%) were inhibitory “hits”, which signified non-toxic compounds with a decrease in fusion greater than 3-standard deviations from the mean (z-score<-3) (Figure 4B**; Table S1**). To validate, we performed 7-point dose-responses for the top-80 most potent compounds, the vast majority of which replicated (Figures 4C; S4A,B). Compounds were then unblinded to select experimenters. Reassuringly, 23 of these hits were redundant across the combined libraries (Figure S4B), and several were identified in previous virus entry screens (Caly et al., 2020; Carbajo-Lozoya et al., 2012; Hoffmann et al., 2020c; Kindrachuk et al., 2015; Riva et al., 2020; Yamamoto et al., 2016). We eliminated batch-dependent effects, purchasing top compounds from independent vendors, and replicating dose-response measurements for 23 of 24 (Figure S4A). To assess cell type specificity, dose-response studies for the same molecules were performed in the VeroE6 heterokaryon assay (Figure S4A). In almost all cases, inhibition of fusion occurred at lower compound concentrations relative to the U2OS assay, possibly due to differences in ACE2 levels between cell lines.

**Figure 4.**
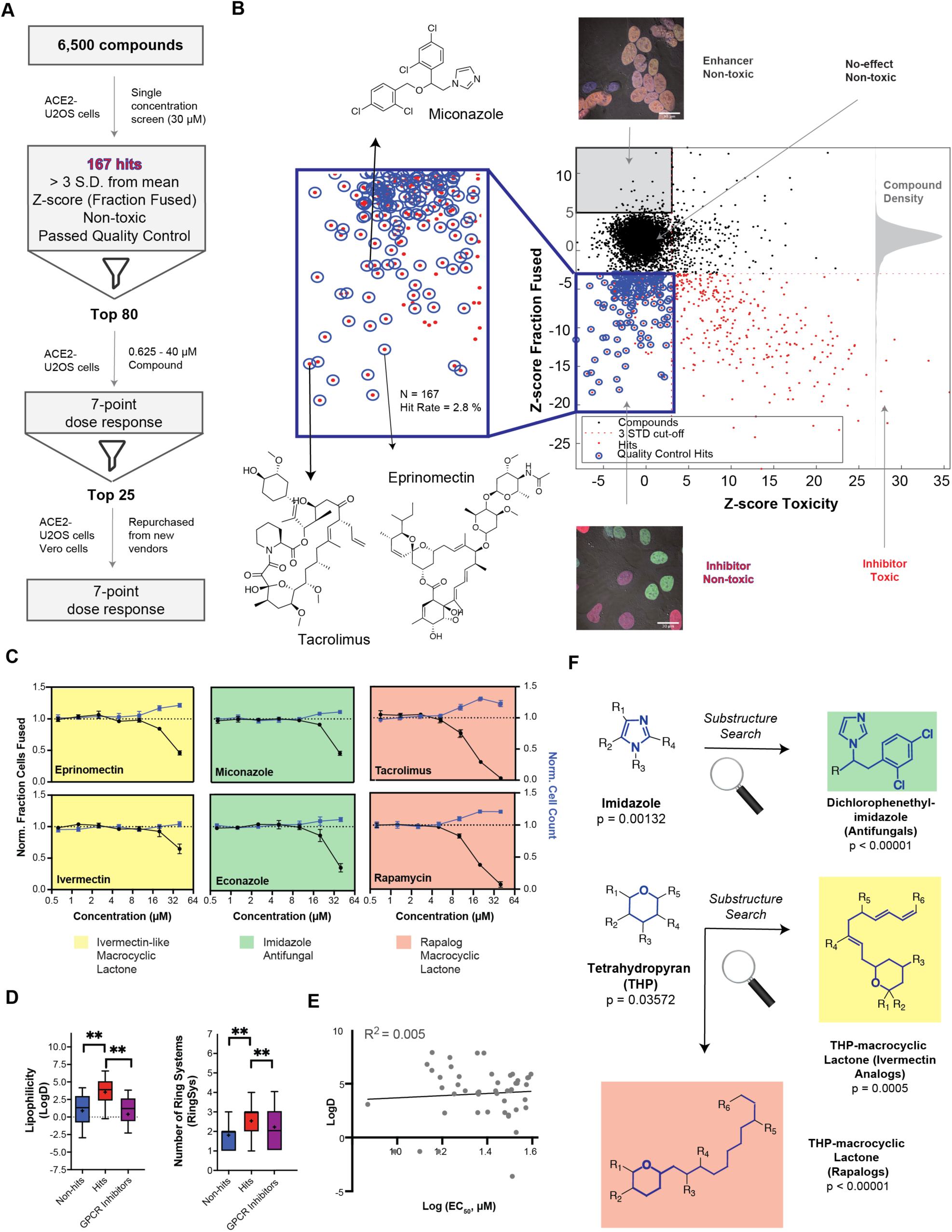
A drug repurposing screen implicates membrane lipid composition in cell-cell fusion. **(A)** High-throughput pipeline and workflow of small molecule screen (∼6000 compounds, 30 µM) in ACE2-U2OS heterokaryon assay. “Hits” refer to non-toxic compounds with a decrease in fusion of >3 standard deviations relative to plate negative control. 7-point dose-response was determined for top-80 inhibitors, followed by validation of select compounds (n=24; obtained from different vendors) in both ACE2-U2OS and VeroE6 heterokaryon assays. **(B)** Results of compound screen. Plot: fraction fused vs. toxicity z-score. Red dots indicate compounds with decreased fusion (z-score<-3); blue inset, potential hits following toxicity filtering (z-score<3); blue circles, quality-controlled hits (inhibitory, non-toxic compounds with normal fluorescence); gray inset, compounds with increase in fraction cells fused (z-score>5) but no toxicity (“enhancers”, see Figure 6J); right histogram, compound density as function of fraction fused z-score. Chemical structures are displayed for select validated hits. See **Table S1** for raw data. **(C)** Dose-responses for select hits in enriched substructure classes (see Figure 4F): imidazoles (e.g. azole antifungals, green) and macrocyclic lactones (ivermectin-like, yellow; rapalogs, pink). Mean and SEM: n=3 biological replicates (16 images per). **(D)** Box-and-whisker plots of select physicochemical properties (lipophilicity, logD; ring systems) for non-hits (blue), inhibitory hits (red), and GPCR inhibitors (purple) as calculated in ChemAxon. Boxes encompass 25-75% of variance; whiskers, 10-90%. Mean values are indicated by “+”; median values, lines. Statistical significance was assessed by Mann-Whitney U test: P-values of <0.05 and <0.001 are represented by * and **, respectively. **(E)** Lack of correlation between inhibitory hit EC_50_ (see Figure S4A) and lipophilicity according to linear regression analysis conducted in GraphPad Prism. **(F)** Three substructure classes based on two scaffolds were identified to have high statistical enrichment in hits over non-hits: dicholorophenethyl-imidazoles (found in azole antifungals, green) and tetrahydropyrans with alkyl moieties (found in macrocyclic lactones; yellow and pink indicate ivemectin-like and rapalog compounds, respectively). See also Figures S4, S5; **Table S1**.

Given their unusually high EC_50_ (>10 µM) (Figure S4A), identified small molecules might act directly on the lipid bilayer (Tsuchiya, 2015), possibly by virtue of shared physicochemical or structural features. To assess this, we compared 20 physicochemical parameters (ChemAxon) for non-hits vs. hits, using GPCR inhibitors (∼35% of FDA-approved drugs) (Sriram and Insel, 2018) as a control library (Figures 4D; S5A,B). Among several statistically significant differences, hits were more lipophilic (LogD) and featured a greater number of ring systems (Figure 4D). Reassuringly, little correlation was observed between EC_50_ values and lipophilicity (Figure 4E), indicating that the trend is not a result of a general increase in lipophilicity with avidity, as is commonly observed for promiscuous compounds in phenotypic screens (Tarcsay and Keserű, 2013).

Next, we asked whether specific chemical scaffolds are over-represented in hit compounds relative to ineffective compounds (Figures 4F; S5C). Two scaffold classes (and corresponding substructures) reached particularly high statistical enrichment: dicholorophenethyl-imidazoles (found in azole antifungals) and tetrahydropyran-containing macrocyclic lactones (found in both ivermectin- and rapamycin-like compounds) (Figures 4C,F; S5C). Such molecules can directly interact with the plasma membrane (François et al., 2009), perturbing cholesterol (e.g. production, transport) (Bauer et al., 2018; Mast et al., 2013; Trinh et al., 2017; Xu et al., 2010), and have been implicated as promising repurposed drugs for COVID-19 treatment, albeit by different mechanism of action (Caly et al., 2020; Gordon et al., 2020; Kindrachuk et al., 2015; Rajter et al., 2020).

### Highly unusual membrane-proximal regions of spike are needed for fusion

Based on the prevalence of lipophilic hits from the small-molecule screen, we posited that membrane-proximal regions of spike and/or ACE2 associate with essential plasma membrane lipids (e.g. cholesterol) to facilitate cell-cell fusion. To test this, we replaced the transmembrane and cytoplasmic domains of both ACE2 and spike with the previously used B7 TM (Figure 1B**, Table S4**). While “chimeric” ACE2 similarly promoted cell fusion relative to wild-type (WT), chimeric spike protein lost this ability (Figure 5A). To determine critical elements that differentiate WT and chimeric spike from one another, we mutated its transmembrane (TM) and cytoplasmic domains (Figure 5B), assessing fusion in co-culture models (Figures 1A, 3A). Replacement of spike’s transmembrane domain with single-pass TMs of unrelated proteins (B7, ITGA1) blocked fusion, despite similar subcellular localization and ACE2-binding (Figures 5C,L; S6A-C). Inclusion of a small extracellular motif of B7 not only eliminated fusion, but also impaired the ability of the chimeric spike to form synapse-like clusters with ACE2 (Figure 5A). This is likely indicative of an essential role of spike’s membrane-proximal aromatic residues in cholesterol engagement (Hu et al., 2019a), as suggested by work on related coronaviruses (Corver et al., 2009; de Jesus and Allen, 2013; Epand et al., 2003; Liao et al., 2015; Lu et al., 2008b; Meher et al., 2019).

**Figure 5.**
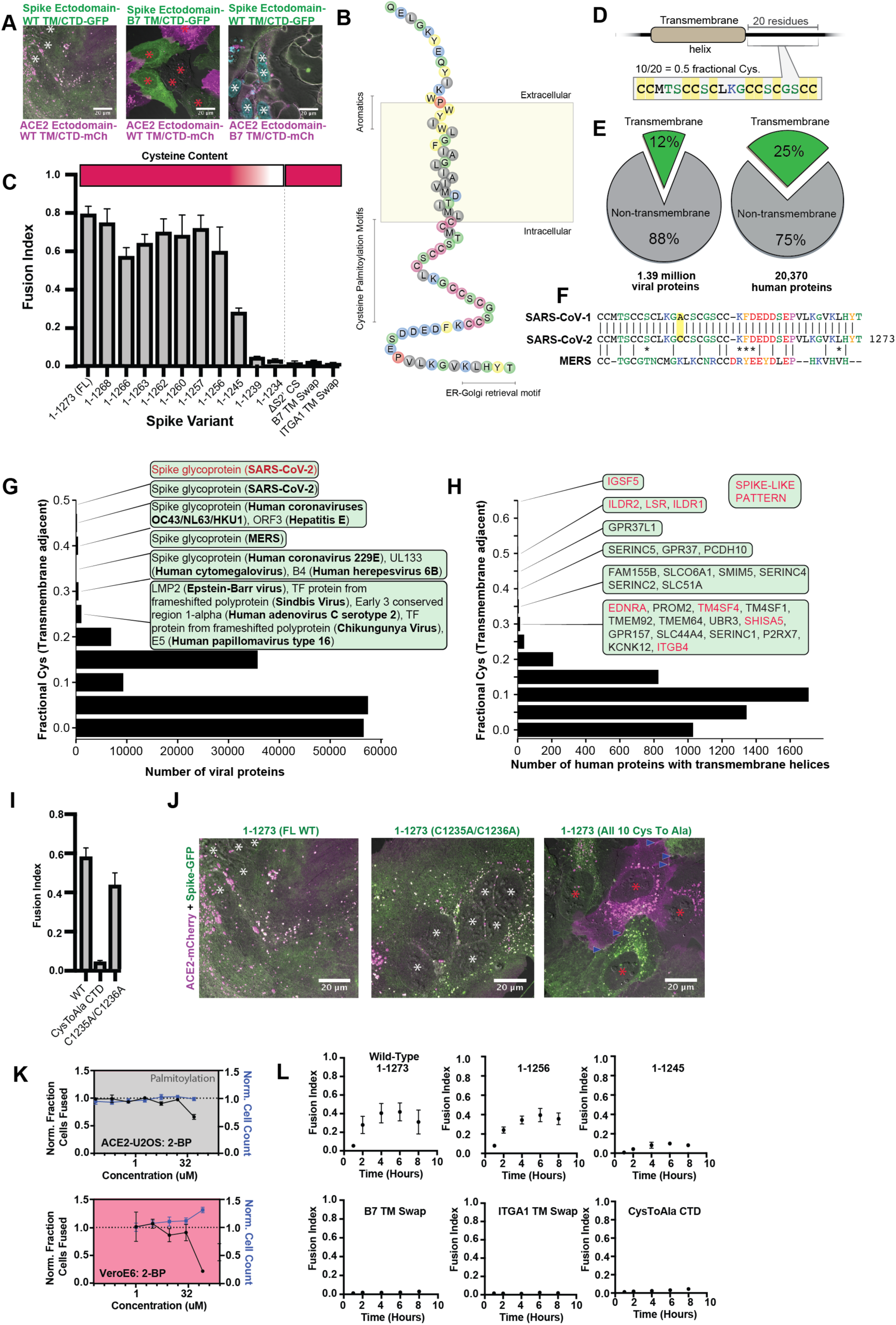
Highly unusual membrane-proximal regions of spike are needed for fusion. **(A)** Representative images of co-cultured (24-hours) U2OS cell lines, stably expressing indicated fluorescently tagged ACE2 or spike. “B7 TM” indicates swap of endogenous transmembrane (TM) and cytoplasmic domain (CTD) of spike or ACE2 with that of the monomeric, single-pass, B7 transmembrane protein, along with its membrane-proximal extracellular region (30-amino acid spacer). White asterisks indicate nuclei in syncytia; red, in isolation. Note lack of arrowheads (synapses) in middle condition. See **Table S4** for residue composition of such “chimeric” proteins. **(B)** Graphical representation of SARS-CoV-2 spike’s TM alpha-helix and membrane-proximal regions, with residues colored by chemical properties (yellow, aromatic; cysteine, magenta; hydrophobic, gray; non-charged hydrophilic, green; charged hydrophilic; blue; proline, red). Of note: aromatic-rich region at ectodomain-TM interface, cysteine-rich cytoplasmic domain (CTD). **(C)** ACE2-U2OS heterokaryon assay but with co-cultured HNRNPA1-EYFP cells expressing spike variants (indicated). Relative CTD cysteine content for variants is depicted with heat map (top; dark red = more cysteines, white = none). Mean and SEM: n=4 biological replicates (16 images per). See Figure S6B for all tested spike variants in both ACE2-U2OS and VeroE6 heterokaryon assays; Figure S6C, for representative images of ACE2-mCherry U2OS cells co-cultured with GFP-tagged spike variants. **(D)** Cartoon representation of SARS-CoV-2 spike with highest cysteine content in a 20-amino acid sliding window, which guided bioinformatic analyses shown in Figure 5E-H. **(E)** Schematic of bioinformatic analysis performed, whereby 20-residue windows around the N- and C-terminal sides of transmembrane helices were scanned for local cysteine density. Pie charts: summary of total set of viral proteins retrieved and analyzed for the human virus proteome (left) and human proteome (right); green slice references proportion of proteins with one or more predicted transmembrane helixes. **(F)** Conservation of cysteine-rich CTD between spike proteins of highly pathogenic human coronaviruses. The only difference between the CTD of SARS-CoV-1 and SARS-CoV-2 is acquisition of an additional cysteine in the latter. MERS is substantially different, yet retains enrichment of cysteines. **(G)** Histogram of fractional cysteine scores for viral proteins, with high-fraction hits explicitly annotated. SARS-CoV-2 spike protein has the highest local cysteine density of any viral protein, closely followed by spike proteins from other coronaviruses. **(H)** Similar to Figure 5G, but for human proteins with one or more predicted transmembrane helix. Red: “spike-like” transmembrane proteins with high cytoplasmic cysteine content and aromatic residues at ectodomain-membrane interface. **(I)** Similar to Figure 5C, but using spike variants with cysteines mutated to alanine (2 of 10 vs. 10 of 10). **(J)** Representative images for 24-hour co-culture of ACE2-mCherry (magenta) U2OS cells with those expressing GFP-tagged spike variant (green). White asterisks indicate nuclei in syncytia; red, those in isolation; arrowhead, synapses (select examples noted). **(K)** Dose-response relationship for 2-bromopalmitate (2-BP, inhibitor of cysteine palmitoylation), in both ACE2-U2OS and VeroE6 heterokaryon assays. Blue indicates number of nuclei (proxy for toxicity); black, percent cells fused; both normalized to DMSO control. Mean and SEM: n=4 biological replicates (16 images per). **(L)** Similar to Figure 5C, but assesses kinetics of fusion by varying co-culture time prior to fixation. See Figure S6A for other tested spike variants. See also Figure S6**; Tables S2-S4.**

In parallel, we serially truncated the spike cytoplasmic domain (CTD). Removal of its COPII-binding, ER-Golgi retrieval motif (Cattin-Ortolá et al., 2020; McBride et al., 2007) (1-1268) had no effect, nor did deletion of its subsequent acidic patch (1-1256) (Figures 5C,L; S6A-C). However, removal of an additional 11 amino acids (1-1245) decreased fusion, and further truncation (1-1239) completely blocked it (Figures 5C,L; S6A-C). Relative fusion correlated with overall cysteine content of the CTD (Figure 5C). These findings are consistent with previous studies on similar coronaviruses, which suggested that membrane-proximal cysteines are post-translationally modified with palmitoylated lipid moieties (McBride and Machamer, 2010a; Petit et al., 2007; Sobocińska et al., 2017).

Palmitoylated proteins typically feature only a few cysteines available for modification (Chlanda et al., 2017; Wan et al., 2007). We wondered whether spike CTD’s peculiarly high cysteine content was unique amongst viral proteins, and performed a bioinformatic analysis of all viral transmembrane proteins, ranking them on maximal cysteine content in 20 amino-acid sliding windows (Figure 5D-G). Of all proteins in viruses that infect humans, SARS-CoV-2 spike features the highest cysteine content, followed closely by spike proteins in related coronaviruses, then hepatitis E ORF3 (Figure 5G**; Table S2**); it should be noted that ORF3 is palmitoylated and critical to viral egress (Ding et al., 2017; Gouttenoire et al., 2018). Consistent with studies on similar coronavirus spike proteins (Liao et al., 2006; McBride and Machamer, 2010a; Petit et al., 2007), mutagenesis of all spike cysteines to alanine severely diminishes cell-cell fusion in both U2OS and Vero models (Figures 5I-L; S6B-C). To examine the role of cysteine palmitoylation, we assessed fusion upon treatment with palmitoylation inhibitor, 2-bromopalmitate (2-BP) (Martin, 2013). The effect was modest in U2OS cells, but clear in Vero cells, suggesting that cysteine palmitoylation is indeed likely central (Figure 5K). Thus, both cysteine palmitoylation and unique characteristics of its aromatic-rich transmembrane domain known to associate with cholesterol are essential for spike’s role in membrane fusion.

Given the synapse-like structures observed in living cells (Figure 1), we asked whether SARS-CoV-2 spike features amino acid motifs similar to human proteins that drive similar assemblies. Of the thousands of human transmembrane proteins, only 15 were “spike-like”, featuring both high membrane-proximal cysteine and aromatic content (Figure 5H**; Table S3**). Remarkably, the top four are all critical to forming specific types of adhesion junctions: three to tricellular tight junctions (ILDR1, ILDR2, LSR) and one to kidney/intestine tight junctions (IGSF5) (Figure S6D) (Higashi et al., 2013; Hirabayashi et al., 2003). In light of the important role for palmitoylation in tricellular tight junction assembly (Oda et al., 2020), these findings suggest that SARS-CoV-2 operates by a similar mechanism to promote adhesion and transcellular interfaces.

### Spike requires membrane cholesterol for fusion but via a raft-independent mechanism

Together, these data suggest that membrane fusion requires spike association with specific elements of the plasma membrane. If so, such assemblies would display slow dynamics relative to transmembrane proteins that more freely diffuse in the two-dimensional lipid bilayer. To test this idea, we utilized fluorescence recovery after photobleaching (FRAP) to determine the recovery rate of a fluorescent molecule in a bleached region, and thereby infer relative apparent molecular diffusion coefficients (Soumpasis, 1983). FRAP experiments were performed on a series of GFP-tagged spike variants and controls (B7 TM and ACE2) to determine whether its transmembrane domain and/or cysteine-rich CTD influence diffusion. Recovery for GFP-tagged ACE2, B7 TM-anchored RBD, and the B7 transmembrane control were similar (Figure 6C), approximating diffusion times for commonly studied transmembrane proteins (Day et al., 2012). In contrast, RBD attached to the native TM/CTD of spike featured significantly reduced recovery, with FL spike displaying even slower dynamics (Figure 6A-C). Swapping the B7 TM for spike TM/CTD rescued the rapid recovery, whereas exchange of just the TM or removal of cysteine-containing regions had an intermediate effect (Figure 6C). Conversely, deletion of regions shown to bind specific intracellular proteins (e.g. COPII-binding ER-Golgi retrieval motif, 1268-1273) had no effect (Figure 6C), implicating lipid-protein and not protein-protein interactions in spike’s dynamics.

**Figure 6.**
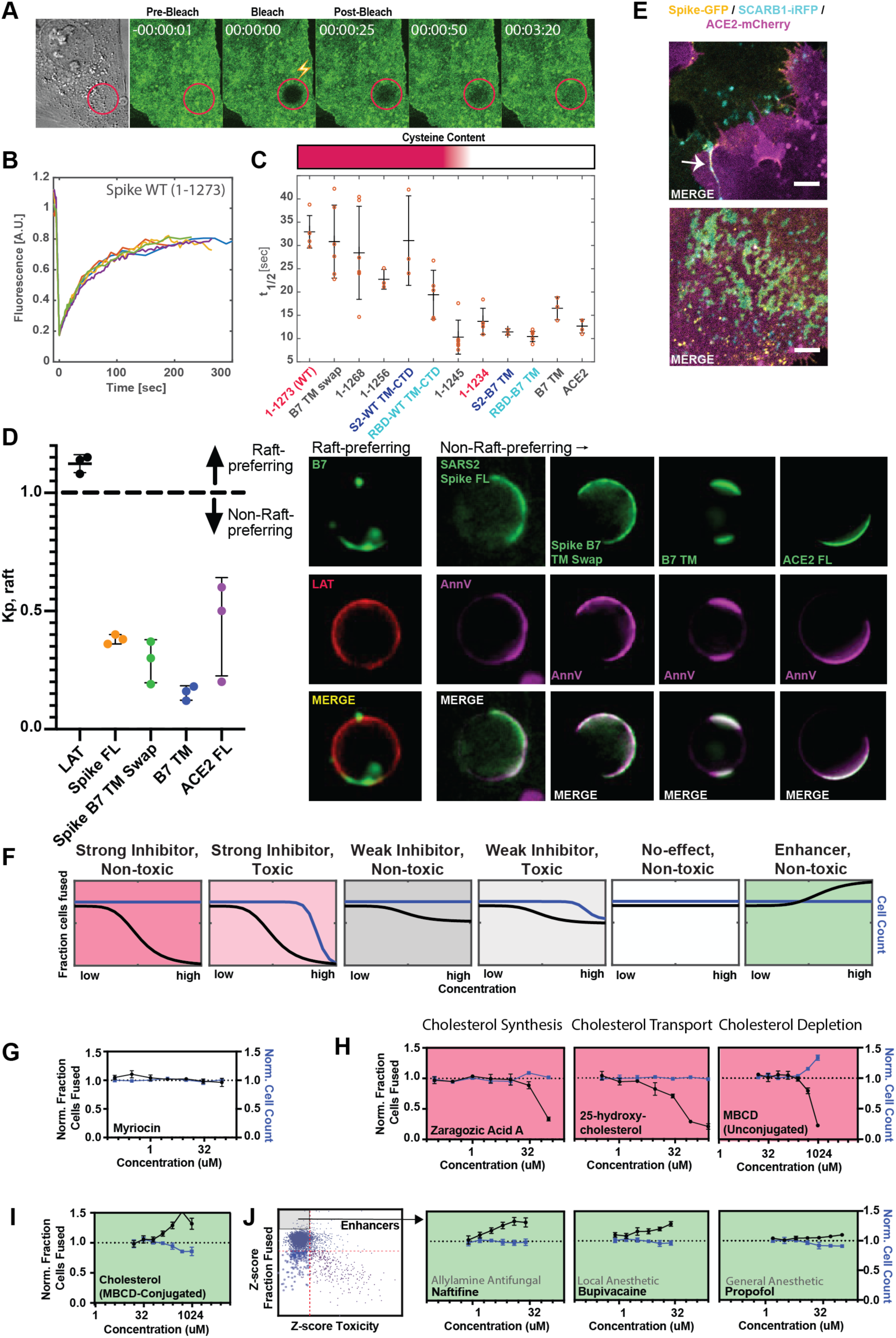
Spike requires membrane cholesterol for fusion but via a raft-independent mechanism. **(A)** Representative trial for fluorescence recovery after photobleaching (FRAP) of spike FL-GFP (green) on U2OS cell plasma membrane. Time since bleach (lightning bolt) of region of interest (red circle) is indicated. **(B)** Quantification of Figure 6A and related trials, with each colored line specifying a separate FRAP experiment (n=6 total). **(C)** Calculated half maximal fluorescence recovery (t_1/2_) for indicated GFP-tagged spike variants. Each hollow red dot indicates the t_1/2_ for a single FRAP trial. Mean and SEM: n=4-6 technical replicates (1 per cell). Heat map (top): relative cysteine content of tested spike variant (dark red = high cysteine content, white = none). **(D)** *Ex vivo* phase separation assay for relative partitioning (K_p_) into lipid raft ordered phase (L_o_) of giant plasma membrane vesicles (GPMVs). Left: quantification of GFP-tagged protein K_p_ with DHPE serving as lipid raft/L_o_ marker; AnnV, non-raft/L_d_ protein marker; LAT, raft/L_o_ protein marker. Mean and SEM: n=3 biological replicates (colored dot; >10 GPMV technical replicates per). Right: representative images. **(E)** U2OS cells expressing spike-GFP (yellow) and SCARB1-iRFP (cyan) were co-cultured with ACE2-mCherry (magenta) cells and cell-cell fusion events were captured with live cell microscopy. Representative images show co-localization between spike and SCARB1 at synapses that precede fusion (top, arrow); and in extracellular deposits (bottom). Scale-bar, 5 µm. See Figure S7A,B for individual fluorescence channels. **(F)** Graphical schematic for ACE2-U2OS assay dose-response and interpretation. Fraction of cells fused (black curve) relative to cell count (blue curve), both normalized by the plate negative control, indicates compound effectiveness (pink, strong inhibitor; gray, weak inhibitor; white, no-effect; green, enhancer). **(G)** Lack of dose-dependent inhibition of fusion by sphingolipid-depleting, and raft-disrupting drug, myriocin, in ACE2-U2OS heterokaryon assay. Mean and SEM: n=4 biological replicates (16 images per). **(H)** Similar to Figure 6G, but measuring effect of cholesterol-lowering drugs in ACE2-U2OS heterokaryon assay. Mean and SEM: n=4 biological replicates (16 images per). **(I)** Similar to Figure 6G, but with MBCD-conjugated cholesterol, which increases plasma membrane cholesterol content. See Figure S7E for controls (i.e. other MBCD-conjugated lipids). **(J)** Similar to Figure 6G, but testing potential fusion-enhancing compounds (see Figure 4B, gray inset), which include allylamine antifungals (e.g. naftifine) and anesthetics (e.g. bupivacaine, propofol). See Figure S7F for similar effects by other small molecules belonging to these compound classes. See also Figure S7**; Table S4**.

Given that membrane-proximal regions (**Table S4**) of spike regulate diffusivity and fusogen behavior, an intriguing possibility is that such features conspire to facilitate engagement of cholesterol-rich membrane domains (or “lipid rafts”) (Levental et al., 2020; Pelkmans and Helenius, 2003; Simons and Ikonen, 1997). Our findings on the requirement for spike’s cysteine residues in fusion is interesting in this context, since palmitoylation of other proteins can drive association with these 10-50 nm protein-lipid clusters in the plasma membrane (Levental et al., 2010). While challenging to study in living cells, lipid rafts can be readily interrogated using *ex vivo*, phase separation assays as a micron-scale proxy for cholesterol association at the nano-scale (Levental and Levental, 2015; Veatch, 2007; Veatch and Keller, 2003). A particularly powerful example employs chemical-induced giant plasma membrane vesicles (GPMVs) from cells expressing a protein of interest (e.g. spike-GFP), allowing membrane components to reach equilibrium at reduced temperature (Baumgart et al., 2007; Holowka and Baird, 1983; Levental et al., 2009; Sengupta et al., 2008; Veatch et al., 2008). Given previous results using indirect detergent fractionation readouts (McBride and Machamer, 2010b; Petit et al., 2007), we were surprised to observe that SARS-CoV-2 spike does not partition strongly into GPMVs’ dense, ordered phase (L_o_) (Figure 6D), which is enriched for cholesterol and sphingolipids (Levental et al., 2009; 2011). Moreover, in our cell fusion assay, treatment with the raft-disrupting drug myriocin, which depletes sphingolipids from the plasma membrane (Castello-Serrano et al., 2020), did not inhibit fusion (Figure 6F,G). Thus, SARS-CoV-2 spike protein appears to facilitate membrane-fusion in a manner that is dependent on palmitoylation of its uniquely cysteine-rich CTD, but through a mechanism unique from canonical membrane nanodomains, although we cannot rule out a discrepancy in lipid raft properties between GPMVs and living cells (Levental et al., 2020).

Despite lack of partitioning into the cholesterol-rich ordered phase of GPMVs, we noted that long-term culture caused spike (but not ACE2) to accumulate in immobile deposits on the glass surface of the culture dish (Figures 6E; S7B). Recent studies determined that cholesterol-rich membrane components are particularly prone to sloughing from the cell and sticking to glass surfaces (He et al., 2018; Hu et al., 2019b). We thus co-expressed the cholesterol-binding protein SCARB1 (Linton et al., 2017) with spike or stained spike cells with fluorescently labeled cholesterol. Both cholesterol markers co-localized with immobilized spike deposits (Figures 6E; S7A-C). Taken together, the data suggests that spike directly associates with a population of cholesterol (Das et al., 2014; Kinnebrew et al., 2019), which is biochemically distinct from the sphingomyelin-associated lipid complexes enriched in canonical rafts.

To interrogate the role of cholesterol in cell fusion, we tested drugs that disrupt cholesterol synthesis (zaragozic acid) or reduce plasma membrane cholesterol (25-hydroxycholesterol; methyl-beta-cyclodextrin or “MBCD”) in the U2OS-ACE2 heterokaryon assay. All compounds inhibited fusion in a dose-dependent manner (Figure 6H). However, such drugs can indirectly lead to cholesterol-independent changes in membrane lipid composition, especially at high concentrations (Zidovetzki and Levitan, 2007). To more directly study the role of cholesterol levels, we harnessed MBCD’s ability to shuttle specific lipids into the plasma membrane (Zidovetzki and Levitan, 2007). Unlike MBCD-conjugated linoleic acid and oleic acid, cholesterol greatly enhanced fusion (Figures 6I; S7D,E).

We surmised that the drug repurposing screen identified compounds that act similarly, thus implicating a counteracting plasma membrane property that increases fusion. Indeed, a small subset of compounds, which include allylamine antifungals (naftifine and terbinafine) and anesthetics (ropivacaine, bupivacaine, propofol), enhance fusion in a dose-dependent manner (Figures 6J; S7F). Whether this is related to an opposing effect on lipid bilayer composition and dynamics relative to cholesterol-depleting drugs requires further inquiry, but is intriguing in light of extensive literature on anesthetics and membrane mobility (Cornell et al., 2017; Goldstein, 1984; Gray et al., 2013; Tsuchiya and Mizogami, 2013).

### SARS2-CoV-2 infection depends on membrane cholesterol of the virus but not the host cell

Our findings on ACE2/spike-mediated fusion, using both U2OS and VeroE6 cells, suggest that many effective compounds prevent fusion by depleting cholesterol from the plasma membrane (Figure 6). While the relevance of such drugs for syncytium formation and disease pathogenesis *in vivo* remains circumstantial (Figure 2), the data nonetheless has implications for virus assembly and entry. Specifically, we predict that such compounds would lack efficacy in virus entry models (Chen et al., 2020; Dittmar et al., 2020; Riva et al., 2020; Wei et al., 2020; Zhu et al., 2020b), instead requiring perturbation of the spike-containing virus membrane derived from the donor cell. To test this, we quantified spike-pseudotyped MLV particle entry into ACE2/TMPRSS2-expressing A549 acceptor cells (Figure 7A), which are primarily infected via the direct fusion pathway (Hoffmann et al., 2020b; Shirato et al., 2018; Zhu et al., 2020b). Apilimod, a PIKFYVE inhibitor and promising therapeutic in multiple SARS-CoV-2 models (Kang et al., 2020; Riva et al., 2020) including heterokaryon assays tested herein (Figures 3E; S3B), prevented pseudoparticle entry at nanomolar concentrations (Figure 7B). By contrast, 25-hydroxycholesterol, which lowers plasma membrane cholesterol by redirection to the cell interior (Abrams et al., 2020; Im et al., 2005; Wang et al., 2020; Yuan et al., 2020; Zhu et al., 2020b; Zu et al., 2020), had no effect (Figure 7C). However, MBCD, which directly “strips” plasma membrane cholesterol without engaging intracellular targets (Zidovetzki and Levitan, 2007), blocked virus entry (Figure 7D).

**Figure 7.**
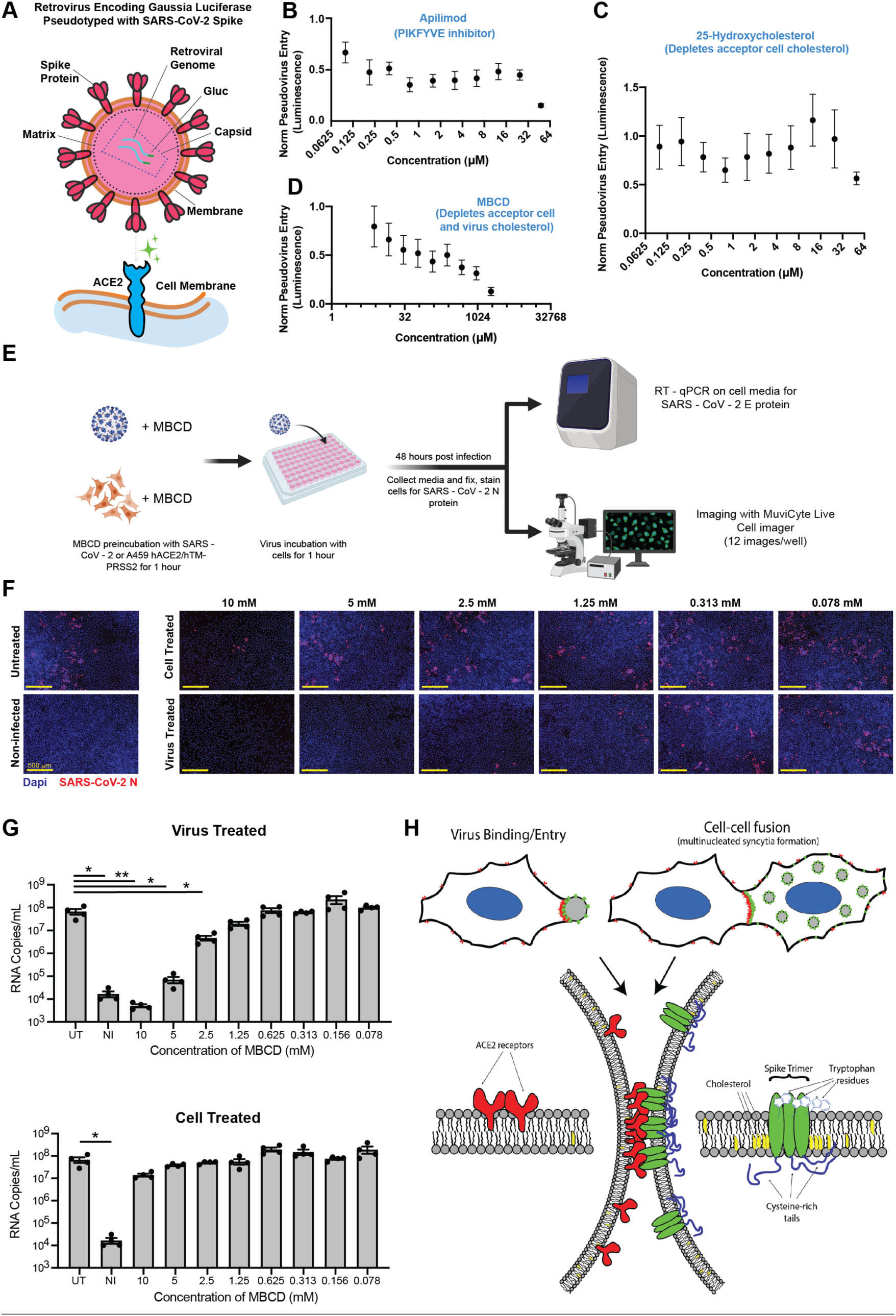
SARS2-CoV-2 infection depends on membrane cholesterol of the virus but not the host cell. **(A)** Schematic representation of pseudotyped virus entry assay in ACE2/TMPRSS2-expressing A549 acceptor cells, which are primarily infected via direct fusion pathway. Pseudovirus encodes *Gaussia* luciferase gene, which allows luminescence-based measure of relative entry as a function of compound concentration. **(B)** Dose-dependent inhibition of pseudovirus entry (luminescence, arbitrary units) for a positive control compound (apilimod, PIKFYVE inhibitor), relative to control (1 = no effect; 0 = complete block). Mean and SEM indicated for six replicates. **(C)** Similar to Figure 7B, but for cholesterol-transport disrupting drug, 25-hydroxycholesterol. **(D)** Similar to Figure 7B, but for plasma membrane cholesterol-stripping compound, MBCD. **(E)** Schematic of SARS-CoV-2 infection assays in ACE2/TMPRSS2 A549 acceptor cells. Relative infection is determined by RT-qPCR or immunohistochemistry of the SARS-CoV-2 nucleocapsid protein (N). **(F)** Representative immunofluorescence (nucleocapsid protein, red; nuclei/DAPI, blue) of A549 cells, 48-hours post-infection by SARS-CoV-2. Top: cells pre-treated with indicated dose of MBCD, followed by wash; bottom: pre-treatment of virus. **(G)** Similar to Figure 7F, but using RT-qPCR to quantify viral titer (RNA copies per mL cell media) following MBCD-treatment of virus (top) or cells (bottom). Identical controls plotted on both graphs for visualization purposes: UT = untreated cells, NI = non-infected cells. Mean and SEM indicated for n=4 independent biological replicates (black dots). P-values of <0.05 and <0.01 are represented by * and **, respectively. **(H)** Graphical model of the biomolecular interactions required for SARS-CoV-2 spike-mediated membrane fusion. Bottom: palmitoylated cysteines (blue) act as multivalent membrane contacts, anchoring trimeric spike peplomers (green) to the phospholipid bilayer (black) and potentially allowing transient higher-order assemblies of trimers. Aromatic residues (e.g. tryptophans) at the spike ectodomain-membrane interface associate with accessible cholesterol (yellow) to promote synapse-like clusters with ACE2 receptors (red) on apposing membranes. Without these collective interactions, spike’s fusion machinery (e.g. fusion peptide and heptad repeats) is unable to surmount the energetically costly barrier to lipid bilayer mixing, both in virus-cell (top, left) and cell-cell fusion (top, right).

Given the relative potency of MBCD in the pseudovirus entry model vs. syncytia assays (Figures 6H; 7D), we hypothesized that it acts by depleting cholesterol from the viral membrane and not the host cell. To discriminate these possibilities, we used the same ACE2/TMPRSS2-expressing A549 cell line, but with a clinical isolate of SARS-CoV-2 virus (Figure 7E). Addition of virus in absence of drug resulted in significant infection of the cell layer, with many cells appearing as multi-nucleated following infection (Figure 7F). Pre-treatment of cells with millimolar doses of MBCD, which strongly inhibits both co-culture syncytia formation and pseudovirus entry, had no effect on infection as determined by RT-PCR and immunohistochemistry (Figure 7F,G). In striking contrast, pre-treatment of SARS-CoV-2 with low micromolar MBCD completely blocked infection (Figure 7F,G). Therefore, cholesterol content of SARS-CoV-2 viral particles, but not the host cell, is critical to infectivity (Figure 7H).

## Discussion

Unprecedented resources devoted to the COVID-19 pandemic have allowed identification of promising stopgap therapeutic measures, as reaching effective vaccination coverage among populations will take time. Virus entry-based assays were particularly critical to discovering essential receptors (ACE2) and proteases for SARS-CoV-2 infection, along with promising repurposed drugs (Dittmar et al., 2020; Hoffmann et al., 2020a; 2020b; Lan et al., 2020; Ou et al., 2020; Riva et al., 2020; Shang et al., 2020; Walls et al., 2020; Wei et al., 2020; Wrapp et al., 2020; Yan et al., 2020; Zhu et al., 2020b). However, many fundamental aspects of the SARS-CoV-2 infectious cycle remain poorly understood, hampering efforts for effective treatment. In particular, commonly used approaches are poorly equipped to interrogate factors that contribute to the formation of a fusion-competent virus, particularly in a high-throughput manner amenable to both small molecule and genetic screens. Here, we show that *in vitro* SARS-CoV-2 spike and ACE2 cell co-culture assays overcome this limitation, and uncover a critical role for viral membrane composition in infection and formation of pathological syncytia.

Our approach relies on a combination of high throughput screening, quantitative live cell imaging, and viral infection assays, all of which implicate cholesterol-rich regions of the plasma membrane in spike-mediated membrane fusion. Cholesterol is known to preferentially interact with certain membrane proteins, particularly those modified with specific lipid moieties (e.g. palmitic acid) (Levental et al., 2010; Martin, 2013; Sobocińska et al., 2017), together clustering into nanodomains which have been referred to as lipid rafts (Levental et al., 2020; Simons and Ikonen, 1997; Veatch and Keller, 2005). In the context of SARS-CoV-2 spike, such cholesterol-rich nanodomains could potentially facilitate the energetically-unfavorable process of lipid bilayer mixing (Heald-Sargent and Gallagher, 2012; Kim and Chen, 2019; Tenchov et al., 2006). However, given its lack of partitioning into ordered GPMV domains (Figure 6D), spike may form nanoscale clusters by a different mechanism. A favored model is that an accessible population of cholesterol, independent of sphingolipids and rafts (Das et al., 2014; Kinnebrew et al., 2019), interacts directly with spike trimers and mediates formation of higher-order protein-lipid assemblies (Figure 7H). Precedent for this model is provided by raft-independent yet cholesterol-dependent mechanisms of biomolecular clustering essential for influenza infection (Goronzy et al., 2018; Zawada et al., 2016). Spike’s cysteine-rich CTD could further amplify this effect, by directly promoting dynamic protein oligomers, similar to what was recently described for the *orthoreovirus* p14 FAST protein (Chan et al., 2020a). The interplay between oligomerization, palmitoylation, cholesterol association, and membrane dynamics will require further inquiry beyond the scope of this study.

In strong support for a key role of lipid structure and composition in spike-mediated membrane fusion, many compound classes over-represented in our drug repurposing screen are implicated in perturbation of the lipid bilayer (e.g. antifungal azoles, rapalog/mTOR inhibitors, ivermectin analogs) (Figure 4C-F) (Bauer et al., 2018; Head et al., 2017; Long et al., 2020; Mast et al., 2013; Trinh et al., 2017; Xu et al., 2010). Interestingly, anti-fungals appear to be enriched in both fusion-inhibiting and - promoting hits. However, the latter tended to favor allylamines rather than azoles, implicating physicochemical and structural differences in direction of effect despite similar action on fungal ergosterol synthesis. We further note that our screen identified anesthetics as promoters of fusion (Figures 4B; 6J). This is intriguing, given that anesthetics are chemically diverse, hydrophobic molecules, which perturb lipid mobility and ordering in membranes (Cornell et al., 2017; Goldstein, 1984; Gray et al., 2013; Tsuchiya and Mizogami, 2013). Whether such similarities can be extended to other fusion-promoting compounds remains to be determined, as is the root membrane-based property that discriminates them from drugs that prevent syncytia formation.

Our high-throughput screening approach uses transcellular membrane fusion as a proxy for virus entry into the host cell. However, beyond a screening tool, we present findings that suggest direct pathological relevance for syncytia. Specifically, human cell populations expressing ACE2 or spike cause cellular pathology with unexpected parallels to COVID-19 patient histology, who feature pervasive multinucleated cells in lung tissue (Figure 2) (Giacca et al., 2020; Tian et al., 2020). These COVID-19 patient syncytia are similar to those found in other respiratory viral infections (Frankel et al., 1996; Johnson et al., 2007; Maudgal and Missotten, 1978). Whether COVID-19 syncytia arise by the cellular pathways described in this work is unclear, although recent data is supportive. For example, contemporary studies report syncytia in cultured cells following exposure to infectious SARS-CoV-2 (Buchrieser et al., 2020; Cattin-Ortolá et al., 2020; Papa et al., 2020; Xia et al., 2020), an observation that we extend to ACE2/TMPRSS2 A549 cells (Figure 7F). Further, the synapses that precede fusion (Figure 1) superficially resemble the virus-filled filopodia observed following SARS-CoV-2 infection (Bouhaddou et al., 2020). We hypothesize that such structures arise via intercellular interactions between ACE2 and virus-unincorporated spike clusters, but cannot rule out the proposed kinase-based mechanism (Bouhaddou et al., 2020).

Whether or not syncytia play a major role in COVID-19 disease progression, the fact that lipid-targeting drugs disrupt spike-mediated membrane fusion has implications for treatment. Indeed, cholesterol-tuned viral infectivity was similarly shown using both spike-pseudotyped MLV and patient isolates of SARS-CoV-2. These data provide strong evidence that viral spike-cholesterol association, and not lipid composition of the host cell membrane or resultant endocytic pathway, mediates effects reported in this study (Figure 7) and perhaps others (Zhu et al., 2020b). Ultimately, viral membranes are produced from the membranes of infected host cells, suggesting that lipid-targeting treatments might disrupt formation of fusion-competent virus particles. Given the advanced state of hyperlipidemia drug development (e.g. statins) (Goldstein and Brown, 2015), insights from this work may be of immediate significance for COVID-19 treatment. Consistent with this, retrospective analyses of patient outcomes observed significant reductions in mortality for those prescribed cholesterol-lowering statins (Daniels et al., 2020; Zhang et al., 2020). A critical role for cholesterol in virus assembly may explain this observation, and could partially account for why COVID-19 risk factors (e.g. menopause, obesity, age) (Costeira et al., 2020; Zhou et al., 2020) similarly correlate with differential sterol processing. Additional work is clearly required, but in the context of the rapidly evolving landscape of COVID-19 treatment options, our findings underscore the potential utility of statins and other lipid modifying treatments.

Invariably, opportunistic infections hijack physiological cellular processes to ensure their survival (Pelkmans and Helenius, 2003). To this end, we speculate that physiological and pathological synapses and resulting syncytia (or lack there of) arise in part from shared lipid bilayer properties at the nanoscale. Consistent with this concept, actin-dependent, ACE2/spike fusion events proceed from “finger-like” projections and synapses between cells to fusion pore dilation and membrane collapse, closely resembling the orderly biogenesis of myoblast-derived syncytia (Chen et al., 2008; Duan et al., 2018; Kim and Chen, 2019; Shi et al., 2017; Shilagardi et al., 2013). Second, we highlight the remarkable similarities between SARS-CoV-2 spike and tricellular tight junction proteins (Higashi et al., 2013; Oda et al., 2020; Sohet et al., 2015), specifically with respect to membrane-anchoring cysteines and aromatics (Figures 5H; S6D,E). These observations suggest that both pathological viruses and adherent cells independently evolved proteins with abnormally strong affinity for the plasma membrane to ensure stability of transcellular complexes (Figure 7H), whether to initiate fusion or maintain tissue integrity. We thus envision that assays presented herein may have broad utility for understanding the biophysics of synapse and fusion pore assembly, representing an exciting example of how inquiry into viral pathogenesis illuminates physiological function.

## Supporting information

TableS1_Compund_Screen

TableS2_Cys-Rich_Viral_TM_Proteins

TableS3_Human_Spike-Like_Proteins

TableS4_TM_Domains_Compounds

VideoS1_Synapses

VideoS2_Synapse_and_Fuse

VideoS3_BuildingASyncytium

## Acknowledgments

We thank all Brangwynne Lab members for helpful discussion and critiques and Evangelos Gatzogiannis for help with live cell microscopy. A.D. wishes to thank the Hargrove lab at Duke University, and particularly Sarah Wicks, for assistance and use of the ChemAxon analysis software, as well as Dr. Brittany Morgan for helpful discussions. This work was supported by Princeton COVID-19 research funds through the Office of the Dean for Research (C.P.B. and A.P. labs); the Howard Hughes Medical Institute (C.P.B. lab); a Boston University start-up fund and Peter Paul Career Development Professorship (F.D.); NIH (GM095467 and HL122531 to B.D.L.; GM134949, GM124072, and GM120351 to I.L.); Volkswagen Foundation (I.L.); Human Frontiers Science Program (I.L.); a Burroughs Wellcome Fund Award for Investigators in Pathogenesis (A.P.); Longer Life Foundation—RGA/Washington University Collaboration (A.S.H.); postdoctoral fellowship awards from the Uehara Memorial Foundation and JSPS Research Fellowships for Young Scientists (T.T.); from the SENSHIN Medical Research Foundation (S.S).

## Author Contributions

Conceptualization, D.W.S., C.C.J, P.J.A., D.B., H.K., C.P.B.; Methodology, D.W.S., C.C.J, P.J.A., D.B., A.D., D.K., I.C.-S., S.S., T.T., A.S.H., R.F.P.; Investigation—cell culture studies (non-virus), D.W.S., C.C.J, P.J.A., D.B., A.D., H.K., I.C.-S; Investigation—informatics, D.W.S., P.J.A., A.D., A.S.H.; Investigation—virus studies, D.K., S.S., T.T.; Investigation—human pathology, R.F.P.; Resources, A.H.T., M.S.; Writing—Original Draft, D.W.S. and C.P.B.; Writing—Review & Editing, all authors; Funding Acquisition, C.P.B., B.D.L., F.D., I.L., A.P.; Supervision, C.P.B., B.D.L., F.D., I.L., A.P.

## Declaration of Interests

C.P.B. is a scientific founder and consultant for Nereid Therapeutics. A.S.H. is a consultant for Dewpoint Therapeutics.

## Figure Legends

**Figure S1, Related to Figure 1.**
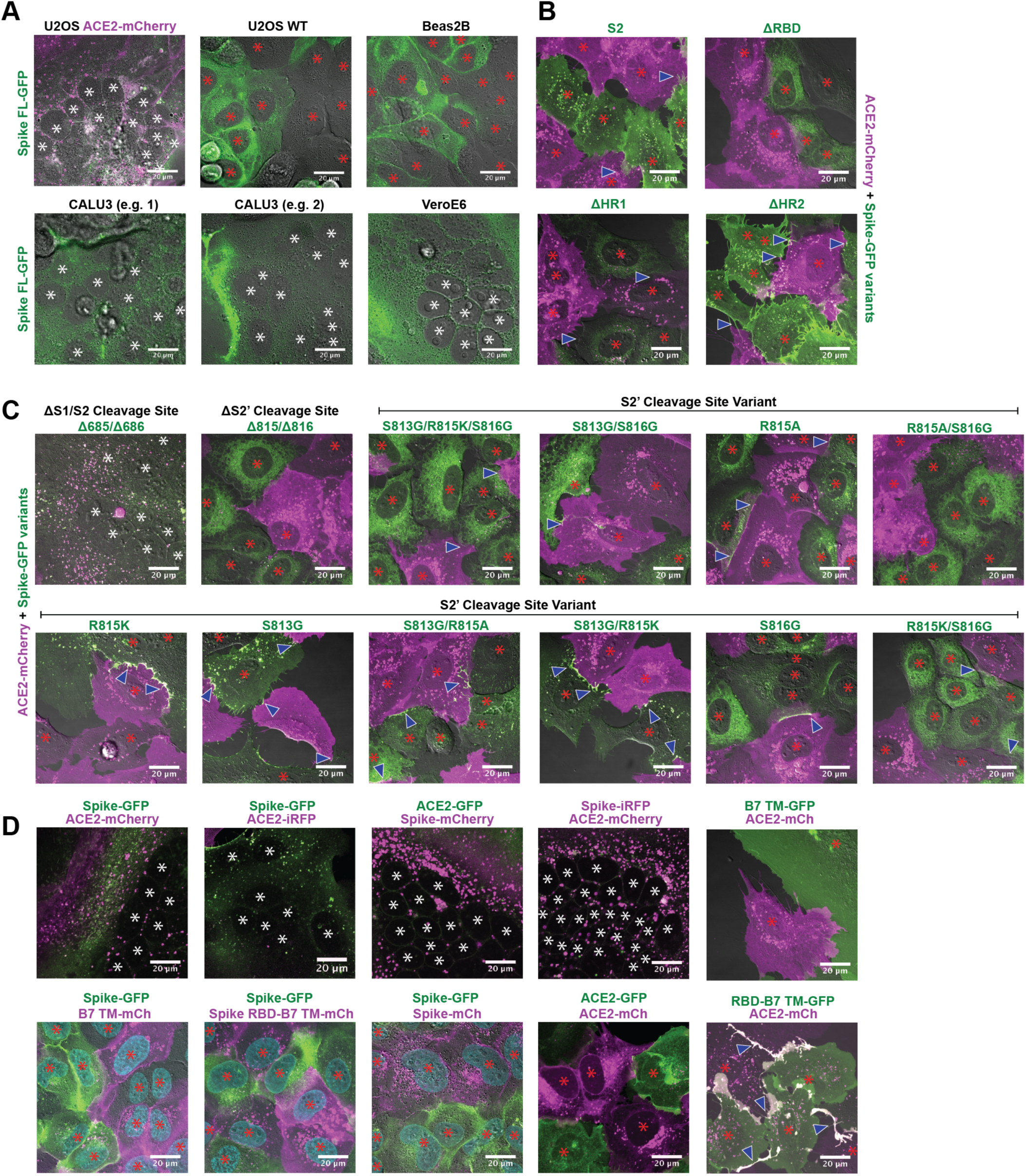
Syncytia derive from fusion events at synapse-like, spike-ACE2 protein clusters. **(A)** Indicated non-transduced cells (or ACE2-mCherry/U2OS control) co-cultured with U2OS spike-GFP (green) cells for 24-hours. White asterisks indicate nuclei in syncytia; red, in isolation. **(B)** Indicated GFP-spike variant (green) U2OS cells co-cultured with U2OS ACE2-mCherry (magenta) cells for 24-hours. White asterisks indicate nuclei in syncytia; red, in isolation; arrowhead, synapses (select examples noted). **(C)** Similar to Figure S1B, but using spike variants that disrupt its two cleavage sites (S1/S2 vs. S2’). **(D)** U2OS Cells expressing spike or ACE2 with indicated fluorescent tag, co-cultured for 24-hours. White asterisks indicate nuclei in syncytia; red, in isolation; arrowhead, synapses (select examples noted). **(E)** Similar to Figure S1D but single fluorescence channels and spike RBD-B7 TM-GFP donor cells. **(F)** Similar to Figure S1E but full-length spike-GFP donor cells.

**Figure S2, Related to Figure 3.**
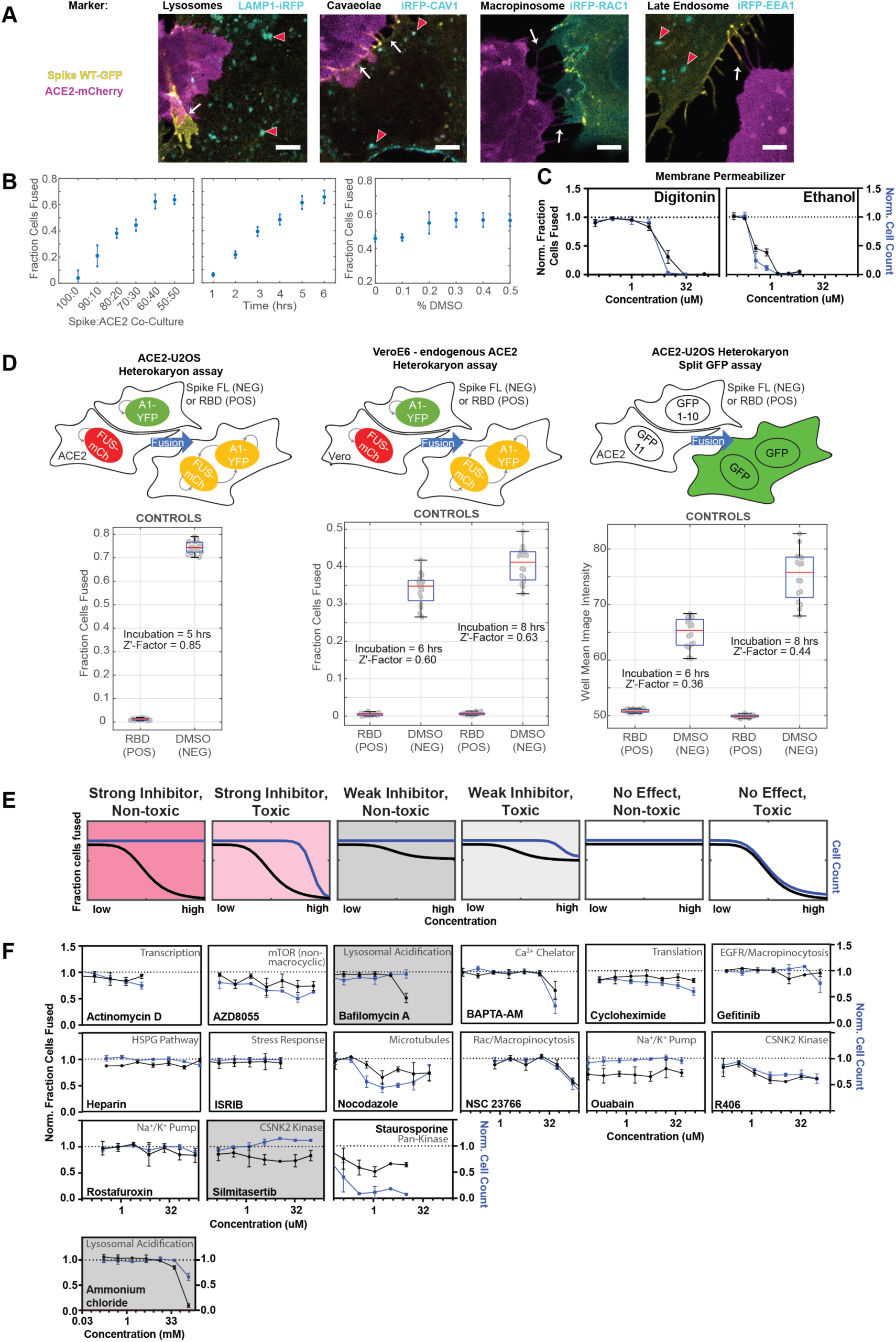
A novel high-throughput screening platform identifies modulators of syncytia formation. **(A)** U2OS cells co-expressing spike-GFP (yellow) and iRFP-tagged (cyan) endocytic markers (indicated) co-cultured with ACE2-mCherry (magenta) cells, with cell-cell fusion events assessed by live cell microscopy. Representative images show lack of co-localization between spike and iRFP-tagged proteins at synapses prior to fusion. Scale-bar, 5 µm. **(B)** Optimization of ACE2-U2OS heterokaryon assay. Co-cultures were performed and fusion was quantified upon altering: spike to ACE2 cell ratio (left); length of time prior to fixation (middle); DMSO concentration (right). Default experimental parameters: 1-to-1 ratio, 5-hours, 0.5% DMSO. Mean and SEM: n=4 biological replicates (16 images per). **(C)** Dose-response for membrane-permeabilizing compounds in ACE2-U2OS heterokaryon assay. Cell count (blue) and fraction fused (black) curves are similar, indicating lack of toxicity-independent inhibition of cell-cell fusion. Mean and SEM: n=4 biological replicates (16 images per). **(D)** Comparison of three different heterokaryon microscopy assays, each of which use fluorescent proteins as readouts for fusion. Bottom: relative fusion for positive (spike RBD donor cells; fusion-incompetent) and negative (spike full-length donor cells + DMSO; fusion-competent) controls for indicated assay (top, with graphical depiction) with measured Z’-factor, which indicates assay strength (higher = more robust). Mean and SEM: n=4 biological replicates (16 images per). **(E)** Schematic of ACE2-U2OS assay dose-response and interpretation. Fraction of cells fused (black curve) relative to cell count (toxicity measure; blue curve), both normalized by plate negative control, indicates compound efficacy (pink, strong inhibitor; gray, weak inhibitor; white, no-effect). **(F)** Dose-response relationships for additional compounds tested in ACE2-U2OS heterokaryon assay. Mean and SEM: n=4 biological replicates (16 images per). See Figure S3 for same compounds in VeroE6 heterokaryon assay.

**Figure S3, Related to Figure 3.**
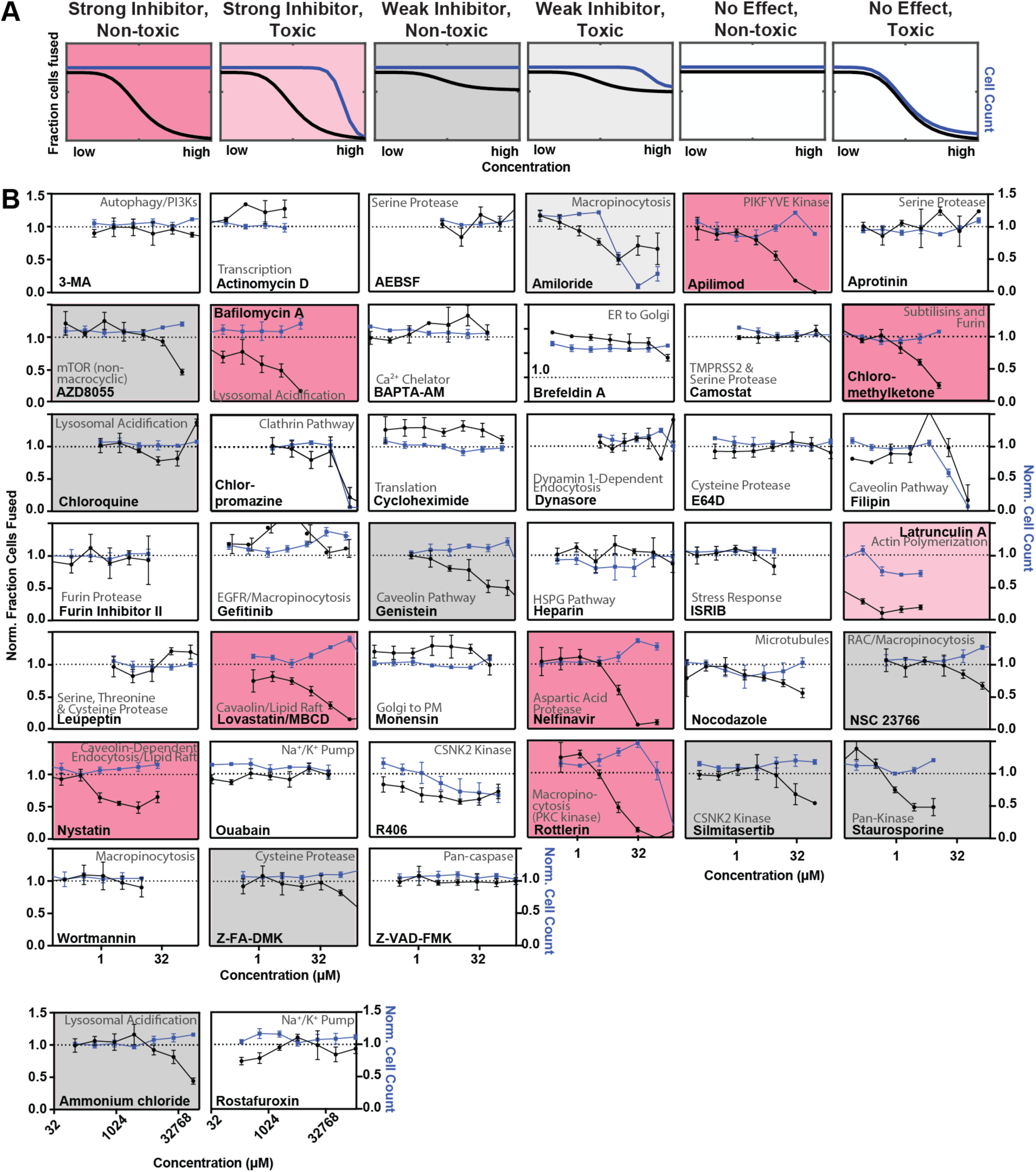
A novel high-throughput screening platform identifies modulators of syncytia formation. **(A)** Schematic of VeroE6 heterokaryon assay (no exogenous ACE2 overexpression) dose-response and interpretation. Fraction of cells fused (black curve) relative to cell count (toxicity measure; blue curve), both normalized by plate negative control, indicates compound efficacy (pink, strong inhibitor; gray, weak inhibitor; white, no-effect). **(B)** Dose-response relationships for all compounds (n=41) tested in VeroE6 heterokaryon assay. Mean and SEM: n=4 biological replicates (16 images per). See Figures 2 and S2 for effect of same compounds in ACE2-U2OS heterokaryon assay.

**Figure S4, Related to Figure 4.**
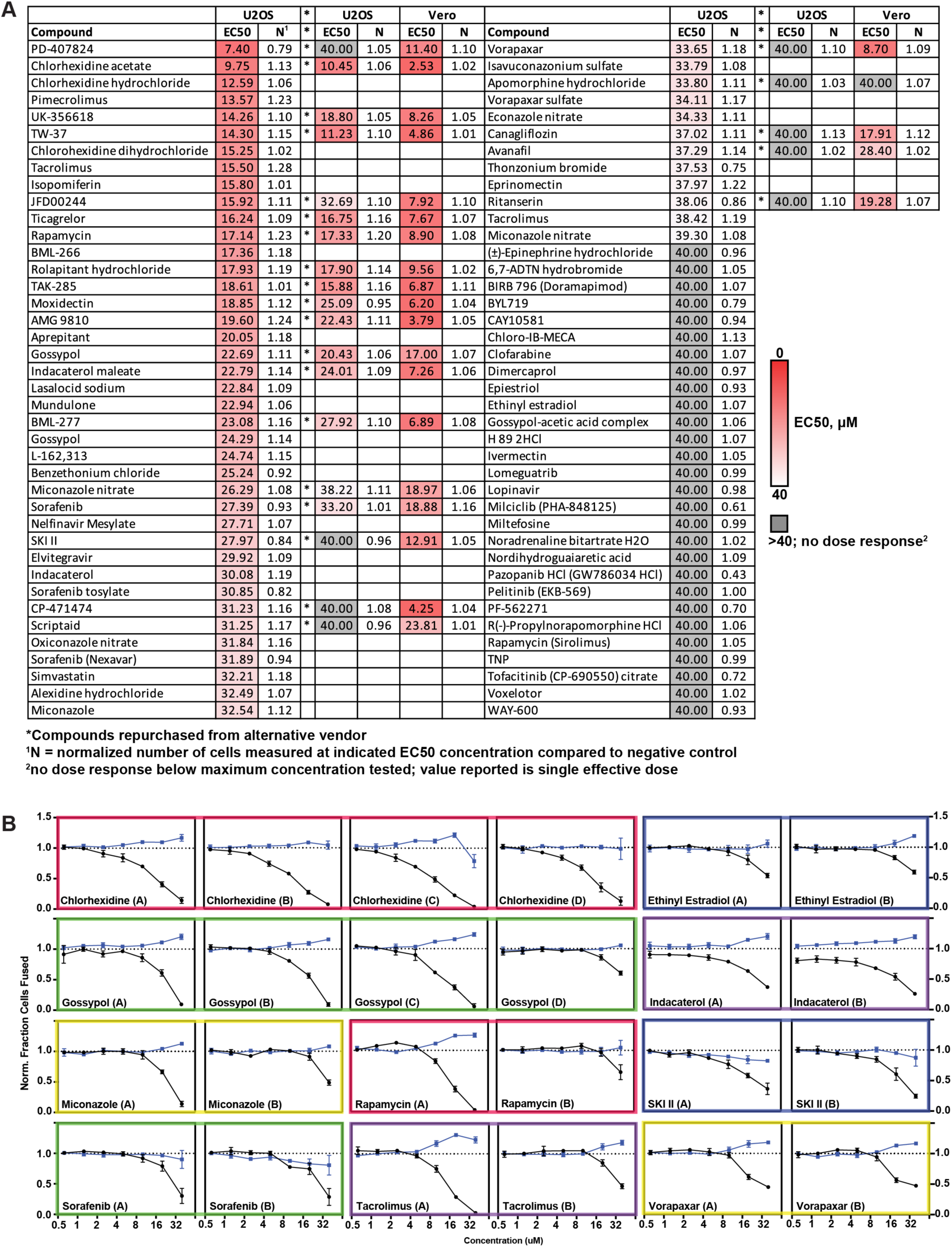
A drug repurposing screen implicates membrane lipid composition in cell-cell fusion. **(A)** Summary of 7-point dose-response for top-80 inhibitors in indicated heterokaryon assay (ACE2-U2OS vs. VeroE6). Asterisks reference compounds validated by re-purchase from independent manufacturers. Heat-map indicates relative compound potency (bright red = higher). **(B)** Dose-response relationships for compounds (grouped by color) present in multiple small molecule libraries, each of whose replicates were identified as top-80 inhibitory hits. Mean and SEM: n=3 biological replicates (16 images per).

**Figure S5, Related to Figure 4.**
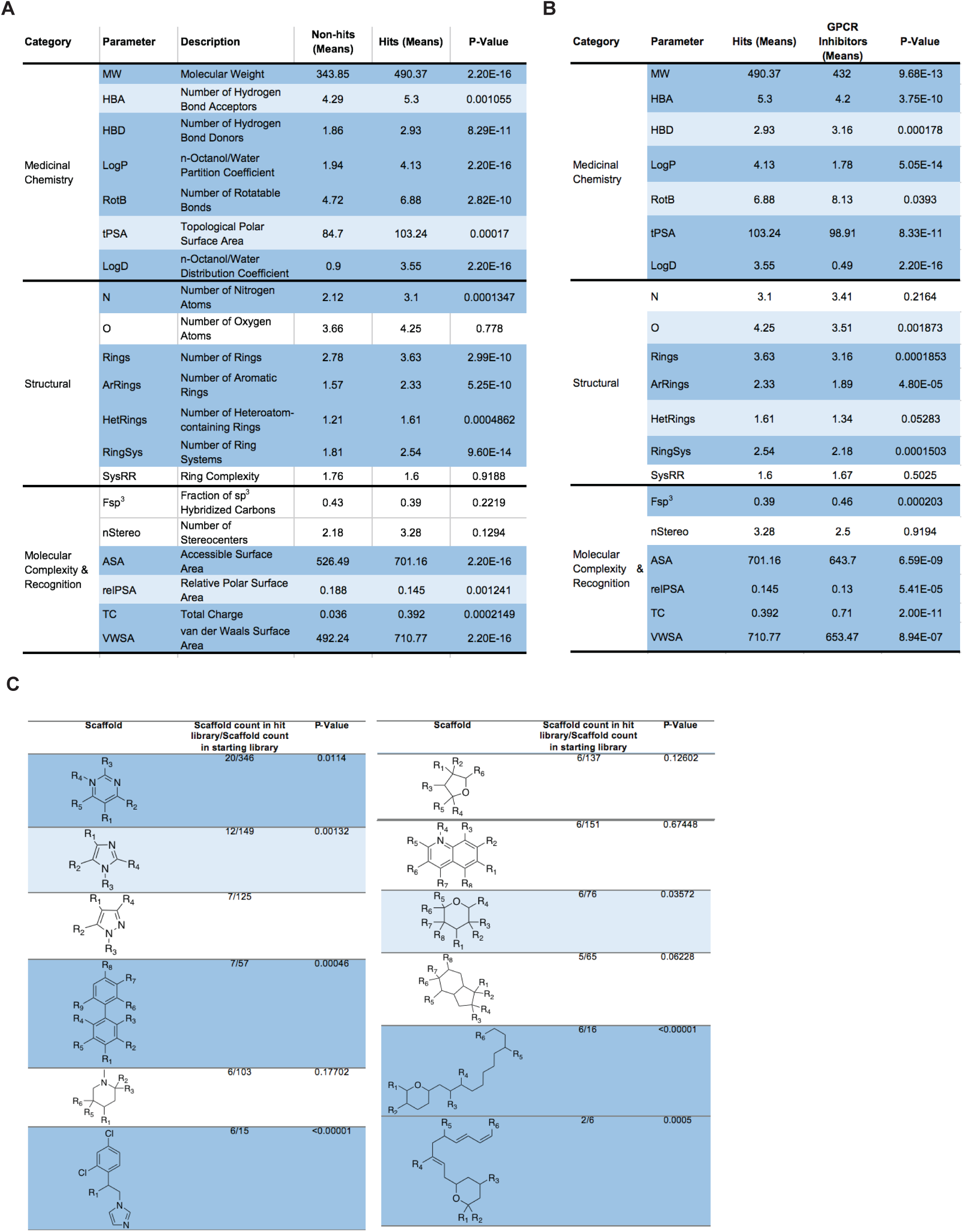
A drug repurposing screen implicates membrane lipid composition in cell-cell fusion. **(A)** Analysis and comparison of 20 physicochemical properties for non-hit and hit libraries. P-values assessed with a Mann-Whitney U test. Dark and light blue represent P-values <0.001 and <0.05, respectively. White indicates no significant differences. **(B)** Similar to Figure S5A, but inhibitory hits vs. a control GPCR inhibitor library obtained from literature. **(C)** List of top-10 most frequent scaffolds and substructures in the hit library as analyzed by Scaffold Hopper and RDKit Substructure Search, respectively. Enrichment significance relative to the starting library was assessed with two-tailed Z-scores used to calculate P-values. Dark and light blue represent P-values <0.001 and <0.05, respectively. White indicates no significant differences.

**Figure S6, Related to Figure 5.**
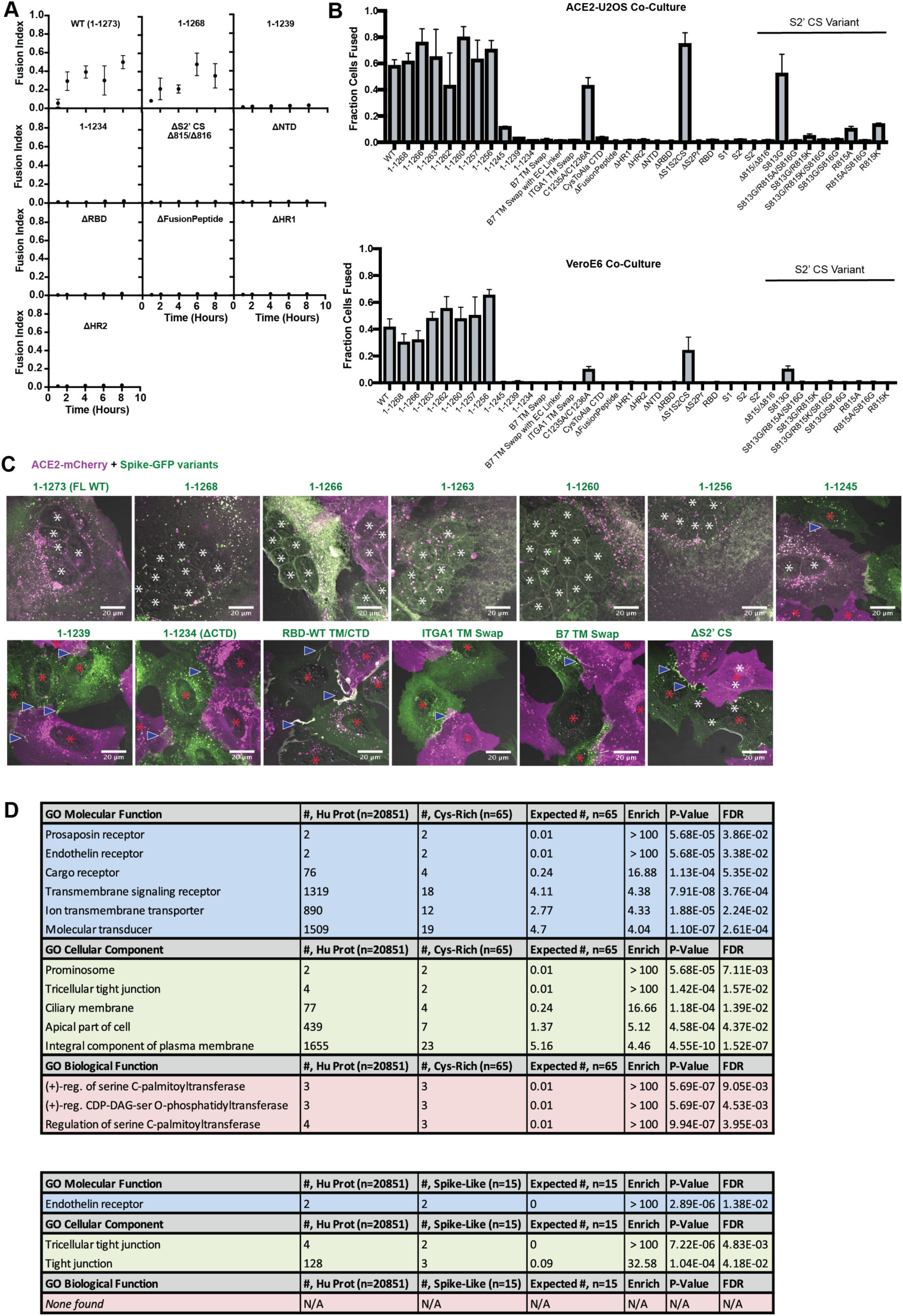
Highly unusual membrane-proximal regions of spike are needed for fusion. **(A)** ACE2-U2OS heterokaryon assay, but assesses kinetics of fusion by varying length of co-culture with spike variant (indicated)-expressing HNRNPA1-EYFP cells. Mean and SEM: n=4 biological replicates (16 images per). **(B)** Similar to Figure S6A but for all studied spike variants at a single time-point (5-hours) and examined in both ACE2-U2OS (top) and VeroE6 (bottom) heterokaryon assays. **(C)** Representative images of ACE2-mCherry (magenta) cells co-cultured for 24-hours with U2OS cells expressing GFP-tagged spike variant (green, labeled). White asterisks indicate nuclei in syncytia; red, in isolation; arrowhead, synapses between cells (select examples noted). **(D)** Top: Gene ontology (PantherDB) for cysteine-rich human transmembrane (TM) proteins, assessing significance of enrichment relative to the entire human proteome. Bottom: similar but for “spike-like” human proteins (aromatics at TM interface, high cysteine content in cytoplasm CTD).

**Figure S7, Related to Figure 6.**
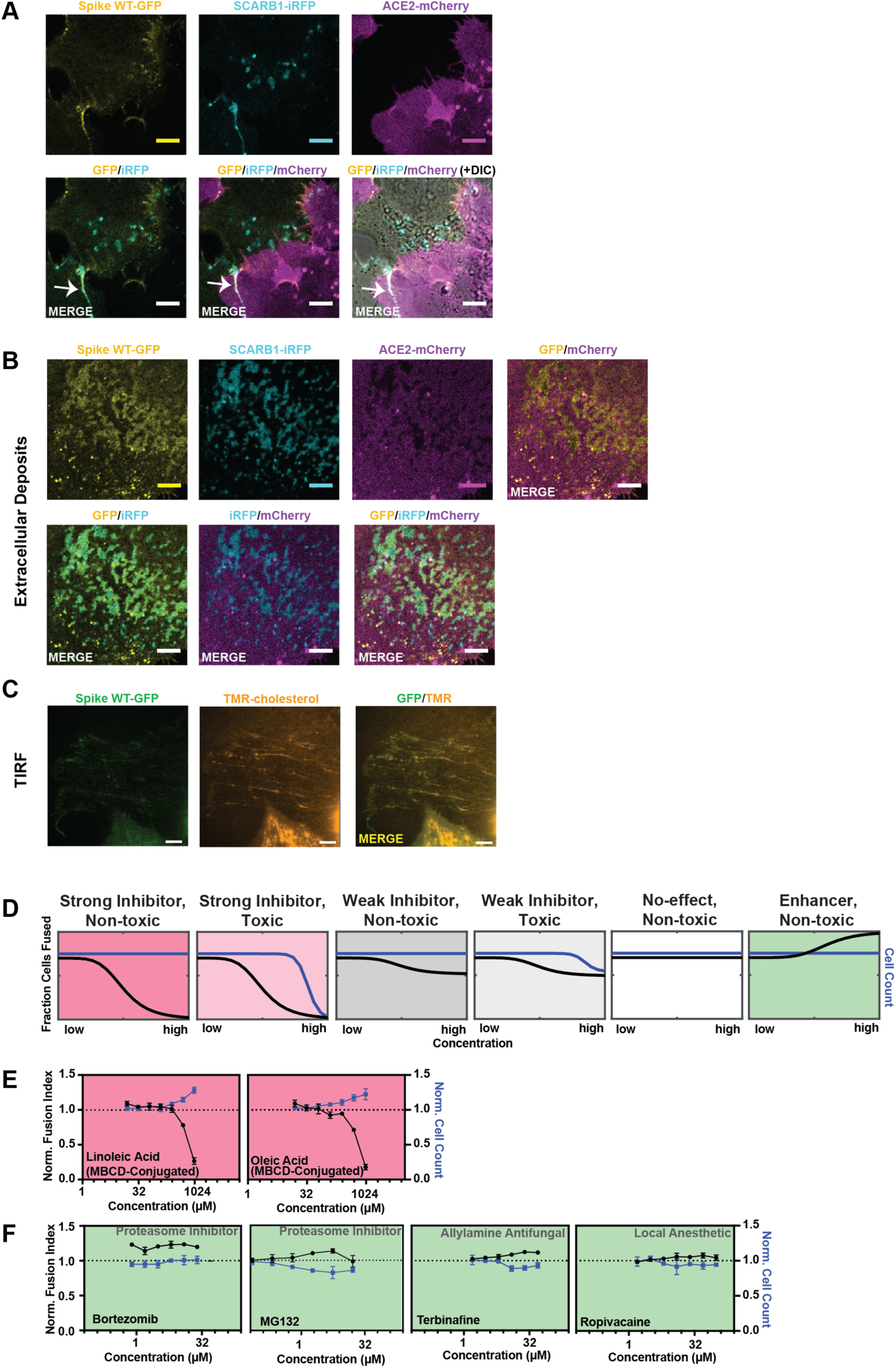
spike requires membrane cholesterol for fusion via a raft-independent mechanism. **(A)** Individual fluorescence channels for merge image shown in Figure 6E **(top)**. Arrow indicates co-localization between cholesterol-binding protein SCARB1-iRFP (cyan) and spike-GFP (yellow) at transcellular synapses. **(B)** Individual fluorescence channels for merge image shown in Figure 6E **(top)**. **(C)** Representative images of extracellular membrane deposits that contain both cholesterol (CholEsteryl-BODIPY stain, orange) and spike-GFP (green). **(D)** Graphical schematic for ACE2-U2OS assay dose-response and interpretation. Fraction of cells fused (black curve) relative to cell count (blue curve), both normalized by the plate negative control, indicates compound effectiveness (pink, strong inhibitor; gray, weak inhibitor; white, no-effect; green, enhancer). **(E)** Dose-response analysis of control MBCD-conjugated lipids in ACE2-U2OS heterokaryon assay. Black line indicates fraction of cells fused; blue, cell count (toxicity measure); both of which are normalized to DMSO negative control. Mean and SEM: n=4 biological replicates (16 images per). **(F)** Similar to Figure S7E, but with compounds predicted by drug repurposing screen to enhance fusion. Drug class indicated.

## The online supplement contains three videos and four tables

**Video S1, Related to** Figure 1**. Transcellular ACE2-spike synapses are long-lived cellular assemblies.** ACE2-mCherry, magenta; Spike-GFP, green. 20-minute time lapse, human U2OS osteosarcoma cells stably expressing indicated proteins. Time since start of imaging is indicated.

**Video S2, Related to** Figure 1**. ACE2-spike synapse formation and cell-cell fusion following co-culture.** ACE2-mCherry, magenta; Spike-GFP, green. 2-hour time lapse, human U2OS osteosarcoma cells stably expressing indicated proteins. Time since plating of spike-GFP cells is indicated.

**Video S3, Related to** Figure 1**. Building a syncytium: multiple cell-cell fusion events following addition of a single ACE2 cell to a spike cell monolayer.** ACE2-mCherry, magenta; Spike-GFP, green. 5-hour time lapse, human U2OS osteosarcoma cells stably expressing indicated proteins. Time since plating of ACE2-mCherry cell is indicated.

**Table S1, Related to** Figure 4. Raw data from unbiased drug repurposing screen.

**Table S2, Related to** Figure 5**. Viral transmembrane proteins with proximal cysteine-rich regions.**

**Table S3, Related to Figures 5 and S6. Human transmembrane proteins with spike-like membrane proximal regions.**

**Table S4, Related to Figures 3-7. Amino acid composition of transmembrane proteins used in this study and cellular targets associated with extensively tested compounds.**

## STAR METHODS

### KEY RESOURCES TABLE

| REAGENT or RESOURCE | SOURCE | IDENTIFIER |
| --- | --- | --- |
| **Antibodies** |  |  |
| Mouse anti-TTF-1 (clone 8G7G3/1) | Agilent Dako | IR05661-2 |
| Rabbit anti-SARS-CoV nucleocapsid (N) protein | Rockland | 200-401-A50 |
| Alexafluor 568 goat anti-rabbit IgG | Thermo Fisher Scientific | A11011 |
| **Virus Strains** |  |  |
| 2019-nCoV (SARS-CoV-2)/USA-WA1/2020 | Center for Disease Control and Prevention and BEI Resources | MN985325 |
| **Biological Samples** |  |  |
| Formalin-fixed, paraffin-embedded, autopsy lung tissue (39 deceased patients or “decedents”) | Brigham and Women’s Hospital, Autopsy Division | N/A |
| **Chemicals, Peptides, and Recombinant Proteins** |  |  |
| Dulbecco’s Modified Eagle Medium (DMEM), High Glucose, Pyruvate | Thermo Fisher Scientific | 11995065 |
| Eagle’s Minimum Essential Media (EMEM) | ATCC | ATCC 30-2003 |
| Opti-MEM Reduced Serum Medium | Thermo Fisher Scientific | 31985062 |
| Penicillin-Streptomycin (10,000 U/mL) | Thermo Fisher Scientific | 15140122 |
| Fetal Bovine Serum, Heat Inactivated | Atlanta Biologicals | S11150H |
| In-Fusion HD Cloning Plus | Takara Bio | 638910 |
| Lipofectamine 3000 Transfection Reagent | Thermo Fisher Scientific | L3000008 |
| Polybrene | Sigma-Aldrich | TR-1003-G |
| Paraformaldehyde (16%) | Electron Microscopy Services | 15710 |
| Phusion High-Fidelity DNA Polymerase | New England Biolabs | M0530L |
| Texas Red DHPE | Thermo Fisher Scientific | T1395MP |
| Annexin V 647 | Thermo Fisher Scientific | A13204 |
| Small molecule library: LOPAC-1280 (1278 bioactive compounds, 96-well) | Sigma-Aldrich | LO1280-1KT |
| Small molecule library: LOPAC-Pfizer (90 Pfizer-developed drugs, 96-well) | Sigma-Aldrich | LO5100 |
| Small molecule library: L1200 (355 kinase inhibitors, 96-well) | Selleck Chemicals | L1200 |
| Small molecule library: L1900 (120 epigenetic-targeting compounds, 96-well) | Selleck Chemicals | L1900 |
| Small molecule library: L2300 (378 cancer-targeting compounds, 96-well) | Selleck Chemicals | L2300 |
| Small molecule library: L4200 (1364 FDA-approved drugs, 96-well) | TargetMol | L4200 |
| Small molecule library: Spectrum collection (2400 bioactive compounds, 96-well) | Microsource Discovery | N/A |
| Nelfinavir mesylate | R&D Systems | 3766/10 |
| Heparin | Sigma-Aldrich | H3393-100KU |
| Nocodazole | Sigma-Aldrich | 487928-10MG |
| Amiloride/EIPA | Sigma-Aldrich | A3085-25MG |
| Wortmannin | Sigma-Aldrich | W1628-1MG |
| Chlorpromazine | Sigma-Aldrich | 215921-500MG |
| Filipin | Sigma-Aldrich | F4767-1MG |
| Nystatin | Thermo Fisher Scientific | BP29495 |
| Leupeptin | Sigma-Aldrich | L2884-5MG |
| AEBSF | Sigma-Aldrich | A8456-25MG |
| Furin inhibitor II (polyarginine) | Sigma-Aldrich | 344931-1MG-M |
| Decanoyl-RVKR-CMK (chloromethylketone) | Tocris | Cat. No. 3501 |
| Methyl-Beta-cyclodextrin (MBCD) | Sigma-Aldrich | C4555-1G |
| Lovastatin | Sigma-Aldrich | 438185-25MG |
| Camostat mesylate | Sigma-Aldrich | SML0057-10MG |
| E-64D (EST) | Sigma-Aldrich | 330005-1MG |
| Ammonium chloride | Sigma-Aldrich | A9434-500G |
| Chloroquine diphosphate | Sigma-Aldrich | C6628 |
| Gefitinib | Sigma-Aldrich | SML1657-10MG |
| BAPTA-AM | Sigma-Aldrich | 196419-25MG |
| Latrunculin A | Invitrogen | 428026-50UG |
| Rottlerin | Sigma-Aldrich | 557370-10MG |
| Dynasore | Sigma-Aldrich | 324410-10MG |
| Sodium chlorate | Sigma-Aldrich | 403016-100G |
| Protease inhibitor cocktail | Sigma-Aldrich | P1860-1ML |
| Ouabain | Sigma-Aldrich | 4995-1GM |
| Rostafuroxin/PST2238 | Sigma-Aldrich | SML1139-5MG |
| Silmitasertib | Selleck Chemicals | Catalog No. S2248 |
| R-406 | Selleck Chemicals | Catalog No. S2194 |
| Apilimod | Selleck Chemicals | Catalog No. S6414 |
| Bafilomycin A | Enzo | BML-CM110-0100 |
| 2-bromopalmitic acid (2-BP) | Sigma-Aldrich | 238422-10G |
| MG132 | Sigma-Aldrich | M8699 |
| Monensin | Sigma-Aldrich | M5273-500MG |
| Brefeldin A | Sigma-Aldrich | B5936-200UL |
| Cycloheximide | Sigma-Aldrich | C4859-1ML |
| Dimethyl sulfoxide (DMSO) | Sigma-Aldrich | D2650-100ML |
| Actinomycin D | Sigma-Aldrich | A5156-1VL |
| Staurosporine | AbCam | ab120056 |
| ISRIB | Sigma-Aldrich | SML0843-5MG |
| 3-methyladenine | Sigma-Aldrich | M9281 |
| Z-FA-FMK | Selleck Chemicals | Catalog No.S7391 |
| NSC 23766 | Selleck Chemicals | Catalog No.S8031 |
| Z-VAD-FMK | Selleck Chemicals | Catalog No.S7023 |
| AZD8055 | Selleck Chemicals | Catalog No.S1555 |
| 25-hydroxycholesterol | Sigma-Aldrich | H1015-10MG |
| Aprotinin (BPTI) | Selleck Chemicals | Catalog No.S7377 |
| Genistein | Selleck Chemicals | Catalog No.S1342 |
| Oleic acid (MBCD-conjugated) | Sigma-Aldrich | O1257-10MG |
| Cholesterol (MBCD-conjugated) | Sigma-Aldrich | C4951-30MG |
| Linoleic acid (MBCD-conjugated) | Sigma-Aldrich | L5900-10MG |
| Myriocin | Sigma-Aldrich | M1177-5MG |
| Zaragozic Acid A | Cayman | Item No. 17452 |
| Digitonin | Sigma-Aldrich | D141-100MG |
| Bortezomib | Selleck Chemicals | S1013 |
| Bupivacaine hydrochloride | Sigma-Aldrich | PHR1128-1G |
| Ropivacaine hydrochloride | Selleck Chemicals | S4058 |
| Naftifine hydrochloride | Cayman | 19234 |
| Terbinafine hydrochloride | Cayman | 10011619 |
| Propofol | Sigma-Aldrich | P-076-1ML |
| Puromycin | Sigma-Aldrich | P7255 |
| Blasticidin S | VWR | 1859-25 |
| DAPI | Sigma-Aldrich | D9542 |
| Hoechst 33342 Solution | Thermo Fisher Scientific | 62249 |
| AMG 9810 | Cayman | 14715 |
| Apomorphine hydrochloride | ApexBio | B6936 |
| Avanafil | Cayman | 23024 |
| BML-277 | Cayman | 17552 |
| Canagliflozin | Cayman | 11575 |
| Chlorohexidine | Targetmol | T1147 |
| CP-471474 | Cayman | 29442 |
| GOSSYPOL, GOSSYPOL, GOSSYPOL! | ApexBio | N2135 |
| Indacaterol maleate | ApexBio | B1369 |
| JFD00244 | Cayman | 14648 |
| Miconazole nitrate | Cayman | 15420 |
| Moxidectin | ApexBio | B3611 |
| PD-407824 | Cayman | 25989 |
| Rapamycin | Targetmol | T1537 |
| Ritanserin | Tocris | 1955 |
| Rolapitant hydrochloride | Targetmol | T3724 |
| Scriptaid | ApexBio | A4106 |
| SKI II | Cayman | 10009222 |
| Sorafenib | Cayman | 10009644 |
| TAK-285 | MCE | HY-15196 |
| Ticagrelor | Cayman | 15425 |
| TW-37 | Cayman | 20999 |
| UK-356618 | ApexBio | A4440 |
| Vorapaxar | ApexBio | A8809 |
| CholEsteryl BODIPY (FL C12) | Thermo Fisher Scientific | C3927MP |
| Fibronectin from bovine plasma | Sigma-Aldrich | F1141-5MG |
| Experimental Models: Cell Lines |  |  |
| Human: HEK293T | Marc Diamond, UTSW | N/A |
| Human: U2OS | Tom Muir, Princeton | N/A |
| Human: Beas2B | Celeste Nelson, Princeton | N/A |
| Human: A549 (ACE2/TMPRSS2) | This study | N/A |
| Human: Calu3 | ATCC | ATCC HTB-55 |
| Monkey: VeroE6 | ATCC | ATCC CRL-1586 |
| Human: U2OS, various | This study | N/A |
| Monkey: VeroE6, various | This study | N/A |
| Recombinant DNA |  |  |
| pDUAL-SCARB1-GFP | Addgene | Plasmid #86979 |
| GFP1-10 | Addgene | Plasmid #80409 |
| GFP11x7-mCherry-a-tubulin | Addgene | Plasmid #70218 |
| pLAMP1-mCherry | Addgene | Plasmid #45147 |
| mCherry-CAV1 | Addgene | Plasmid #55008 |
| GFP-EEA1 | Addgene | Plasmid #42307 |
| pcDNA3.1-SARS-CoV-1-Spike | Addgene | Plasmid #145031 |
| RAC1 (H. sapiens) in pANT7_cGST | DNASU | HsCD00632727 |
| ACE2 cDNA in pcDNA3.1 | Genscript | OHu20260 |
| SARS-CoV-2 (2019-nCoV) Spike ORF (Codon Optimized) | Sino Biological | VG40589-UT |
| pCAGGS-SARS-CoV2-S | BEI Resources, NIH | N/A |
| pMLV gag-pol | Ploss Lab, Princeton | N/A |
| pLMN8-Gluc | This study | N/A |
| p-LAT Transmembrane-mRFP | Levental Lab, Virginia | N/A |
| PSP (lentivirus packaging plasmid) | Marc Diamond, UTSW | N/A |
| VSVG (lentivirus packaging plasmid) | Marc Diamond, UTSW | N/A |
| FM5 lentiviral vector (Ubiquitin C promoter) | Sanders et al., *Neuron*, 2014 | N/A |
| FM5-mGFP-Standardized glycine-serine (GS) Linker-Ascl Site-STOP | Sanders et al., *Cell*, 2020 | N/A |
| FM5-mCherry-Standardized GS Linker-Ascl Site-STOP | Sanders et al., *Cell*, 2020 | N/A |
| FM5-miRFP670-Standardized GS Linker-Ascl Site-STOP | Sanders et al., *Cell*, 2020 | N/A |
| FM5-NheI Site-Standardized GS Linker-mGFP | Sanders et al., *Cell*, 2020 | N/A |
| FM5-NheI Site-Standardized GS Linker-mCherry | Sanders et al., *Cell*, 2020 | N/A |
| FM5-NheI Site-Standardized GS Linker-miRFP670 | Sanders et al., *Cell*, 2020 | N/A |
| FM5-NheI Site-Standardized GS Linker-EYFP | This study | N/A |
| FM5-HNRNPA1-EYFP | This study | N/A |
| FM5-FUS-mCherry | This study | N/A |
| FM5-GFP1-10 | This study | N/A |
| FM5-GFP11x7 | This study | N/A |
| FM5-mGFP-EEA1 | This study | N/A |
| FM5-miRFP670-EEA1 | This study | N/A |
| FM5-mGFP-RAC1 | This study | N/A |
| FM5-miRFP670-RAC1 | This study | N/A |
| FM5-mGFP-CAV1 | This study | N/A |
| FM5-miRFP670-CAV1 | This study | N/A |
| FM5-LAMP1-mGFP | This study | N/A |
| FM5-LAMP1- miRFP670 | This study | N/A |
| FM5-SCARB1-mGFP | This study | N/A |
| FM5-SCARB1-miRFP670 | This study | N/A |
| FM5-MYMK-mGFP | This study | N/A |
| FM5-MYMK-mCherry | This study | N/A |
| FM5-MYMK-miRFP670 | This study | N/A |
| FM5-MYMX-mGFP | This study | N/A |
| FM5-MYMX-mCherry | This study | N/A |
| FM5-MYMX-miRFP670 | This study | N/A |
| FM5-p14 (*Reptilian orthoreovirus*)-mGFP | This study | N/A |
| FM5-p14 (*Reptilian orthoreovirus*)-mCherry | This study | N/A |
| FM5-p14 (*Reptilian orthoreovirus*)- miRFP670 | This study | N/A |
| FM5-Albumin signal sequence (SS)-8 amino acid GS Linker-B7 Transmembrane (TM)-mGFP | This study | N/A |
| FM5-Albumin SS-8 amino acid GS Linker-B7 TM-mCherry | This study | N/A |
| FM5-Albumin SS-8 amino acid GS Linker-B7 TM-miRFP670 | This study | N/A |
| FM5-ACE2-mGFP | This study | N/A |
| FM5-ACE2-mCherry | This study | N/A |
| FM5-ACE2-miRFP670 | This study | N/A |
| FM5-ACE2 Ectodomain-8 amino acid GS Linker-B7 TM/CTD-mGFP | This study | N/A |
| FM5-ACE2 Ectodomain-8 amino acid GS Linker-B7 TM-mCherry | This study | N/A |
| FM5-ACE2 Ectodomain-8 amino acid GS Linker-B7 TM-miRFP670 | This study | N/A |
| FM5-SARS-CoV-1 Spike-mGFP | This study | N/A |
| FM5-SARS-CoV-1 Spike-mCherry | This study | N/A |
| FM5-SARS-CoV-1 Spike-miRFP670 | This study | N/A |
| FM5-Spike Full-length (FL) (1-1273)-mGFP | This study | N/A |
| FM5-Spike Full-length (FL) (1-1273)-mCherry | This study | N/A |
| FM5-Spike Full-length (FL) (1-1273)-miRFP670 | This study | N/A |
| FM5-Spike SS (1-12)-RBD (319-541)-8 amino acid GS Linker-B7 TM/CTD-mGFP | This study | N/A |
| FM5-Spike SS (1-12)-RBD (319-541)-8 amino acid GS Linker-B7 TM/CTD-mCherry | This study | N/A |
| FM5-Spike SS (1-12)-RBD (319-541)-8 amino acid GS Linker-B7 TM/CTD-miRFP670 | This study | N/A |
| FM5-Spike SS (1-12)-RBD (319-541)-WT TM/CTD (1203-1273)-mGFP | This study | N/A |
| FM5-Spike SS (1-12)-RBD (319-541)-WT TM/CTD (1203-1273)-mCherry | This study | N/A |
| FM5-Spike SS (1-12)-RBD (319-541)-WT TM/CTD (1203-1273)-miRFP670 | This study | N/A |
| FM5-Spike Ectodomain (1-1213)-8 amino acid GS Linker-B7 TM/CTD-mGFP | This study | N/A |
| FM5-Spike Ectodomain (1-1213)-8 amino acid GS Linker-B7 TM/CTD-mCherry | This study | N/A |
| FM5-Spike Ectodomain (1-1213)-8 amino acid GS Linker-B7 TM/CTD-miRFP670 | This study | N/A |
| FM5-Spike B7 TM Swap with Extracellular Linker (1-1213-8 amino acid GS linker-B7 TM-1235-1273)-mGFP | This study | N/A |
| FM5-Spike B7 TM Swap with Extracellular Linker (1-1213-8 amino acid GS linker-B7 TM-1235-1273)-miRFP670 | This study | N/A |
| FM5-Spike B7 TM Swap (1-1213-B7 TM-1235-1273)-mGFP | This study | N/A |
| FM5-Spike B7 TM Swap (1-1213-B7 TM-1235-1273)-miRFP670 | This study | N/A |
| FM5-Spike ITGA1 TM Swap (1-1213-ITGA1 TM-1235-1273)-mGFP | This study | N/A |
| FM5-Spike ITGA1 TM Swap (1-1213-ITGA1 TM-1235-1273)-miRFP670 | This study | N/A |
| FM5-Spike 1-1268-mGFP | This study | N/A |
| FM5-Spike 1-1268-miRFP670 | This study | N/A |
| FM5-Spike 1-1266-mGFP | This study | N/A |
| FM5-Spike 1-1266-miRFP670 | This study | N/A |
| FM5-Spike 1-1263-mGFP | This study | N/A |
| FM5-Spike 1-1263-miRFP670 | This study | N/A |
| FM5-Spike 1-1260-mGFP | This study | N/A |
| FM5-Spike 1-1260-miRFP670 | This study | N/A |
| FM5-Spike 1-1256-mGFP | This study | N/A |
| FM5-Spike 1-1256-miRFP670 | This study | N/A |
| FM5-Spike 1-1245-mGFP | This study | N/A |
| FM5-Spike 1-1245-miRFP670 | This study | N/A |
| FM5-Spike 1-1239-mGFP | This study | N/A |
| FM5-Spike 1-1239-miRFP670 | This study | N/A |
| FM5-Spike 1-1234 (ΔCTD)-mGFP | This study | N/A |
| FM5-Spike 1-1234 (ΔCTD)-miRFP670 | This study | N/A |
| FM5-Spike CysAlaMutant (C1235A/C1236A/<br>C1240A/C1241A/C1243A/C1247A/C1248A/C1250A<br>/C1253A/C1254A)-mGFP | This study | N/A |
| FM5-Spike CysAlaMutant (C1235A/C1236A/<br>C1240A/C1241A/C1243A/C1247A/C1248A/C1250A<br>/C1253A/C1254A)-miRFP670 | This study | N/A |
| FM5-Spike C1235A/C1236A-mGFP | This study | N/A |
| FM5-Spike C1235A/C1236A-miRFP670 | This study | N/A |
| FM5-Spike SS (1-12)-ΔNTD (319-1273)-mGFP | This study | N/A |
| FM5-Spike SS (1-12)-ΔNTD (319-1273)-miRFP670 | This study | N/A |
| FM5-Spike ΔRBD (1-318, 540-1273)-mGFP | This study | N/A |
| FM5-Spike ΔRBD (1-318, 540-1273)-miRFP670 | This study | N/A |
| FM5-Spike ΔFusionPeptide (1-787, 807-1273)-<br>mGFP | This study | N/A |
| FM5-Spike ΔFusionPeptide (1-787, 807-1273)-<br>miRFP670 | This study | N/A |
| FM5-Spike ΔHR1 (1-919, 971-1273)-mGFP | This study | N/A |
| FM5-Spike ΔHR1 (1-919, 971-1273)-miRFP670 | This study | N/A |
| FM5-Spike ΔHR2 (1-1162, 1203-1273)-mGFP | This study | N/A |
| FM5-Spike ΔHR2 (1-1162, 1203-1273)-miRFP670 | This study | N/A |
| FM5-Spike ΔBothCleavageSites ("CS")<br>(Δ685/Δ686/Δ815/Δ816)-mGFP | This study | N/A |
| FM5-Spike ΔBothCS (Δ685/Δ686/Δ815/Δ816)-<br>miRFP670 | This study | N/A |
| FM5-Spike ΔS1/S2CS (Δ685/Δ686)-mGFP | This study | N/A |
| FM5-Spike ΔS1/S2CS (Δ685/Δ686)-miRFP670 | This study | N/A |
| FM5-Spike ΔS2'CS (Δ815/Δ816)-mGFP | This study | N/A |
| FM5-Spike ΔS2'CS (Δ815/Δ816)-miRFP670 | This study | N/A |
| FM5-Spike S813G-mGFP | This study | N/A |
| FM5-Spike S813G-miRFP670 | This study | N/A |
| FM5-Spike S816G-mGFP | This study | N/A |
| FM5-Spike S816G-miRFP670 | This study | N/A |
| FM5-Spike R815A-mGFP | This study | N/A |
| FM5-Spike R815A-miRFP670 | This study | N/A |
| FM5-Spike R815K-mGFP | This study | N/A |
| FM5-Spike R815K-miRFP670 | This study | N/A |
| FM5-Spike R815A/S816G-mGFP | This study | N/A |
| FM5-Spike R815A/S816G-miRFP670 | This study | N/A |
| FM5-Spike R815K/S816G-mGFP | This study | N/A |
| FM5-Spike R815K/S816G-miRFP670 | This study | N/A |
| FM5-Spike S813G/R815A-mGFP | This study | N/A |
| FM5-Spike S813G/R815A-miRFP670 | This study | N/A |
| FM5-Spike S813G/R815K-mGFP | This study | N/A |
| FM5-Spike S813G/R815K-miRFP670 | This study | N/A |
| FM5-Spike S1 (1-685)-WT TM/CTD (1203-1273)-<br>mGFP | This study | N/A |
| FM5-Spike S1 (1-685)-WT TM/CTD (1203-1273)-<br>miRFP670 | This study | N/A |
| FM5-Spike S1 (1-685)-8 amino acid GS Linker-B7 TM/CTD-mGFP | This study | N/A |
| FM5-Spike S1 (1-685)-8 amino acid GS Linker-B7 TM/CTD-miRFP670 | This study | N/A |
| FM5-Spike SS (1-12)-S2 (686-1273)-mGFP | This study | N/A |
| FM5-Spike SS (1-12)-S2 (686-1273)-miRFP670 | This study | N/A |
| FM5-Spike SS (1-12)-S2 (686-1213)-8 amino acid GS Linker-B7 TM/CTD Swap-mGFP | This study | N/A |
| FM5-Spike SS (1-12)-S2 (686-1213)-8 amino acid GS Linker B7 TM/CTD Swap-miRFP670 | This study | N/A |
| **Software and Algorithms** |  |  |
| ChemAxon Physicochemical Property Analysis | Morgan et al., *Angew. Chem. Int. Ed.*, 2018 | https://chemaxon.com/ |
| R Software Mann-Whitney U Statistical Analysis | Morgan et al., *Angew. Chem. Int. Ed.*, 2018 | https://www.r-project.org/ |
| NCGC Scaffold Hopper | NCATS Chemical Genomics Centre (NCGC) | https://tripod.nih.gov |
| RDKit Substructure Analysis | RDKit | https://www.rdkit.org/ |
| Python Software | Python | https://www.python.org/ |
| GraphPad Prism Nonlinear Regression | GraphPad | https://www.graphpad.com/ |
| MATLAB R2017b | Mathworks | https://www.mathworks.com/ |
| ImageJ | Schneider et al., *Nature Methods*, 2012 | https://imagej.nih.gov/ij/ |
| PANTHER v14.0 Gene Ontology | Mi et al., *Nature Protocols*, 2019 | https://pantherdb.org |
| BioRender (Paid Academic Plan) | BioRender | https://bio-render.com |
| Microsoft Excel | My computer | IT'S EXCEL-LENT! |

### RESOURCE AVAILABILITY

#### Lead Contact

Further information and requests for resources and reagents should be directed to and will be fulfilled by the lead contact, Clifford P. Brangwynne.

#### Materials Availability

Plasmids and cell lines generated in this study are available from the lead contact.

#### Data and Code Availability

Results from the drug repurposing screen are available in Table S1. Raw imaging data was not deposited online due to the size of the dataset (>1 TB), but can be obtained from the lead contact upon reasonable request.

### EXPERIMENTAL MODEL AND SUBJECT DETAILS

Please see **METHOD DETAILS** for information on cell lines and culture conditions. All cell lines were mycoplasma free at the beginning of the study.

### METHOD DETAILS

#### Plasmid construction

Lentiviral plasmids encoding fluorescently tagged proteins of interest were cloned as described in previous work (Sanders et al., 2020), which introduced monomeric fluorescent protein (GFP, mCherry, miRFP670) lentiviral vectors (FM5, ubiquitin C promoter) with standardized linkers/overlaps to allow Gibson assembly-based cloning in high \-throughput. With the exception of IDT-synthesized open-reading frames (p14, MYMK, MYMX), DNA fragments coding proteins of interest were amplified by PCR (oligonucleotides synthesized by IDT; see **KEY RESOURCES** table for origin of cDNA template), using Phusion® High-Fidelity DNA Polymerase (New England Biolabs or “NEB”). Gibson assembly (In-Fusion HD cloning kit, Takara) was used to insert gel-purified DNA (Qiagen, Gel Extraction Kit) into the desired lentiviral vector, linearized by NheI restriction enzyme (NEB) or AscI restriction enzyme (NEB) digestion. DNA was extracted from transformed Stellar competent bacteria (Takara) by mini-prep (Qiagen). DNA inserts were confirmed by Sanger sequencing (GENEWIZ), reading from both ends of the open reading frame.

To generate pLMN8-Gluc, the secreted version of Gluc gene was PCR amplified using Q5 polymerase (NEB) using gene specific primers purchased from IDT. The PCR amplicon was inserted into the SfiI-digested pLMN8 plasmid (Ploss et al., 2009) via In-Fusion cloning (Takara). The sequence of the resulting pLMN8 plasmid encoding secreted Gluc (pLMN8-Gluc) was confirmed by Sanger sequencing (Eton Bioscience). Mammalian cell, codon optimized pCAGGS-SARS-CoV-2 spike (S) was kindly provided by BEI resources (NIH). Retroviruses pseudotyped with vesicular stomatitis virus G protein (VSV-G) were used as positive controls. pLMN8-Gluc and pMLV gag-pol plasmids were amplified in NEB stable competent cells (NEB); pCAGGS-SARS-CoV-2 spike and pVSV-G, DH5α competent cells (ThermoFisher), then plasmids were purified by Nucleobond Xtra Midi (Takara).

#### Cell culture

A549, VeroE6, and Calu3 cells were obtained from the American Type Culture Collection (ATCC). 293T cells were a kind gift from Marc Diamond (UT Southwestern); U2OS cells, Tom Wuir (Princeton University); Beas2B cells, Celeste Nelson (Princeton University); A549 cells expressing human ACE2 and TMPRSS2 (A549-hACE2/TMPRSS2), Mohsan Saeed (Boston University). All cell lines were grown in Dulbecco’s Modified Eagle’s Medium (DMEM with high glucose and pyruvate, ThermoFisher) supplemented with 10% fetal bovine serum (FBS, Atlanta Biologicals) and 1% penicillin/streptomycin (P/S, Gibco), with the exception of: A549-ACE2/TMPRSS2, which were additionally maintained under puromycin (Sigma) and blasticidin (VWR) selection (both at 0.5 μg/mL); and Calu3 cells, which were grown in Eagle’s minimum essential media (EMEM, ATCC) with 10% FBS and 1% P/S. All cells were propagated at 37°C in a 5% CO_2_, 20% O_2_ environment.

#### Lentivirus production

Lentiviruses encoding fluorescently tagged proteins of interest were produced by using a previously optimized protocol (Sanders et al., 2014). Briefly, HEK293T cells were co-transfected with indicated FM5 construct and two helper plasmids (VSVG and PSP; driven cross-country from Marc Diamond lab, UT Southwestern) with Lipofectamine-3000 (Invitrogen; manufactured in Lithuania for some reason). Given the fact that is the year 2020 and transient transfection is a boondoggle of confounds, lipid-based transfection reagents were avoided with exception of virus production. Supernatant was collected 2-3 days post-transfection, cell debris was pelleted/discarded using centrifugation (1000xg), and lentivirus-containing media was used to infect indicated cell line in 96-well plates or stored at −80°C.

#### Generation of stable U2OS and VeroE6 cell lines

Lentivirus transduction was performed in 96-well plates as described (Sanders et al., 2020) with minor modifications. For large, inefficiently packaged spike constructs, 180 μL lentivirus supernatant was added to a single well of 96-well plate. For smaller constructs, 30 or 60 μL lentivirus supernatant was applied. Washed, trypsinized, and media-quenched cells were added directly to virus, so at ∼10-20% confluency upon adhesion to the dish surface. Cells were grown for three days in lentivirus to obtain confluence, which maximizes viral transduction efficiency and resulting protein expression. Stable lines were washed, trypsinized, and passaged to 12-well then 6-well dishes prior to long-term storage in liquid nitrogen (freezing media = 90% FBS, 10% DMSO). All stable lines were subsequently maintained and passaged in 6-well plates until needed for live-cell microscopy experiments. Typically, stable cell lines were passaged at least 3-times over 8+ days prior to analysis. This allows elimination of cells expressing lethal levels of the fusion protein of interest. In all experiments, at least 90% of cells in a given population expressed indicated protein (or proteins). With few exceptions (e.g. B7 TM-GFP cells), stable cells featured <5 μM protein of interest on plasma membrane (much less if averaged across entire cell), as estimated by fluorescence calibrations described in previous work (Sanders et al., 2020). The utilized protocol minimizes over-expression artifacts and is preferred to lipid-based transient transfection (e.g. Lipofectamine-3000), which can further disrupt plasma membrane integrity and induce stress responses/toxicity.

For all described fluorescent fusion proteins, proper localization to the cell’s plasma membrane was confirmed by live cell confocal microscopy. In the case of certain spike variants (e.g. Δ815/Δ816, “ΔS2’CS”), expression was unexpectedly low despite multiple rounds of lentivirus transduction, and tagged protein was confined to the endoplasmic reticulum. Such behavior was independent of fluorescent tag (e.g. mCherry, miRFP670, mGFP) and is assumed to indicate misfolding, which decreases the likelihood of effective post-translational processing by the secretory pathway. These constructs were discarded or not studied in detail, as direct contribution to a given phenotype would be impossible to ascertain. In the case of spike “ΔS2’CS” (Δ815/Δ816), additional variants were analyzed (e.g. S813G), binding to ACE2 was confirmed (transcellular synapses), and cleavage site essentiality for membrane fusion was assessed.

#### Live cell confocal microscopy

Stable cell lines were plated on fibronectin-coated, 96-well glass bottom dishes (Cellvis) and immediately imaged (in the case of experiments requiring observation of individual fusion events) or following 24-hours culture (e.g. comparison of relative fusion between spike variants). A Nikon A1 laser-scanning confocal microscope equipped with 60x oil immersion lens (numerical aperture of 1.4) was used to collect confocal images. A humidified incubator kept cells at 37°C and 5% CO_2_. Proteins tagged with EYFP, mGFP (“GFP”), mCherry, or miRFP670 (“iRFP”) were imaged with 488, 488, 560, and 640 nm laser lines, respectively, and settings were optimized to minimize photobleaching and to negate bleed-through between channels. With the exception of heterokaryon co-culture assays, all confocal microscopy was performed on living cells to eliminated fixation-associated artifacts in subcellular localization.

#### MBCD TMR-cholesterol labeling and TIRF imaging

Cells were incubated with TopFluor® TMR-cholesterol (Avanti Polar Lipids #810385) complexed with methyl-beta-cyclodextrin (MβCD) in a 1:10 ratio (chol: MβCD). Total Internal Reflection Fluorescence (TIRF) Microscopy Images were collected on Nikon A1R+STORM (Nikon Ti2 frame) equipped with 405 nm, 488 nm, 561 nm, and 640 nm laser sources (Nikon LUN-F), a Princeton Instruments ProEM EMCCD camera, and SR HP Apo TIRF 100x/1.49 oil lens (MRD01995). The N-STORM module was used in TIRF mode and the TIRF angle was adjusted manually.

#### Heterokaryon co-culture assay

U2OS cells expressing SARS-CoV-2 spike-iRFP and ACE2-iRFP with their respective nuclear markers HNRNPA1-EYFP and FUS-mCherry were grown in 10-cm cell culture dishes (ThermoFisher), trypsinized with 0.05% EDTA-trypsin (ThermoFisher), resuspended in DMEM (10% FBS, 1% P/S), and mixed in 1:1 ratio. 5.4×10^!^ cells were immediately seeded per well into a glass-bottomed 384-well plate (CellVis) to a total of 80 μL volume using a Multidrop Combi SMART liquid-handling dispenser (ThermoFisher). Unless indicated, cells were incubated at 37°C for 5-hours, fixed with 4% paraformaldehyde (Electron Microscopy Services 16% PFA stock solution from freshly opened glass ampules was added directly to media to minimize variability between wells) for 10-minutes, washed with DPBS (Gibco), and stained with Hoechst (200 ng/mL). For VeroE6 cells co-cultures, the above procedure was followed exactly, replacing the ACE2-expressing U2OS cells with VeroE6 cells expressing FUS-mCh nuclear markers. Unlike U2OS cells, VeroE6 monkey cells feature endogenous ACE2 expression and are readily infected with SARS-CoV-2 virus.

#### Targeted compound dose-response assays

For the targeted compound screen (Figure 3), compounds were purchased and dissolved in water, methanol or DMSO to achieve stock solutions at ∼2000-fold concentration commonly reported by literature. Serial dilutions (7-doses, 3-fold dilutions unless indicated) were prepared in 20 μL DMEM per well and 5.4×10^!^ cells of each cell type (40 μL volume per) were added to a final volume of 100 μL (0.5% DMSO). Heterokaryon co-culture assays were carried out as described above.

#### Unbiased drug repurposing screen

For drug repurposing screen of 5985 compound library (derived from seven different commercial small molecule libraries; see **KEY RESOURCES** table), the described ACE2-U2OS heterokaryon assay was carried out by adding co-culture to 384-well plates with compounds pre-dispensed. Specifically, 240 nL of compound (10 mM, in dissolved in DMSO) was added using ECHO 550 (Labcyte) liquid dispenser to generate a final compound concentration of 30 μM upon addition of 80 μL cell co-culture.

#### Dose-response validation of hits from drug repurposing screen

For 7-point, dose-response assay, appropriate volumes of compound as 10 mM DMSO solution were dispensed using ECHO 550 (Labcyte) liquid dispenser to generate final concentrations of 40, 20, 10, 5.0, 2.5, 1.25, 0.625 μM upon addition of 80 μL cell co-culture. Wells were back-filled as necessary to keep the total DMSO volume of 320 nL consistent for all wells, including negative control, so as to maintain 0.4% DMSO concentration. To validate the top-24 hits, compounds were purchased from independent suppliers, dissolved in DMSO at 10 mM stock concentrations, and dispensed in 7-point dilutions according to procedure above.

#### Automated fixed cell confocal imaging and data acquisition

Heterokaryon assay development, characterization and high-throughput screening was carried out on a Eclipse Ti2 inverted scanning confocal microscope (Nikon) equipped with an automated Water Immersion Dispenser (WID). Wells were characterized by 16 full field of view regions (211×211 μm) imaged with a 60x, 1.2-numerical aperture, water-immersion, Nikon objective with 512×512 resolution. Bi-directional scanning with Hoechst (405 excitation/425-475 emission filter; channel 1), GFP (488/500-550; channel 2), and mCherry (561/570-620; channel 3) channels were acquired by a [2]->[1,3] line series through a 50 µm pinhole at a rate of one image per second. An automated image acquisition protocol was developed in the Nikon NIS-Elements JOBS module to navigate within each well and over the 384-well plate. Automated image processing and all subsequent analyses were implemented in MATLAB R2017b. See **QUANTIFICATION AND STATISTICAL ANALYSIS** for details.

#### Fluorescence recovery after photobleaching (FRAP)

Stable U2OS cell lines expressing indicated GFP-labeled proteins of interest were cultured for 24-hours on a 96-well glass-bottom dish (CellVis) and imaged using a Nikon A1 laser-scanning confocal microscope as described. Photobleaching was performed by scanning a 488-nm laser over a circular region of interest ∼6.5 µm in diameter, while focusing on the plasma membrane of single cells, validated by carefully tuning the focus to a plane bellow the non-fluorescent nuclei until they were no longer observable (compare transmitted light and fluorescent images in Figure 6A) and fluorescence signal within the surrounding area reached its maximum. See **QUANTIFICATION AND STATISTICAL ANALYSIS** for analysis.

#### Protein partitioning measurements in giant plasma membrane vesicles (GPMVs)

Cell membranes were stained with 5 µg/ml of Texas Red DHPE or Annexin V 647 (ThermoFisher), respectively, red or far-red fluorescent lipid dyes that strongly partition to disordered phases (Baumgart et al., 2007; Klymchenko and Kreder, 2014; Stone et al., 2017). Following staining, GPMVs were isolated as described (Sezgin et al., 2012) from U2OS stable cells lines expressing the protein of interest (LAT results were obtained from transient co-transfections). Briefly, GPMV formation was induced by 2 mM N-ethylmaleimide (NEM) in hypotonic buffer containing 100 mM NaCl, 10 mM HEPES, and 2 mM CaCl_2_, pH 7.4. To quantify partitioning, GPMVs were observed on an inverted epifluorescence microscope (Nikon) at 4°C after treatment with 200 µM DCA to stabilize phase separation; this treatment has been previously demonstrated not to affect raft affinity of various proteins (Castello-Serrano et al., 2020). The partition coefficient (K_p,raft_) for each protein was calculated from fluorescence intensity of the construct in the raft and non-raft phase for >10 vesicles/trial, with multiple independent experiments (n=3) for each construct.

#### Generation of retroviral pseudoparticles

All pseudotyped retroviruses were generated by co-transfection of plasmids encoding (1) a provirus containing the Gaussia luciferase reporter gene (LMN8-Gluc), (2) mouse leukemia virus (MLV) gag-pol (Ploss et al., 2009), and (3) codon-optimized SARS-CoV-2 spike.

On the day prior to transfection, 1.4 x 10^7^ 293T cells were seeded in a 150-mm tissue culture dish. The following day, a total of 15 µg of total DNA was transfected using 90 µL X-tremeGENE HP DNA Transfection Regent (Roche). To generate luciferase reporter SARS CoV-2-Spp and VSV-Gpp controls, (1) pLMN8-Gluc, (2) MLV gag-pol, and (3) either SARS CoV-2 spike or VSV-G were co-transfected at a ratio of 4.5:4.5:1, giving rise to SARS-CoV-2pp and VSV-Gpp, respectively. No envelope pseudoparticles (NEpp) was also generated using (1) pLMN8-Gluc and (2) MLV gag-pol at a ratio of 1:1. Media was replaced after 6-18 hours with DMEM containing 3% FBS, nonessential amino acids (NEAA, 0.1 mM, ThermoFisher), HEPES (20 mM, ThermoFisher), polybrene (4 µg/mL, Sigma-Aldrich). Supernatants were harvested at 48- and 72-hours after transfection, pooled and filtered (0.45 μm pore size), aliquoted, and stored at −80°C until usage.

#### Pseudovirus blocking assay

Blocking assays with luciferase reporter pseudovirus were performed in poly-L-lysine coated flat-bottom 96-well plates using 1.5×10^4^ A549-ACE2-TMPRSS2 cells per well. The next day, all compounds (10 mM diluted in DMSO) except for MBCD were diluted to 50 µM by DMEM containing 3% FBS, NEAA (0.1 mM), HEPES (20 mM), polybrene (4 µg/ml) and penicillin-streptomycin. MBCD (40 mM diluted in PBS) was diluted to 2 mM by DMEM containing 0.5% DMSO, 3% FBS, NEAA (0.1 mM), HEPES (20 mM), polybrene (4 µg/ml), and penicillin-streptomycin. The final concentration of DMSO for all compounds was 0.5%. Two-fold serial dilutions of all compounds were co-cultured with cells for 2-hours at 37°C, and subsequently, the same volume of pseudovirus was added into the cells and incubated for 4-hours at 37°C. After incubation, wells were washed once with 100 μl Hank’s Buffered Saline Solution (HBSS, ThermoFisher), and the media changed to 100 μL DMEM containing 3% FBS, NEAA (0.1 mM), HEPES (20 mM), polybrene (4 µg/mL), and penicillin-streptomycin.

#### Pseudovirus luciferase assay

Luciferase assay were performed 48-hours after incubation. The supernatants were collected to assess Gaussia luciferase activity using Genecopoeia Luc-Pair Renilla luciferase HS Assay Kit (GeneCopoeia) following the manufacturer’s instruction and measured on a Tristar2 LB942 luminometer (Berthold Technologies).

#### SARS-CoV-2 isolate stock preparation and titration

All replication-competent SARS-CoV-2 experiments were performed in a biosafety level 3 laboratory (BSL-3) at the Boston University’ National Emerging Infectious Diseases Laboratories. Experiments were performed in a biosafety level 3 laboratory. 2019-nCoV/USA-WA1/2020 isolate (NCBI accession number: MN985325) of SARS-CoV-2 was obtained from the Centers for Disease Control and Prevention and BEI Resources. To generate the passage 1 (P1) virus stock, Vero E6 cells, pre-seeded the day before at a density of 10 million cells, were infected in T175 flasks with the master stock, diluted in 10ml final volume of Opti-MEM. Following virus adsorption to the cells at 37°C for 1h, 15ml DMEM containing 10% FBS and 1X penicillin/streptomycin was added to the flask. The next day, media was removed, cell were rinsed with 1X PBS and 25 ml of fresh DMEM containing 2% FBS was added. Two days later, when the cytopathic effect of the virus was clearly visible, culture medium was collected, filtered through a 0.2µ filter, and stored at −80°C. Our P2 working stock of the virus was prepared by infecting Vero E6 cells with the P1 stock, at a multiplicity of infection (MOI) of 0.1. Cell culture media was harvested at day two and day three post infection, and after the last harvest, ultracentrifuged (Beckman Coulter Optima L-100k; SW32 Ti rotor) for 2h at 25,000rpm over a 20% sucrose cushion. Following centrifugation, the media and sucrose were discarded and pellets were left to dry for 5 minutes at room temperature. Pellets were then resuspended over night at 4°C in 500ul of 1X PBS. The next day, concentrated virions were aliquoted at stored at −80°C.

The titer of our viral stock was determined by plaque assay. Vero E6 cells were seeded into a 12-well plate at a density of 2.5 x 10^5^ cells per well, and infected the next day with serial 10-fold dilutions of the virus stock for 1h at 37°C. Following virus adsorption, 1 ml of overlay media, consisting of 2X DMEM supplemented with 4% FBS and mixed at a 1:1 ratio with 1.2% Avicel (DuPont; RC-581), was added in each well. Three days later, the overlay medium was removed, the cell monolayer was washed with 1X PBS and fixed for 30 minutes at room temperature with 4% paraformaldehyde. Fixed cells were then washed with 1X PBS and stained for 1h at room temperature with 0.1% crystal violet prepared in 10% ethanol/water. After rinsing with tap water, the number of plaques were counted and the virus titer was calculated. The titer of our P2 virus stock was 4 x 10^8^ PFU/ml.

#### SARS-CoV-2 cholesterol depletion assay

The day prior infection, A549 expressing hACE2 and hTMPRSS2 cells were seeded at a density of 2×10^4^ per well in a poly-L-lysine coated flat-bottom 96-well plate. The next day, MBCD initial stock was prepared at a concentration of 20 mM in 1X PBS and 2-fold serial dilutions were then made using 1X PBS. Prior to infecting cells, a one-hour pretreatment of MBCD with SARS-CoV-2 virus or cells was carried out. Viral pretreatment was performed by mixing 25 ul of each MBCD dilution with 25 ul of SARS-CoV-2 (MOI of 0.5; 1×10^4^ PFU) per well then incubated for one-hour at 37°C. For cell pretreatment, media was removed from wells and cells were washed once with 1X PBS. Cells were then incubated for one-hour at 37°C with 25 ul of each MBCD dilution further diluted in 25ul of 1X PBS. Following pretreatments, media or PBS/MBCD mixes were removed from wells and wells were washed twice with 1X PBS. Untreated and MBCD-treated cells were then infected for one-hour at 37°C with 50ul of SARS-CoV-2 pretreated with MBCD, or with untreated SARS-CoV-2 (MOI of 0.5), respectively. One hour following virus adsorption, media was removed, cells were washed twice with PBS, and 200uL of DMEM containing 10% FBS and 1% penicillin-streptomycin was added in each well. Forty-eight hours post infection, cell culture media was harvested and stored at −80°C. Cells were washed twice with 1X PBS, and fixed with 200ul of 10% neutral buffered formalin for one hour at room temperature. Cells were then washed twice with 1X PBS, and taken out of the BSL-3 laboratory.

#### SARS-CoV-2 RT-qPCR

To determine SARS-CoV-2 RNA copies, total viral RNA was isolated from cell culture media using a Zymo Research Corporation Quick-RNA^TM^ Viral Kit (Zymo Research) according to manufacturer’s instructions. Viral RNA was quantified using single-step RT-quantitative real-time PCR (Quanta qScript One-Step RT-qPCR Kit; VWR) with primers and Taqman probes targeting the SARS-CoV-2 E gene as previously described (Corman et al., 2020). Briefly, a 20 µL reaction mixture containing 10 µL of Quanta qScript™ XLT One-Step RT-qPCR ToughMix, 0.5 mM Primer E_Sarbeco_F1 (ACAGGTACGTTAATAGTTAATAGCGT), 0.5 mM Primer E_Sarbeco_R2 (ATATTGCAGCAGTACGCA CACA), 0.25 mM Probe E_Sarbeco_P1 (FAM-ACACTAGCCATCCTTACTGCGCTTCG-BHQ1), and 2 µL of total RNA was subjected to RT-qPCR using Applied Biosystems QuantStudio 3 (ThermoFisher). The following cycling conditions were used: reverse transcription for 10-minutes at 55°C and denaturation at 94°C for 3-minutes followed by 45 cycles of denaturation at 94°C for 15-seconds and annealing/ extension at 58°C for 30-seconds. Ct values were determined using QuantStudio^TM^ Design and Analysis software V1.5.1 (ThermoFisher). For absolute quantification of viral RNA, a 389 bp fragment from the SARS-CoV-2 E gene was cloned onto pIDTBlue plasmid under an SP6 promoter using NEB PCR cloning kit (New England Biosciences). The cloned fragment was then *in vitro* transcribed (mMessage mMachine SP6 transcription kit; ThermoFisher) to generate a RT-qPCR standard.

#### SARS-CoV-2 immunofluorescence

Virus-infected cells were fixed in 4% paraformaldehyde for 30-minutes. The fixative was removed and the cell monolayer was washed twice with 1x PBS. The cells were permeabilized in 1x PBS + 0.1% Triton-X (PBT) for 15-minutes at room temperature and washed twice with 1x PBS. The cells were blocked in PBT + 10% goat serum (v/v) and 1% BSA (w/v) for 1-hour at room temperature before incubating overnight at 4°C with rabbit anti-SARS-CoV nucleocapsid antibody (1:2,000 dilution). The cells were then washed 5 times with 1x PBS and stained with Alexa568-conjugated goat anti-rabbit antibody (1:1000 dilution) in the dark at room temperature for 1-hour. The cells were washed 5 times with 1x PBS and counterstained with DAPI (1:1000). Images were acquired using the MuviCyte Live Cell Imaging System (PerkinElmer). Six images were captured per well with a 4x objective lens in an unbiased manner.

#### Human pathology

Human pathology studies were performed with the approval of the Institutional Review Board at Brigham and Women’s Hospital. Clinical autopsies with full anatomic dissection were performed on SARS-CoV-2 decedents by a board-certified anatomic pathologist (RFP) with appropriate infectious precautions. Lung samples were fixed in 10% neutral buffered formalin, embedded in paraffin, sectioned, and stained with hematoxylin and eosin using standard methods. Immunohistochemistry was performed on 4-µm-thick tissue sections following pressure cooker antigen retrieval (Target Retrieval Solution; pH 6.1; Agilent Dako) using a mouse monoclonal antibody directed against TTF-1 (clone 8G7G3/1; Agilent Dako) at 1:200 dilution. Control lung slides were obtained from the BWH Department of Pathology Autopsy Division archives. Glass slides were reviewed by a RFP using an Olympus BX41 microscope, and microscopic photographs were obtained with an Olympus DP27 camera and Olympus CellSens Entry software.

### QUANTIFICATION AND STATISTICAL ANALYSIS

#### Automated fixed cell image analysis

Automated image analyses were performed using MATLAB R2017b (MathWorks). The fraction of cells fused was measured by adaptive segmentation of a maximal intensity projection of the three image channels to delineate nuclei. For each nucleus, area and mean intensity of both nuclear markers in the GFP and mCherry channels was measured. Nuclei with area less than 10-μm^2^ after gaussian filtering and erosion were removed. Remaining nuclei with mean GFP or mCherry channel signal more than 50-digital levels above background were considered nuclear marker-positive. GFP/mCherry double-positive nuclei were designated as part of a syncytium. The fraction of cells fused was calculated per well as the ratio of syncytium-to nuclear marker-positive nuclei. Total nuclei per well was z-score-normalized to the plate negative control wells and considered viable when >- 3.

#### Fluorescence recovery after photobleaching (FRAP)

Analysis was performed in MATLAB R2017b (MathWorks) by selecting a circular ROI, ∼4 µm in diameter, co-centered with the photobleached spot as well as a reference unbleached region of similar size for photobleaching correction. FRAP traces were corrected for photobleaching and fitted to a standard exponential decay model of the form: *I*(*t*) = *A*(1 − *e*^−*τt*^), with the mobile fraction *A* and the decay time constant *τ* being free parameters. Recovery half-life *τ*_!/!_ was derived using the relation *τ*_1/2_ = *ln*(2)/*τ*.

#### Cheminformatics analysis

SMILES strings of all analyzed compound libraries were batch-processed in ChemAxon (version 20.8.2) by first correcting each compound to its major tautomeric and protonation state at a physiologically relevant pH of 7.4. A total of 20 physicochemical parameters were calculated for each compound. A Mann Whitney U test was conducted in R software (version 3.4.3, 2017) to assess statistically significant differences between libraries.

The compounds tested in the fusion screen were divided into two libraries: “non-hits”, containing all non-toxic molecules that passed quality control and with a z-score (fusion) > −3.0 (n=5551); and “hits”, containing non-toxic molecules that passed quality and had a z-score (fusion) < 3.0 (n=163). Both libraries were filtered for empty wells for which no SMILES codes were available (n=210 in non-hits and 1 in hits), yielding a final number of 5504 compounds of non-hits and 162 compounds in hits.

A list of GPCR inhibitors, which represent approximately 35% of FDA-approved drugs, was provided by previous work (Sriram and Insel, 2018). DrugBank (version 5.1.7) was downloaded and queried with the list to obtain SMILES codes, yielding the final GPCR library (n=459) for batch processing and analysis in ChemAxon as described above.

Box and whisker plots were plotted and linear regression analysis was conducted in GraphPad Prism (version 8.0.2) for MacOS.

#### Scaffold and substructure enrichment analyses

Initial scaffold enrichment analysis was conducted in NCGC Scaffold Hopper software (version 1.0). The batch-processed SMILES strings from ChemAxon were input for the hit library (n=163) to identify most common scaffolds present in this library. To assess enrichment relative to the starting library, the combined hit and non-hit libraries, i.e. all non-toxic molecules screened in the fusion assay that passed quality control and did not contain empty wells (n=5714), were input in the software as well. 34 and 981 scaffolds were identified in the hit and starting libraries, respectively.

To identify the compounds containing the enriched scaffolds, SMILES strings from the top-10 enriched scaffolds in the hit group were analyzed in RDKit (version 2020.02.5) substructure search module using Python programming language (version 3.6.12) and IPython (version 7.12.0). Enrichment was assessed by conducting a pooled population comparison of the frequency of a scaffold in the hit library and the frequency of that same scaffold in the starting library. These values were then used to calculate z-scores and two-tailed p-values. Compounds containing statistically significant scaffolds (p-value < 0.05) were visually inspected to assess if unique and more complex substructures exist. If identified, those SMILES codes were subjected to another round of substructure search followed by two-tailed p-value calculation as described.

#### Bioinformatics

We first acquired the complete set of viral proteins from viruses that infect humans yielding 1,391,780 proteins (data retrieved October 2020) (UniProt Consortium, 2015). Next, we filtered for proteins in which two or more transmembrane prediction tools predicted an overlapping transmembrane helix or a transmembrane helix has been experimentally verified, yielding 168,094 proteins (Käll et al., 2004; Sonnhammer et al., 1998). Of these proteins, we applied a sliding window approach to assess local density of cysteine residues around the transmembrane helices. Specifically, we scanned the thirty-residue regions that lie on the N- or C-terminal sides of each transmembrane helix, using a window size of 20. For each protein the transmembrane-adjacent window with the highest fraction of cysteine was taken as the protein’s cysteine fractional ‘score’. The complete set of protein scores is provided in Table S2. To summarize high-confidence hits we first removed redundancy by filtering for duplicate sequence entries that originated from strain-specific sequence deposition. This final set is provided as Table S2, with high density hits called out in Figure 5G.

In parallel, we acquired the complete set of human proteins (n = 20370) from Uniprot (data retrieved October 2020) (UniProt Consortium, 2015). We then similarly filtered for predicted transmembrane proteins, yielding 5182 candidates (Käll et al., 2004; Sonnhammer et al., 1998). Of these proteins, we applied the same sliding window approach as for viral proteins as described above. The complete set of protein scores is provided in Table S3. We further subjected these putatively cysteine-rich transmembrane proteins to manual filtering to identify “spike-like” human proteins, which feature cysteine motifs in cytosol and aromatics at the ectodomain-plasma membrane interface. Results are summarized in Figure 5H with gene ontology (PantherDB) presented in Figure S6D.

## References

Abrams, M.E., Johnson, K.A., Perelman, S.S., Zhang, L.-S., Endapally, S., Mar, K.B., Thompson, B.M., McDonald, J.G., Schoggins, J.W., Radhakrishnan, A., et al. (2020). Oxysterols provide innate immunity to bacterial infection by mobilizing cell surface accessible cholesterol. Nat Microbiol 5, 929– 942.

Bauer, L., Ferla, S., Head, S.A., Bhat, S., Pasunooti, K.K., Shi, W.Q., Albulescu, L., Liu, J.O., Brancale, A., van Kuppeveld, F.J.M., et al. (2018). Structure-activity relationship study of itraconazole, a broad-range inhibitor of picornavirus replication that targets oxysterol-binding protein (OSBP). Antiviral Res. 156, 55–63.

Baumgart, T., Hammond, A.T., Sengupta, P., Hess, S.T., Holowka, D.A., Baird, B.A., and Webb, W.W. (2007). Large-scale fluid/fluid phase separation of proteins and lipids in giant plasma membrane vesicles. Proc. Natl. Acad. Sci. U.S.a. 104, 3165–3170.

Belouzard, S., Millet, J.K., Licitra, B.N., and Whittaker, G.R. (2012). Mechanisms of coronavirus cell entry mediated by the viral spike protein. Viruses 4, 1011–1033.

Bi, P., Ramirez-Martinez, A., Li, H., Cannavino, J., McAnally, J.R., Shelton, J.M., Sánchez-Ortiz, E., Bassel-Duby, R., and Olson, E.N. (2017). Control of muscle formation by the fusogenic micropeptide myomixer. Science 356, 323–327.

Bouhaddou, M., Memon, D., Meyer, B., White, K.M., Rezelj, V.V., Correa Marrero, M., Polacco, B.J., Melnyk, J.E., Ulferts, S., Kaake, R.M., et al. (2020). The Global Phosphorylation Landscape of SARS-CoV-2 Infection. Cell 182, 685–712.e19.

Brüning, A., Rahmeh, M., and Friese, K. (2013). Nelfinavir and bortezomib inhibit mTOR activity via ATF4-mediated sestrin-2 regulation. Mol Oncol 7, 1012–1018.

Bryce, C., Grimes, Z., Pujadas, E., Ahuja, S., Beasley, M.B., Albrecht, R., Hernandez, T., Stock, A., Zhao, Z., Rasheed, Al, M., et al. (2020). Pathophysiology of SARS-CoV-2: targeting of endothelial cells renders a complex disease with thrombotic microangiopathy and aberrant immune response. The Mount Sinai COVID-19 autopsy experience. medRxiv 1–24.

Buchrieser, J., Dufloo, J., Hubert, M., Monel, B., Planas, D., Rajah, M.M., Planchais, C., Porrot, F., Guivel-Benhassine, F., Van der Werf, S., et al. (2020). Syncytia formation by SARS-CoV-2 infected cells. 395, 497–25.

Cai, X., Xu, Y., Cheung, A.K., Tomlinson, R.C., Alcázar-Román, A., Murphy, L., Billich, A., Zhang, B., Feng, Y., Klumpp, M., et al. (2013). PIKfyve, a class III PI kinase, is the target of the small molecular IL-12/IL-23 inhibitor apilimod and a player in Toll-like receptor signaling. Chem Biol 20, 912–921.

Caly, L., Druce, J.D., Catton, M.G., Jans, D.A., and Wagstaff, K.M. (2020). The FDA-approved drug ivermectin inhibits the replication of SARS-CoV-2 in vitro. Antiviral Res. 178, 104787.

Carbajo-Lozoya, J., Müller, M.A., Kallies, S., Thiel, V., Drosten, C., and Brunn von, A. (2012). Replication of human coronaviruses SARS-CoV, HCoV-NL63 and HCoV-229E is inhibited by the drug FK506. Virus Res. 165, 112–117.

Castello-Serrano, I., Lorent, J.H., Ippolito, R., Levental, K.R., and Levental, I. (2020). Myelin-Associated MAL and PLP Are Unusual among Multipass Transmembrane Proteins in Preferring Ordered Membrane Domains. J Phys Chem B 124, 5930–5939.

Cattin-Ortolá, J., Welch, L., Maslen, S.L., Skehel, J.M., Papa, G., James, L.C., and Munro, S. (2020). Sequences in the cytoplasmic tail of SARS-CoV-2 spike facilitate syncytia formation. 583, 830–830.

Chan, K.M.C., Arthur, A.L., Morstein, J., Jin, M., Bhat, A., Schlesinger, D., Son, S., Stevens, D.A., Drubin, D.G., and Fletcher, D.A. (2020a). Evolutionarily related small viral fusogens hijack distinct but modular actin nucleation pathways to drive cell-cell fusion. bioRxiv 6, 341–29.

Chan, K.M.C., Son, S., Schmid, E.M., and Fletcher, D.A. (2020b). A viral fusogen hijacks the actin cytoskeleton to drive cell-cell fusion. Elife 9, 3864.

Chen, A., Leikina, E., Melikov, K., Podbilewicz, B., Kozlov, M.M., and Chernomordik, L.V. (2008). Fusion-pore expansion during syncytium formation is restricted by an actin network. J. Cell. Sci. 121, 3619–3628.

Chen, C.Z., Xu, M., Pradhan, M., Gorshkov, K., Petersen, J., Straus, M.R., Zhu, W., Shinn, P., Guo, H., Shen, M., et al. (2020). Identifying SARS-CoV-2 entry inhibitors through drug repurposing screens of SARS-S and MERS-S pseudotyped particles. bioRxiv 33, 125.

Chlanda, P., Mekhedov, E., Waters, H., Sodt, A., Schwartz, C., Nair, V., Blank, P.S., and Zimmerberg, J. (2017). Palmitoylation Contributes to Membrane Curvature in Influenza A Virus Assembly and Hemagglutinin-Mediated Membrane Fusion. J. Virol. 91, 229.

Ciechonska, M., and Duncan, R. (2014). Reovirus FAST proteins: virus-encoded cellular fusogens. Trends Microbiol 22, 715–724.

Cohen, L.D., and Ziv, N.E. (2019). Neuronal and synaptic protein lifetimes. Curr Opin Neurobiol 57, 9–16.

Compton, A.A., and Schwartz, O. (2017). They Might Be Giants: Does Syncytium Formation Sink or Spread HIV Infection? PLoS Pathog. 13, e1006099.

Corman, V.M., Landt, O., Kaiser, M., Molenkamp, R., Meijer, A., Chu, D.K., Bleicker, T., Brünink, S., Schneider, J., Schmidt, M.L., et al. (2020). Detection of 2019 novel coronavirus (2019-nCoV) by real-time RT-PCR. Euro Surveill 25, 2431.

Cornell, C.E., McCarthy, N.L.C., Levental, K.R., Levental, I., Brooks, N.J., and Keller, S.L. (2017). n-Alcohol Length Governs Shift in Lo-Ld Mixing Temperatures in Synthetic and Cell-Derived Membranes. Biophys. J. 113, 1200–1211.

Corver, J., Broer, R., van Kasteren, P., and Spaan, W. (2009). Mutagenesis of the transmembrane domain of the SARS coronavirus spike glycoprotein: refinement of the requirements for SARS coronavirus cell entry. Virol. J. 6, 230–13.

Costeira, R., Lee, K.A., Murray, B., Christiansen, C., Castillo-Fernandez, J., Ni Lochlainn, M., Capdevila Pujol, J., Buchan, I., Kenny, L.C., Wolf, J., et al. (2020). Estrogen and COVID-19 symptoms: associations in women from the COVID Symptom Study. 1–16.

Daniels, L.B., Sitapati, A.M., Zhang, J., Zou, J., Bui, Q.M., Ren, J., Longhurst, C.A., Criqui, M.H., and Messer, K. (2020). Relation of Statin Use Prior to Admission to Severity and Recovery Among COVID-19 Inpatients. Am. J. Cardiol.

Das, A., Brown, M.S., Anderson, D.D., Goldstein, J.L., and Radhakrishnan, A. (2014). Three pools of plasma membrane cholesterol and their relation to cholesterol homeostasis. Elife 3, 19316.

Day, C.A., Kraft, L.J., Kang, M., and Kenworthy, A.K. (2012). Analysis of protein and lipid dynamics using confocal fluorescence recovery after photobleaching (FRAP). Curr Protoc Cytom Chapter 2, Unit2.19–2.19.29.

De Gassart, A., Bujisic, B., Zaffalon, L., Decosterd, L.A., Di Micco, A., Frera, G., Tallant, R., and Martinon, F. (2016). An inhibitor of HIV-1 protease modulates constitutive eIF2α dephosphorylation to trigger a specific integrated stress response. Proc. Natl. Acad. Sci. U.S.a. 113, E117–E126.

de Jesus, A.J., and Allen, T.W. (2013). The role of tryptophan side chains in membrane protein anchoring and hydrophobic mismatch. Biochim. Biophys. Acta 1828, 864–876.

Ding, Q., Heller, B., Capuccino, J.M.V., Song, B., Nimgaonkar, I., Hrebikova, G., Contreras, J.E., and Ploss, A. (2017). Hepatitis E virus ORF3 is a functional ion channel required for release of infectious particles. Proc. Natl. Acad. Sci. U.S.a. 114, 1147–1152.

Dittmar, M., Lee, J.S., Whig, K., Segrist, E., Li, M., Jurado, K., Samby, K., Ramage, H., Schultz, D., and Cherry, S. (2020). Drug repurposing screens reveal FDA approved drugs active against SARS-Cov-2. 26, 1266–33.

Duan, L., Zheng, Q., Zhang, H., Niu, Y., Lou, Y., and Wang, H. (2020). The SARS-CoV-2 Spike Glycoprotein Biosynthesis, Structure, Function, and Antigenicity: Implications for the Design of Spike-Based Vaccine Immunogens. Front Immunol 11, 576622.

Duan, R., Kim, J.H., Shilagardi, K., Schiffhauer, E.S., Lee, D.M., Son, S., Li, S., Thomas, C., Luo, T., Fletcher, D.A., et al. (2018). Spectrin is a mechanoresponsive protein shaping fusogenic synapse architecture during myoblast fusion. Nat. Cell Biol. 20, 688–698.

Duelli, D., and Lazebnik, Y. (2007). Cell-to-cell fusion as a link between viruses and cancer. Nat Rev Cancer 7, 968–976.

Dustin, M.L. (2014). The immunological synapse. Cancer Immunol Res 2, 1023–1033.

Epand, R.M., Sayer, B.G., and Epand, R.F. (2003). Peptide-induced formation of cholesterol-rich domains. Biochemistry 42, 14677–14689.

Feng, S., Sekine, S., Pessino, V., Li, H., Leonetti, M.D., and Huang, B. (2017). Improved split fluorescent proteins for endogenous protein labeling. Nat Commun 8, 370–11.

François, I.E.J.A., Bink, A., Vandercappellen, J., Ayscough, K.R., Toulmay, A., Schneiter, R., van Gyseghem, E., Van den Mooter, G., Borgers, M., Vandenbosch, D., et al. (2009). Membrane rafts are involved in intracellular miconazole accumulation in yeast cells. J. Biol. Chem. 284, 32680–32685.

Frankel, S.S., Wenig, B.M., Burke, A.P., Mannan, P., Thompson, L.D., Abbondanzo, S.L., Nelson, A.M., Pope, M., and Steinman, R.M. (1996). Replication of HIV-1 in dendritic cell-derived syncytia at the mucosal surface of the adenoid. Science 272, 115–117.

Giacca, M., Bussani, R., Schneider, E., Zentilin, L., Collesi, C., Ali, H., Braga, L., Secco, I., Volpe, M.C., Colliva, A., et al. (2020). Persistence of viral RNA, widespread thrombosis and abnormal cellular syncytia are hallmarks of COVID-19 lung pathology. medRxiv 1–22.

Goldstein, D.B. (1984). The effects of drugs on membrane fluidity. Annu Rev Pharmacol Toxicol 24, 43–64.

Goldstein, J.L., and Brown, M.S. (2015). A century of cholesterol and coronaries: from plaques to genes to statins. Cell 161, 161–172.

Gordon, D.E., Jang, G.M., Bouhaddou, M., Xu, J., Obernier, K., White, K.M., O’Meara, M.J., Rezelj, V.V., Guo, J.Z., Swaney, D.L., et al. (2020). A SARS-CoV-2 protein interaction map reveals targets for drug repurposing. Nature 583, 459–468.

Goronzy, I.N., Rawle, R.J., Boxer, S.G., and Kasson, P.M. (2018). Cholesterol enhances influenza binding avidity by controlling nanoscale receptor clustering. Chem Sci 9, 2340–2347.

Gouttenoire, J., Pollán, A., Abrami, L., Oechslin, N., Mauron, J., Matter, M., Oppliger, J., Szkolnicka, D., Dao Thi, V.L., van der Goot, F.G., et al. (2018). Palmitoylation mediates membrane association of hepatitis E virus ORF3 protein and is required for infectious particle secretion. PLoS Pathog. 14, e1007471.

Gray, E., Karslake, J., Machta, B.B., and Veatch, S.L. (2013). Liquid general anesthetics lower critical temperatures in plasma membrane vesicles. Biophys. J. 105, 2751–2759.

Haynes, B.F., Corey, L., Fernandes, P., Gilbert, P.B., Hotez, P.J., Rao, S., Santos, M.R., Schuitemaker, H., Watson, M., and Arvin, A. (2020). Prospects for a safe COVID-19 vaccine. Sci Transl Med 58, eabe0948.

He, C., Hu, X., Weston, T.A., Jung, R.S., Sandhu, J., Huang, S., Heizer, P., Kim, J., Ellison, R., Xu, J., et al. (2018). Macrophages release plasma membrane-derived particles rich in accessible cholesterol. Proc. Natl. Acad. Sci. U.S.a. 115, E8499–E8508.

Head, S.A., Shi, W.Q., Yang, E.J., Nacev, B.A., Hong, S.Y., Pasunooti, K.K., Li, R.-J., Shim, J.S., and Liu, J.O. (2017). Simultaneous Targeting of NPC1 and VDAC1 by Itraconazole Leads to Synergistic Inhibition of mTOR Signaling and Angiogenesis. ACS Chem Biol 12, 174–182.

Heald-Sargent, T., and Gallagher, T. (2012). Ready, set, fuse! The coronavirus spike protein and acquisition of fusion competence. Viruses 4, 557–580.

Higashi, T., Tokuda, S., Kitajiri, S.-I., Masuda, S., Nakamura, H., Oda, Y., and Furuse, M. (2013). Analysis of the “angulin” proteins LSR, ILDR1 and ILDR2--tricellulin recruitment, epithelial barrier function and implication in deafness pathogenesis. J. Cell. Sci. 126, 966–977.

Hirabayashi, S., Tajima, M., Yao, I., Nishimura, W., Mori, H., and Hata, Y. (2003). JAM4, a junctional cell adhesion molecule interacting with a tight junction protein, MAGI-1. Mol. Cell. Biol. 23, 4267– 4282.

Hoffmann, M., Kleine-Weber, H., and Pöhlmann, S. (2020a). A Multibasic Cleavage Site in the Spike Protein of SARS-CoV-2 Is Essential for Infection of Human Lung Cells. Mol. Cell 78, 779–784.e5.

Hoffmann, M., Kleine-Weber, H., Schroeder, S., Krüger, N., Herrler, T., Erichsen, S., Schiergens, T.S., Herrler, G., Wu, N.-H., Nitsche, A., et al. (2020b). SARS-CoV-2 Cell Entry Depends on ACE2 and TMPRSS2 and Is Blocked by a Clinically Proven Protease Inhibitor. Cell 181, 271–280.e278.

Hoffmann, M., Schroeder, S., Kleine-Weber, H., Müller, M.A., Drosten, C., and Pöhlmann, S. (2020c). Nafamostat Mesylate Blocks Activation of SARS-CoV-2: New Treatment Option for COVID-19. Antimicrob. Agents Chemother. 64, 269.

Holowka, D., and Baird, B. (1983). Structural studies on the membrane-bound immunoglobulin E-receptor complex. 1. Characterization of large plasma membrane vesicles from rat basophilic leukemia cells and insertion of amphipathic fluorescent probes. Biochemistry 22, 3466–3474.

Hu, B., Höfer, C.T., Thiele, C., and Veit, M. (2019a). Cholesterol Binding to the Transmembrane Region of a Group 2 Hemagglutinin (HA) of Influenza Virus Is Essential for Virus Replication, Affecting both Virus Assembly and HA Fusion Activity. J. Virol. 93, 531.

Hu, X., Weston, T.A., He, C., Jung, R.S., Heizer, P.J., Young, B.D., Tu, Y., Tontonoz, P., Wohlschlegel, J.A., Jiang, H., et al. (2019b). Release of cholesterol-rich particles from the macrophage plasma membrane during movement of filopodia and lamellipodia. Elife 8, 2453.

Iijima, M., Suzuki, M., Tanabe, A., Nishimura, A., and Yamada, M. (2006). Two motifs essential for nuclear import of the hnRNP A1 nucleocytoplasmic shuttling sequence M9 core. FEBS Lett 580, 1365–1370.

Im, Y.J., Raychaudhuri, S., Prinz, W.A., and Hurley, J.H. (2005). Structural mechanism for sterol sensing and transport by OSBP-related proteins. Nature 437, 154–158.

Johnson, J.E., Gonzales, R.A., Olson, S.J., Wright, P.F., and Graham, B.S. (2007). The histopathology of fatal untreated human respiratory syncytial virus infection. Mod Pathol 20, 108–119.

Kang, Y.-L., Chou, Y.-Y., Rothlauf, P.W., Liu, Z., Soh, T.K., Cureton, D., Case, J.B., Chen, R.E., Diamond, M.S., Whelan, S.P.J., et al. (2020). Inhibition of PIKfyve kinase prevents infection by Zaire ebolavirus and SARS-CoV-2. Proc. Natl. Acad. Sci. U.S.a. 117, 20803–20813.

Kawase, M., Shirato, K., van der Hoek, L., Taguchi, F., and Matsuyama, S. (2012). Simultaneous treatment of human bronchial epithelial cells with serine and cysteine protease inhibitors prevents severe acute respiratory syndrome coronavirus entry. J. Virol. 86, 6537–6545.

Käll, L., Krogh, A., and Sonnhammer, E.L.L. (2004). A combined transmembrane topology and signal peptide prediction method. J. Mol. Biol. 338, 1027–1036.

Ke, Z., Oton, J., Qu, K., Cortese, M., Zila, V., McKeane, L., Nakane, T., Zivanov, J., Neufeldt, C.J., Cerikan, B., et al. (2020). Structures and distributions of SARS-CoV-2 spike proteins on intact virions. Nature 1–7.

Kim, J.H., and Chen, E.H. (2019). The fusogenic synapse at a glance. J. Cell. Sci. 132, jcs213124.

Kindrachuk, J., Ork, B., Hart, B.J., Mazur, S., Holbrook, M.R., Frieman, M.B., Traynor, D., Johnson, R.F., Dyall, J., Kuhn, J.H., et al. (2015). Antiviral potential of ERK/MAPK and PI3K/AKT/mTOR signaling modulation for Middle East respiratory syndrome coronavirus infection as identified by temporal kinome analysis. Antimicrob. Agents Chemother. 59, 1088–1099.

Kinnebrew, M., Iverson, E.J., Patel, B.B., Pusapati, G.V., Kong, J.H., Johnson, K.A., Luchetti, G., Eckert, K.M., McDonald, J.G., Covey, D.F., et al. (2019). Cholesterol accessibility at the ciliary membrane controls hedgehog signaling. Elife 8, 86.

Kirby, B.J., Collier, A.C., Kharasch, E.D., Whittington, D., Thummel, K.E., and Unadkat, J.D. (2011). Complex drug interactions of HIV protease inhibitors 1: inactivation, induction, and inhibition of cytochrome P450 3A by ritonavir or nelfinavir. Drug Metab Dispos 39, 1070–1078.

Klein, S., Cortese, M., Winter, S.L., Wachsmuth-Melm, M., Neufeldt, C.J., Cerikan, B., Stanifer, M.L., Boulant, S., Bartenschlager, R., and Chlanda, P. (2020). SARS-CoV-2 structure and replication characterized by in situcryo-electron tomography. 27, 1–11.

Klymchenko, A.S., and Kreder, R. (2014). Fluorescent probes for lipid rafts: from model membranes to living cells. Chem Biol 21, 97–113.

Lan, J., Ge, J., Yu, J., Shan, S., Zhou, H., Fan, S., Zhang, Q., Shi, X., Wang, Q., Zhang, L., et al. (2020). Structure of the SARS-CoV-2 spike receptor-binding domain bound to the ACE2 receptor. Nature 581, 215–220.

Levental, I., Byfield, F.J., Chowdhury, P., Gai, F., Baumgart, T., and Janmey, P.A. (2009). Cholesterol-dependent phase separation in cell-derived giant plasma-membrane vesicles. Biochem. J. 424, 163–167.

Levental, I., Grzybek, M., and Simons, K. (2011). Raft domains of variable properties and compositions in plasma membrane vesicles. Proc. Natl. Acad. Sci. U.S.a. 108, 11411–11416.

Levental, I., Levental, K.R., and Heberle, F.A. (2020). Lipid Rafts: Controversies Resolved, Mysteries Remain. Trends Cell Biol. 30, 341–353.

Levental, I., Lingwood, D., Grzybek, M., Coskun, U., and Simons, K. (2010). Palmitoylation regulates raft affinity for the majority of integral raft proteins. Proc. Natl. Acad. Sci. U.S.a. 107, 22050–22054.

Levental, K.R., and Levental, I. (2015). Isolation of giant plasma membrane vesicles for evaluation of plasma membrane structure and protein partitioning. Methods Mol. Biol. 1232, 65–77.

Li, G.-M., Li, Y.-G., Yamate, M., Li, S.-M., and Ikuta, K. (2007). Lipid rafts play an important role in the early stage of severe acute respiratory syndrome-coronavirus life cycle. Microbes Infect. 9, 96–102.

Li, W., Moore, M.J., Vasilieva, N., Sui, J., Wong, S.K., Berne, M.A., Somasundaran, M., Sullivan, J.L., Luzuriaga, K., Greenough, T.C., et al. (2003). Angiotensin-converting enzyme 2 is a functional receptor for the SARS coronavirus. Nature 426, 450–454.

Liao, K.W., Chou, W.C., Lo, Y.C., and Roffler, S.R. (2001). Design of transgenes for efficient expression of active chimeric proteins on mammalian cells. Biotechnol. Bioeng. 73, 313–323.

Liao, Y., Yuan, Q., Torres, J., Tam, J.P., and Liu, D.X. (2006). Biochemical and functional characterization of the membrane association and membrane permeabilizing activity of the severe acute respiratory syndrome coronavirus envelope protein. Virology 349, 264–275.

Liao, Y., Zhang, S.M., Neo, T.L., and Tam, J.P. (2015). Tryptophan-dependent membrane interaction and heteromerization with the internal fusion peptide by the membrane proximal external region of SARS-CoV spike protein. Biochemistry 54, 1819–1830.

Lin, Y.-C., Chen, B.-M., Lu, W.-C., Su, C.-I., Prijovich, Z.M., Chung, W.-C., Wu, P.-Y., Chen, K.-C., Lee, I.-C., Juan, T.-Y., et al. (2013). The B7-1 cytoplasmic tail enhances intracellular transport and mammalian cell surface display of chimeric proteins in the absence of a linear ER export motif. PLoS ONE 8, e75084.

Linton, M.F., Tao, H., Linton, E.F., and Yancey, P.G. (2017). SR-BI: A Multifunctional Receptor in Cholesterol Homeostasis and Atherosclerosis. Trends Endocrinol Metab 28, 461–472.

Long, T., Qi, X., Hassan, A., Liang, Q., De Brabander, J.K., and Li, X. (2020). Structural basis for itraconazole-mediated NPC1 inhibition. Nat Commun 11, 152–11.

Lu, Y., Liu, D.X., and Tam, J.P. (2008a). Lipid rafts are involved in SARS-CoV entry into Vero E6 cells. Biochem. Biophys. Res. Commun. 369, 344–349.

Lu, Y., Neo, T.L., Liu, D.X., and Tam, J.P. (2008b). Importance of SARS-CoV spike protein Trp-rich region in viral infectivity. Biochem. Biophys. Res. Commun. 371, 356–360.

Martin, B.R. (2013). Chemical approaches for profiling dynamic palmitoylation. Biochem. Soc. Trans. 41, 43–49.

Mast, N., Zheng, W., Stout, C.D., and Pikuleva, I.A. (2013). Antifungal Azoles: Structural Insights into Undesired Tight Binding to Cholesterol-Metabolizing CYP46A1. Mol. Pharmacol. 84, 86–94.

Maudgal, P.C., and Missotten, L. (1978). Histopathology of human superficial herpes simplex keratitis. Br J Ophthalmol 62, 46–52.

McBride, C.E., and Machamer, C.E. (2010a). Palmitoylation of SARS-CoV S protein is necessary for partitioning into detergent-resistant membranes and cell-cell fusion but not interaction with M protein. Virology 405, 139–148.

McBride, C.E., and Machamer, C.E. (2010b). A single tyrosine in the severe acute respiratory syndrome coronavirus membrane protein cytoplasmic tail is important for efficient interaction with spike protein. J. Virol. 84, 1891–1901.

McBride, C.E., Li, J., and Machamer, C.E. (2007). The cytoplasmic tail of the severe acute respiratory syndrome coronavirus spike protein contains a novel endoplasmic reticulum retrieval signal that binds COPI and promotes interaction with membrane protein. J. Virol. 81, 2418–2428.

Meher, G., Bhattacharjya, S., and Chakraborty, H. (2019). Membrane Cholesterol Modulates Oligomeric Status and Peptide-Membrane Interaction of Severe Acute Respiratory Syndrome Coronavirus Fusion Peptide. J Phys Chem B 123, 10654–10662.

Menter, T., Haslbauer, J.D., Nienhold, R., Savic, S., Hopfer, H., Deigendesch, N., Frank, S., Turek, D., Willi, N., Pargger, H., et al. (2020). Postmortem examination of COVID-19 patients reveals diffuse alveolar damage with severe capillary congestion and variegated findings in lungs and other organs suggesting vascular dysfunction. Histopathology 77, 198–209.

Millet, J.K., and Whittaker, G.R. (2014). Host cell entry of Middle East respiratory syndrome coronavirus after two-step, furin-mediated activation of the spike protein. Proc. Natl. Acad. Sci. U.S.a. 111, 15214–15219.

Mittal, A., Manjunath, K., Ranjan, R.K., Kaushik, S., Kumar, S., and Verma, V. (2020). COVID-19 pandemic: Insights into structure, function, and hACE2 receptor recognition by SARS-CoV-2. PLoS Pathog. 16, e1008762.

Oda, Y., Sugawara, T., Fukata, Y., Izumi, Y., Otani, T., Higashi, T., Fukata, M., and Furuse, M. (2020). The extracellular domain of angulin-1 and palmitoylation of its cytoplasmic region are required for angulin-1 assembly at tricellular contacts. J. Biol. Chem. 295, 4289–4302.

Ou, X., Liu, Y., Lei, X., Li, P., Mi, D., Ren, L., Guo, L., Guo, R., Chen, T., Hu, J., et al. (2020). Characterization of spike glycoprotein of SARS-CoV-2 on virus entry and its immune cross-reactivity with SARS-CoV. Nat Commun 11, 1620–12.

Papa, G., Mallery, D.L., Albecka, A., Welch, L., Cattin-Ortolá, J., Luptak, J., Paul, D., McMahon, H.T., Goodfellow, I.G., Carter, A., et al. (2020). Furin cleavage of SARS-CoV-2 Spike promotes but is not essential for infection and cell-cell fusion. bioRxiv 1, e14–e24.

Pelkmans, L. (2005). Secrets of caveolae- and lipid raft-mediated endocytosis revealed by mammalian viruses. Biochim. Biophys. Acta 1746, 295–304.

Pelkmans, L., and Helenius, A. (2003). Insider information: what viruses tell us about endocytosis. Curr. Opin. Cell Biol. 15, 414–422.

Petit, C.M., Chouljenko, V.N., Iyer, A., Colgrove, R., Farzan, M., Knipe, D.M., and Kousoulas, K.G. (2007). Palmitoylation of the cysteine-rich endodomain of the SARS-coronavirus spike glycoprotein is important for spike-mediated cell fusion. Virology 360, 264–274.

Ploss, A., Evans, M.J., Gaysinskaya, V.A., Panis, M., You, H., de Jong, Y.P., and Rice, C.M. (2009). Human occludin is a hepatitis C virus entry factor required for infection of mouse cells. Nature 457, 882–886.

Rajter, J.C., Sherman, M.S., Fatteh, N., Vogel, F., Sacks, J., and Rajter, J.-J. (2020). Use of Ivermectin Is Associated With Lower Mortality in Hospitalized Patients With Coronavirus Disease 2019: The ICON Study. Chest.

Riva, L., Yuan, S., Yin, X., Martin-Sancho, L., Matsunaga, N., Pache, L., Burgstaller-Muehlbacher, S., De Jesus, P.D., Teriete, P., Hull, M.V., et al. (2020). Discovery of SARS-CoV-2 antiviral drugs through large-scale compound repurposing. Nature 1–11.

Rockx, B., Kuiken, T., Herfst, S., Bestebroer, T., Lamers, M.M., Oude Munnink, B.B., de Meulder, D., van Amerongen, G., van den Brand, J., Okba, N.M.A., et al. (2020). Comparative pathogenesis of COVID-19, MERS, and SARS in a nonhuman primate model. Science 368, 1012–1015.

Sanders, D.W., Kaufman, S.K., DeVos, S.L., Sharma, A.M., Mirbaha, H., Li, A., Barker, S.J., Foley, A.C., Thorpe, J.R., Serpell, L.C., et al. (2014). Distinct tau prion strains propagate in cells and mice and define different tauopathies. Neuron 82, 1271–1288.

Sanders, D.W., Kedersha, N., Lee, D.S.W., Strom, A.R., Drake, V., Riback, J.A., Bracha, D., Eeftens, J.M., Iwanicki, A., Wang, A., et al. (2020). Competing Protein-RNA Interaction Networks Control Multiphase Intracellular Organization. Cell 181, 306–324.e328.

Sengupta, P., Hammond, A., Holowka, D., and Baird, B. (2008). Structural determinants for partitioning of lipids and proteins between coexisting fluid phases in giant plasma membrane vesicles. Biochim. Biophys. Acta 1778, 20–32.

Sezgin, E., Kaiser, H.-J., Baumgart, T., Schwille, P., Simons, K., and Levental, I. (2012). Elucidating membrane structure and protein behavior using giant plasma membrane vesicles. Nat Protoc 7, 1042–1051.

Shang, J., Wan, Y., Luo, C., Ye, G., Geng, Q., Auerbach, A., and Li, F. (2020). Cell entry mechanisms of SARS-CoV-2. Proc. Natl. Acad. Sci. U.S.a. 117, 11727–11734.

Shi, J., Bi, P., Pei, J., Li, H., Grishin, N.V., Bassel-Duby, R., Chen, E.H., and Olson, E.N. (2017). Requirement of the fusogenic micropeptide myomixer for muscle formation in zebrafish. Proc. Natl. Acad. Sci. U.S.a. 114, 11950–11955.

Shilagardi, K., Li, S., Luo, F., Marikar, F., Duan, R., Jin, P., Kim, J.H., Murnen, K., and Chen, E.H. (2013). Actin-propelled invasive membrane protrusions promote fusogenic protein engagement during cell-cell fusion. Science 340, 359–363.

Shirato, K., Kawase, M., and Matsuyama, S. (2018). Wild-type human coronaviruses prefer cell-surface TMPRSS2 to endosomal cathepsins for cell entry. Virology 517, 9–15.

Simons, K., and Ikonen, E. (1997). Functional rafts in cell membranes. Nature 387, 569–572.

Sobocińska, J., Roszczenko-Jasińska, P., Ciesielska, A., and Kwiatkowska, K. (2017). Protein Palmitoylation and Its Role in Bacterial and Viral Infections. Front Immunol 8, 2003.

Sohet, F., Lin, C., Munji, R.N., Lee, S.Y., Ruderisch, N., Soung, A., Arnold, T.D., Derugin, N., Vexler, Z.S., Yen, F.T., et al. (2015). LSR/angulin-1 is a tricellular tight junction protein involved in blood-brain barrier formation. J. Cell Biol. 208, 703–711.

Sonnhammer, E.L., Heijne von, G., and Krogh, A. (1998). A hidden Markov model for predicting transmembrane helices in protein sequences. Proc Int Conf Intell Syst Mol Biol 6, 175–182.

Soumpasis, D.M. (1983). Theoretical analysis of fluorescence photobleaching recovery experiments. Biophys. J. 41, 95–97.

Sriram, K., and Insel, P.A. (2018). G Protein-Coupled Receptors as Targets for Approved Drugs: How Many Targets and How Many Drugs? Mol. Pharmacol. 93, 251–258.

Stone, M.B., Shelby, S.A., Núñez, M.F., Wisser, K., and Veatch, S.L. (2017). Protein sorting by lipid phase-like domains supports emergent signaling function in B lymphocyte plasma membranes. Elife 6, 212.

Stratton, C.W., Tang, Y.-W., and Lu, H. (2020). Pathogenesis-Directed Therapy of 2019 Novel Coronavirus Disease. J Med Virol jmv.26610.

Tarcsay, Á., and Keserű, G.M. (2013). Contributions of molecular properties to drug promiscuity. J Med Chem 56, 1789–1795.

Tay, M.Z., Poh, C.M., Rénia, L., MacAry, P.A., and Ng, L.F.P. (2020). The trinity of COVID-19: immunity, inflammation and intervention. Nat. Rev. Immunol. 20, 363–374.

Tenchov, B.G., MacDonald, R.C., and Siegel, D.P. (2006). Cubic phases in phosphatidylcholine-cholesterol mixtures: cholesterol as membrane “fusogen”. Biophys. J. 91, 2508–2516.

Thorp, E.B., and Gallagher, T.M. (2004). Requirements for CEACAMs and cholesterol during murine coronavirus cell entry. J. Virol. 78, 2682–2692.

Tian, S., Hu, W., Niu, L., Liu, H., Xu, H., and Xiao, S.-Y. (2020). Pulmonary Pathology of Early-Phase 2019 Novel Coronavirus (COVID-19) Pneumonia in Two Patients With Lung Cancer. Journal of Thoracic Oncology 15, 700–704.

Trinh, M.N., Lu, F., Li, X., Das, A., Liang, Q., De Brabander, J.K., Brown, M.S., and Goldstein, J.L. (2017). Triazoles inhibit cholesterol export from lysosomes by binding to NPC1. Proc. Natl. Acad. Sci. U.S.a. 114, 89–94.

Tsuchiya, H. (2015). Membrane Interactions of Phytochemicals as Their Molecular Mechanism Applicable to the Discovery of Drug Leads from Plants. Molecules 20, 18923–18966.

Tsuchiya, H., and Mizogami, M. (2013). Interaction of local anesthetics with biomembranes consisting of phospholipids and cholesterol: mechanistic and clinical implications for anesthetic and cardiotoxic effects. Anesthesiol Res Pract 2013, 297141–18.

UniProt Consortium (2015). UniProt: a hub for protein information. Nucleic Acids Res. 43, D204– D212.

V’kovski, P., Kratzel, A., Steiner, S., Stalder, H., and Thiel, V. (2020). Coronavirus biology and replication: implications for SARS-CoV-2. Nat Rev Microbiol 5, 536–16.

Veatch, S.L. (2007). Electro-formation and fluorescence microscopy of giant vesicles with coexisting liquid phases. Methods Mol. Biol. 398, 59–72.

Veatch, S.L., and Keller, S.L. (2003). Separation of liquid phases in giant vesicles of ternary mixtures of phospholipids and cholesterol. Biophys. J. 85, 3074–3083.

Veatch, S.L., and Keller, S.L. (2005). Seeing spots: complex phase behavior in simple membranes. Biochim. Biophys. Acta 1746, 172–185.

Veatch, S.L., Cicuta, P., Sengupta, P., Honerkamp-Smith, A., Holowka, D., and Baird, B. (2008). Critical fluctuations in plasma membrane vesicles. ACS Chem Biol 3, 287–293.

Walls, A.C., Park, Y.-J., Tortorici, M.A., Wall, A., McGuire, A.T., and Veesler, D. (2020). Structure, Function, and Antigenicity of the SARS-CoV-2 Spike Glycoprotein. Cell 181, 281–292.e286.

Wan, J., Roth, A.F., Bailey, A.O., and Davis, N.G. (2007). Palmitoylated proteins: purification and identification. Nat Protoc 2, 1573–1584.

Wang, S., Li, W., Hui, H., Tiwari, S.K., Zhang, Q., Croker, B.A., Rawlings, S., Smith, D., Carlin, A.F., and Rana, T.M. (2020). Cholesterol 25-Hydroxylase inhibits SARS-CoV-2 and coronaviruses by depleting membrane cholesterol. Embo J. e2020106057.

Wei, J., Alfajaro, M.M., Hanna, R.E., DeWeirdt, P.C., Strine, M.S., Lu-Culligan, W.J., Zhang, S.-M., Graziano, V.R., Schmitz, C.O., Chen, J.S., et al. (2020). Genome-wide CRISPR screen reveals host genes that regulate SARS-CoV-2 infection. bioRxiv 20, 533.

Wrapp, D., Wang, N., Corbett, K.S., Goldsmith, J.A., Hsieh, C.-L., Abiona, O., Graham, B.S., and McLellan, J.S. (2020). Cryo-EM structure of the 2019-nCoV spike in the prefusion conformation. Science 367, 1260–1263.

Xia, S., Liu, M., Wang, C., Xu, W., Lan, Q., Feng, S., Qi, F., Bao, L., Du, L., Liu, S., et al. (2020). Inhibition of SARS-CoV-2 (previously 2019-nCoV) infection by a highly potent pan-coronavirus fusion inhibitor targeting its spike protein that harbors a high capacity to mediate membrane fusion. Cell Res. 30, 343–355.

Xu, J., Dang, Y., Ren, Y.R., and Liu, J.O. (2010). Cholesterol trafficking is required for mTOR activation in endothelial cells. Proc. Natl. Acad. Sci. U.S.a. 107, 4764–4769.

Yamamoto, M., Matsuyama, S., Li, X., Takeda, M., Kawaguchi, Y., Inoue, J.-I., and Matsuda, Z. (2016). Identification of Nafamostat as a Potent Inhibitor of Middle East Respiratory Syndrome Coronavirus S Protein-Mediated Membrane Fusion Using the Split-Protein-Based Cell-Cell Fusion Assay. Antimicrob. Agents Chemother. 60, 6532–6539.

Yan, R., Zhang, Y., Li, Y., Xia, L., Guo, Y., and Zhou, Q. (2020). Structural basis for the recognition of SARS-CoV-2 by full-length human ACE2. Science 367, 1444–1448.

Yuan, S., Chan, C.C.-Y., Chik, K.K.-H., Tsang, J.O.-L., Liang, R., Cao, J., Tang, K., Cai, J.-P., Ye, Z.-W., Yin, F., et al. (2020). Broad-Spectrum Host-Based Antivirals Targeting the Interferon and Lipogenesis Pathways as Potential Treatment Options for the Pandemic Coronavirus Disease 2019 (COVID-19). Viruses 12, 628.

Zawada, K.E., Wrona, D., Rawle, R.J., and Kasson, P.M. (2016). Influenza viral membrane fusion is sensitive to sterol concentration but surprisingly robust to sterol chemical identity. Sci Rep 6, 29842– 10.

Zhang, J., Chung, T., and Oldenburg, K. (1999). A Simple Statistical Parameter for Use in Evaluation and Validation of High Throughput Screening Assays. J Biomol Screen 4, 67–73.

Zhang, X.-J., Qin, J.-J., Cheng, X., Shen, L., Zhao, Y.-C., Yuan, Y., Lei, F., Chen, M.-M., Yang, H., Bai, L., et al. (2020). In-Hospital Use of Statins Is Associated with a Reduced Risk of Mortality among Individuals with COVID-19. Cell Metab. 32, 176–187.e4.

Zhou, F., Yu, T., Du, R., Fan, G., Liu, Y., Liu, Z., Xiang, J., Wang, Y., Song, B., Gu, X., et al. (2020). Clinical course and risk factors for mortality of adult inpatients with COVID-19 in Wuhan, China: a retrospective cohort study. Lancet 395, 1054–1062.

Zhou, Y., Vedantham, P., Lu, K., Agudelo, J., Carrion, R., Nunneley, J.W., Barnard, D., Pöhlmann, S., McKerrow, J.H., Renslo, A.R., et al. (2015). Protease inhibitors targeting coronavirus and filovirus entry. Antiviral Res. 116, 76–84.

Zhu, N., Zhang, D., Wang, W., Li, X., Yang, B., Song, J., Zhao, X., Huang, B., Shi, W., Lu, R., et al. (2020a). A Novel Coronavirus from Patients with Pneumonia in China, 2019. N Engl J Med 382, 727– 733.

Zhu, Y., Feng, F., Hu, G., Wang, Y., Yu, Y., Zhu, Y., Xu, W., Cai, X., Sun, Z., Han, W., et al. (2020b). The S1/S2 boundary of SARS-CoV-2 spike protein modulates cell entry pathways and transmission. 34, 7253–49.

Zidovetzki, R., and Levitan, I. (2007). Use of cyclodextrins to manipulate plasma membrane cholesterol content: evidence, misconceptions and control strategies. Biochim. Biophys. Acta 1768, 1311–1324.

Zinszner, H., Sok, J., Immanuel, D., Yin, Y., and Ron, D. (1997). TLS (FUS) binds RNA in vivo and engages in nucleo-cytoplasmic shuttling. J. Cell. Sci. 110 ( *Pt 15*), 1741–1750.

Zu, S., Deng, Y.-Q., Zhou, C., Li, J., Li, L., Chen, Q., Li, X.-F., Zhao, H., Gold, S., He, J., et al. (2020). 25-Hydroxycholesterol is a potent SARS-CoV-2 inhibitor. Cell Res. 46, 446–3.

